# A role for prefrontal dopamine signaling in control of goal-directed actions

**DOI:** 10.1101/2025.10.10.681663

**Authors:** A. Petitbon, A. Contini, R. Walle, M-C. Allichon, L. Hardt, F. Ducrocq, P. Couty, A. Oummadi, M-F. Angelo, S. Delcasso, M. Cazala, M. Depret, Y. Harouna, A. Ilie, F. Acedo, J. Correa Vazquez, R. Ortole, R. Thebeaud, A. Blanc, M. Rousseaud, A. Andrianarivelo, G. Ferreira, J. Barik, P. Vanhoutte, P. Trifilieff

## Abstract

Impairments in behavioral flexibility commonly found across psychiatric disorders have often been attributed to a dysfunction of the medial prefrontal cortex (mPFC), notably in relation with altered dopamine transmission. However, how dopamine transmission shapes neuronal activity in the mPFC to allow adapting to a changing situation remains largely elusive. Here, we show that dopamine dynamics in the mPFC encode the value of instrumental actions, in particular during reversal learning. Such signal shapes the activity of mPFC dopaminoceptive neurons during reversal learning through the recruitment of heteromers formed by dopamine D1 and D2 receptors with NMDA receptors. In accordance, blockade of either D1/NMDA or D2/NMDA heteromers in the mPFC selectively impairs reversal of outcome identities but not learning and expression of initial action-outcome associations. Our findings provide mechanistic evidence for a central role of dopamine in the mPFC to allow updating goal-directed actions.

## INTRODUCTION

Prevailing theories of decision-making propose that actions are goal-directed when they are guided by desire for a specific outcome, which is underpinned by the representation that a particular action will lead to that outcome^1–3^. This requires the ability to flexibly adjust actions according to their consequences, a function that is impaired across a variety of mental disorders such as schizophrenia, affective disorders or drug addiction^4–6^. Executive control deficits in psychiatric diseases have been closely associated with altered dopaminergic transmission, particularly within the mesocorticolimbic pathway^7,8^. In accordance with its role in encoding reward prediction error signal, numerous evidence highlight a role for mesolimbic dopamine transmission in the ability to adapt to changes in stimulus- or action-outcome relationships^9–12^. However, although the medial prefrontal cortex (mPFC) is well-established as a critical hub for behavioral flexibility^13,14^, the contribution of mesocortical dopamine transmission in this process remains poorly understood^15,16^.

In the mPFC, dopamine largely acts through D1 and D2 receptors, that are expressed by mostly segregated subpopulations^17^. Evidence support a role of both mPFC D1 and D2 receptor-dependent transmissions in behavioral flexibility^18,19^, suggesting that mPFC dopamine participates in coordinating the activity of neuronal subpopulations to allow for flexible actions. As a neuromodulator, dopamine acts in concert with neurotransmitters such as glutamate, through crosstalk in intracellular signaling^20,21^. More recently, heteromerization - that corresponds to direct physical interaction – between dopamine and glutamate NMDA receptors has been identified as a main molecular mechanism that allows integration of dopamine and NMDA-mediated signals^22^. On the one hand, physical interaction of the D1 receptor with the GluN1 subunit of the NMDA receptor^23–25^ tends to act as detector of coincidence of dopamine and glutamate transmissions by facilitating the respective functions of each protomers^26,27^. On the other hand, interaction of the D2 receptor with the GluN2B subunit of NMDA receptors decreases NMDA currents^28,29^. Here, we hypothesized that D1/NMDA and D2/NMDA heteromers could be central molecular mechanisms for the integration of dopamine and glutamate signals by mPFC neurons to allow behavioral flexibility.

We provide evidence that dopamine dynamics in the mPFC signal action value during encoding of action-outcome relationships, in particular during updating of associations. In accordance, silencing mesocorticolimbic dopamine transmission impairs reversal learning of outcome identities, but not acquisition or expression of initial action-outcome associations. Similarly, activity of dopaminoceptive neurons of the mPFC correlates with learning of instrumental associations, but display a unique signature during reversal learning, with D1-expressing neurons maintaining their activity selectively for newly-reinforced actions, while D2-expressing neurons become virtually unresponsive over learning of the new associations. Strikingly, D1/NMDA and D2/NMDA heteromers of the mPFC are necessary for such patterns of activity associated with reversal learning, but not with initial action-outcome associations. In line with these results, blockade of either D1/NMDA or D2/NMDA heteromers in the mPFC spares acquisition and expression of initial action-outcome associations but impairs reversal of outcome identities. Our findings provide evidence for a central role of dopamine in the mPFC to allow adaptation to action-outcome relationships, which is mediated by D1/NMDA and D2/NMDA heteromers.

## RESULTS

### Inhibiting VTA dopaminergic neurons selectively impairs the ability to adapt to changes in outcome identities but spares acquisition and expression of initial action-outcome associations

In order to disentangle the implication of mesocorticolimbic dopamine transmission in discrete components of goal-directed behaviors in mice, we chemogenetically inhibited VTA dopamine neurons during either learning of specific action-outcome associations, flexible expression of these associations, or during the reversal of associations (Fig.1a-d,h,l). To do so, animals first learned associations between a lever and a specific food reward. The devaluation procedure consisted in decreasing the value of one of the outcomes through ad libitum access (prefeeding), followed by a choice phase where the two levers were presented, with no reward delivered. Proper devaluation translates in a preference for the lever associated to the non-devalued (i.e. valued) outcome. Efficacy of sensory-specific devaluation was then tested through consumption of each reward given ad libitum (postfeeding).

**Figure 1:**
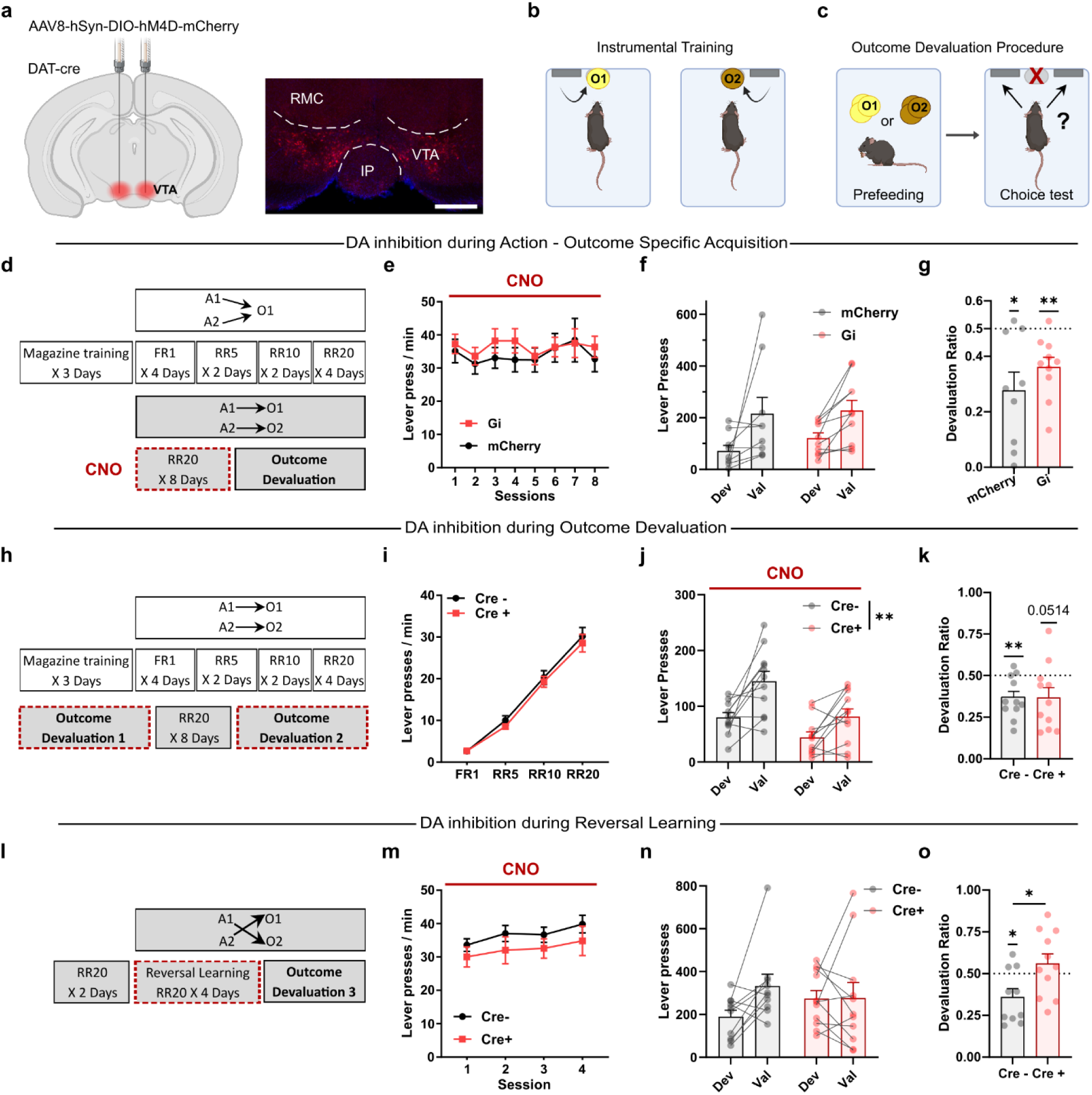
Chemogenetic inhibition of VTA dopaminergic neurons impairs the update of action-outcome contingencies but preserves learning and flexible expression of action-outcome associations. **a**, Schematic representation of the chemogenetic design together with a representative example of viral infection in the VTA (mCherry) of DAT-cre mice and nuclear staining with DAPI. Scale bar = 500 µm. **b**, Representation of the instrumental training paradigm. **c**, Representation of the outcome devaluation procedure. **d**, Detailed design of the action-outcome specific acquisition protocol with CNO administration during learning phase of specific action-outcome associations. **e**, Lever pressing rates during specific action – outcome acquisition (A1→O1 ; A2→O2) under CNO (2 mg/kg), DREADD Gi (n = 10), mCherry (n = 10). **f**-**g**, Extinction test during the outcome devaluation procedure, mCherry (n = 9), DREADD Gi (n = 10), (**f**) Number of lever presses on the devalued (Dev) or valued (Val) levers (Planned orthogonal contrast, within group effect F(1,17) = 11.094, *p* = 0.0040), (**g**) devaluation ratio 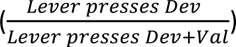 (One sample t-test against 0.5, mCherry *p* = 0.0100, Gi *p* = 0.0035). **h**, Detailed design of the learning protocols tested through outcome devaluation under CNO administration. **i**, Lever pressing rates during action – outcome training for Cre+ (n = 11) and Cre- (n = 11) groups. **j-k**, Extinction test during the outcome devaluation procedure under CNO (2 mg/kg), (**j**) Number of lever presses on each lever (Planned orthogonal contrast, between group effect F(1,20) = 11.541, *p* = 0.0029) (**k**) Devaluation ratio (One sample t-test against 0.5, Cre- *p* = 0.0054, Cre+ *p* = 0.0514). **l**, Representation of the reversal learning procedure under CNO and following outcome devaluation tests. **m**, Rates of lever pressing during the reversal of action – outcome associations under CNO (2 mg/kg), Cre+ (n = 11) and Cre- (n= 11). **n-o**, Extinction test during the outcome devaluation procedure, Cre+ (n = 10) and Cre- (n = 10), (**n**) number of lever presses on the Val and Dev levers (Planned orthogonal contrast, Devaluation effect F(1,19) = 2.767, *p* = 0.1126, and (**o**) devaluation ratio (One sample t-test against 0.5, Cre- *p* = 0.0217, Cre+ *p* = 0.3316 and two-tailed unpaired t-test, *p* = 0.0191). All datasets represent mean values +/- s.e.m. \**p*<0.05, \*\**p*<0.01. A, action; CNO, clozapine-N-oxide; Dev, devalued; FR, fixed ratio; IP, interpeduncular nucleus; O, outcome; RMC, magnocellular red nucleus; RR, random ratio; Val, valued; VTA, ventral tegmental area. Detailed statistics are shown in Supplementary Table 1

To assess the implication of VTA dopamine neurons in learning of specific action-outcome associations, animals were initially trained to press on two separate levers to obtain one reward (A1&A2→O1, Supplementary Fig.1b). Following acquisition, another reward was associated to one of the levers and mice were trained with this new protocol (A1→O1; A2→O2) for 8 sessions, and treated with the DREADD ligand CNO before each session (Fig.1d). This had no effect on instrumental performance (Fig.1e), nor did it impact the ability of animals to express these associations in an outcome-specific manner in the subsequent outcome devaluation procedure in the absence of CNO (Fig.1f-g). The ability to discriminate between outcome values was also similar between groups as evaluated in the postfeeding choice test (Supplementary Fig.1a,d).

We next assessed the implication of VTA dopaminergic projections for the expression of outcome devaluation, i.e. the ability of the animal to adapt actions in face of a change in outcome value, in an independent batch of mice (Fig.1h). There was no difference between groups in initial training (A1→O1 ; A2→O2, Fig.1i) in the absence of CNO administration. In the following outcome devaluation test under CNO administration, chemogenetic inhibition of VTA projections lowered overall lever pressing but did not alter the preference for the non-devalued lever (ie, valued) (Fig.1j-k). Of note, both groups displayed similar ability to differentiate both outcomes in the subsequent postfeeding test, although the Cre+ consumed overall more food than the Cre-group (Supplementary Fig.1f-g).

Last, we tested the implication of VTA dopaminergic neurons in reversal learning, i.e. the adaptation to an inversion of associations between lever pressings and outcomes (A1→O2, A2→O1). Animals were treated with CNO before each session of reversal (Fig.1l). Chemogenetic inhibition of VTA dopaminergic neurons did not impact performance over training sessions (Fig.1m), but impaired discrimination between the valued and devalued levers in a subsequent outcome devaluation test, in the absence of CNO administration (Fig.1n-o). Such a deficit is likely due to an impairment in updating action-outcome associations and not to an altered ability to discriminate between outcomes since both groups displayed preference for the non-devalued reward when freely available in the subsequent postfeeding test (Supplementary Fig.1i). Over the three outcome devaluation protocols, the amount of food consumed during the prefeeding session was similar between groups (Supplementary Fig.1c,e,h).

Altogether, these findings demonstrate that VTA dopaminergic neurons play a major role in adapting to changes in outcome identities, but are not required neither for the acquisition of A-O associations, nor for their flexible expression in face of a change in outcome value.

### Temporal sharpness of dopamine dynamics in the dmPFC correlates with action value

In order to characterize the processes underlying such a role of dopamine transmission, we next measured dopamine dynamics in relation with distinct phases of goal-directed behavior, i.e. associative learning, action-outcome association and reversal learning. We focused on the dorsomedial part of the prefontal cortex (dmPFC) in light of its central role in the modulation of goal-directed behavior^15,30,31^.

We recorded in vivo dopamine dynamics in the dmPFC through expression of the dopamine sensor GRAB-DA2m coupled with fiber photometry (Fig.2a). Animals were first subjected to Pavlovian conditioning (Fig.2b), where the presentation of a conditioned stimulus (CS, sound + LED) lasting 10 seconds led to the delivery of a reward (unconditioned stimulus, US) 3 seconds after the onset of the CS (Fig.2b-c, Supplementary Fig.3a). The occurrence of the CS did not lead to changes in dopamine signal neither in early nor late stage of training (Fig.2d-e, Supplementary Fig.2a). However, dopamine dynamics significantly increased just before, and peaked at licking onset during early pavlovian training (Fig.2f-g, Supplementary Fig.2b), but decreased with training (Fig.2g (left and right)). These data are in line with recent findings showing that the small proportion of reward-responding dopamine axons in the dmPFC are not involved in reward prediction, and that their activity in relation with rewarding stimuli decreases over training in a similar pavlovian conditioning task^32^.

We then assessed dopamine dynamics in relation with the encoding of an action–outcome association. Two levers were introduced, one triggering the CS-US schedule (reinforced lever, RL) as in the Pavlovian conditioning, and another with no outcome (unreinforced lever, UL) (Fig.2b-c, Supplementary Fig.3b-c). Comparison of dopamine responses between RL and UL across all sessions of instrumental training revealed an increase in dopamine dynamics following lever pressing (LP) on either RL or UL (Fig.2h-i, Supplementary Fig.2d). There was neither significant difference for AUC nor in peak z-score between RL and UL for the pre-LP or post-LP periods. However, there was a significant inhibition (below baseline) of the response following the rise in activity only for the RL (Fig.2i, Supplementary Fig.2d). Such a response is unlikely to be simply related to the occurrence of the CS following pressing on the RL, as no such dopamine response was present in Pavlovian conditioning (Fig.2d-f). Instead, this activity pattern might be related to learning of the association between RL and the reward, i.e. linked to the encoding of action value.

To further test this hypothesis, we examined the evolution of dopamine dynamics across learning sessions for the RL (Fig.2j-k, Supplementary Fig.2c). Of note, we could not conduct such an analysis for the UL due to the very low number of epochs, especially for late sessions. While the amplitude of the response following lever pressing was not different between early and late sessions, the subsequent decrease in dynamics was significant only for late sessions (Fig.2k), reinforcing the idea that such an inhibition of dopamine signal is related to learning of the lever pressing-reward association. Similarly, the increase in dopamine dynamics before lever pressing, i.e. during action engagement, in early sessions was significantly decreased with training (Fig.2k). Altogether, these findings suggest that, more than the lever pressing-induced increase of dopamine dynamics, learning of action-outcome association is accompanied by a temporal sharpening of the dopamine response during action exertion.

If such a dynamic is relevant for the coding of action value, one would expect the development of a similar pattern with learning of a new association, while it should be blunted when an action is not anymore associated with reward occurrence. We then asked how an inversion of action-outcome association – which implies the repression of previously learned associations while forming new ones^33,34^ - translates into changes in dopamine dynamics in the dmPFC. Animals were therefore trained in reversal learning sessions in which RL and UL were inverted (Fig.2b-c, Supplementary Fig.3d-e). When considering all sessions of reversal learning, we found that dopamine signal associated with the newly-reinforced lever was very similar to late training of operant learning, i.e. a significant decrease of the signal during action engagement and a sharp increase after lever pressing followed by a significant inhibition right before reward obtention (Fig.2l-m). Such a pattern was quite distinct to the one related to the newly unreinforced lever press that translated in a significant, though lower, increase of dopamine dynamics after lever pressing (Fig.2l-m, Supplementary Fig.2g). This dissociation between RL and UL was apparent early in reversal training (Fig.2n-o, Supplementary Fig.2e) and correlated with lever preference (Fig.2s) which was not the case during early sessions of initial instrumental training (Fig.2r). As for initial training, late sessions of reversal learning were accompanied with temporal sharpening of dopamine response related to the newly-reinforced lever, with strong increase in activity at the RL-induced CS that was preceded and followed by significant inhibition (Fig.2p-q, Supplementary Fig.2f). Lever pressing on the UL still translated in dopamine response, although much less synchronized compared to RL (Fig. 2p-q), which was due to highly temporally dispersed signal between animals (Supplementary Fig.2f). Of note, significant dopamine release was still observed at reward consumption (Supplementary Fig.2h). Our data suggest that dopamine dynamics in the dmPFC encode action value, with a temporally sharper response developing over learning of action-outcome association. Relevance of such signal better illustrates in reversal learning, in which a clear discrimination builds faster between a newly-reinforced action and a previously-reinforced one.

**Figure 2:**
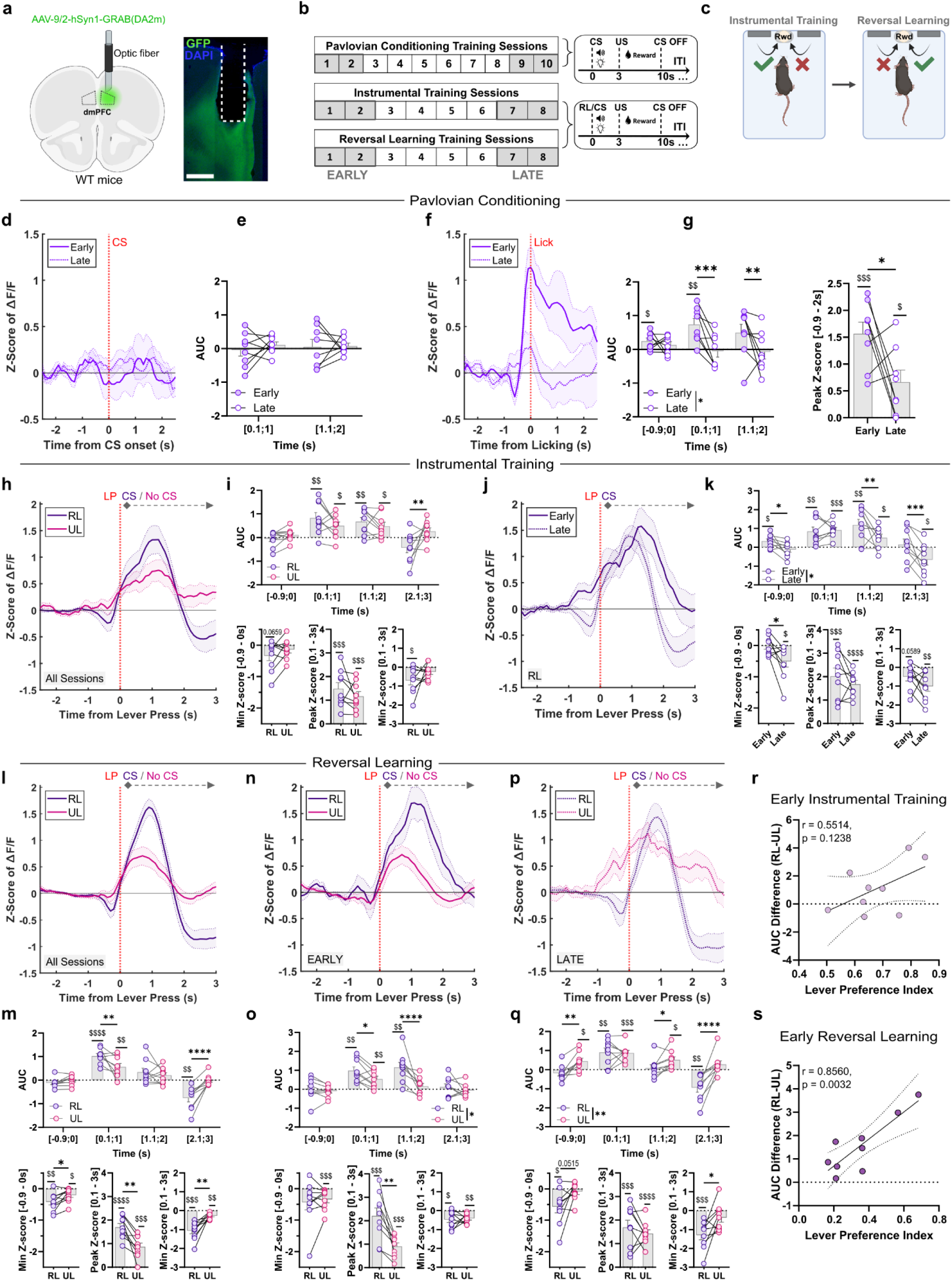
dmPFC dopamine encodes action value particularly during reversal of action-outcome associations. **a**, (**left**) Schematic representation of the viral vector strategy to record dopamine dynamics in the dmPFC, (**right**) histological example of the expression of GRAB-DA2m (GFP), scale bar = 500 µm. **b**, Experimental design. The early stage corresponds to the first 2 sessions and late stage to the last 2 sessions for pavlovian conditioning, instrumental training and reversal learning. **c**, (**left**) representation of a trial sequence where a CS (light + sound lasting 10 seconds) is followed by a US (reward delivery) 3 seconds after CS onset, (**right**) Schematic representation of instrumental training followed by reversal learning. **d-g**, Pavlovian conditioning during Early vs. Late sessions, n = 8 (**d**) Z-Score of ΔF/F at CS and (**e**) corresponding Area Under the Curve. (**f**) Z-Score of ΔF/F at licking and (**g**) (**left**) corresponding Area Under the Curve (Two-Factor RM ANOVA, Training Effect F (1, 7) = 7.976, *p* = 0.0256, Time x Training, F (2, 14) = 4.127 *p* = 0.0390, Tukey’s multiple comparisons test, [0.1s;1s] Early vs. Late *p* = 0.0004, [1.1s;2s] Early vs. Late *p* = 0.0047 ; One sample t-test against 0, [-0.9 s ; 0 s] Early *p* = 0.0283, [0.1 s ; 1 s] Early, *p* = 0.0050) (**right**) and Peak Z-Score over [-0.9-2s] (Paired t-test *p* = 0.0493 ; One sample t-test against 0, Early *p* = 0.0002, Late *p* = 0.0201). **h-k**, Instrumental training, n = 9. **h-i**, Reinforced vs. Unreinforced lever over 8 sessions (**h**) Z-Score of ΔF/F at lever pressing (**i**) (**top**) Area Under the Curve (Two-Factor RM ANOVA, Time x Lever F (3, 24) = 4.966 *p* = 0.0080, Tukey’s multiple comparisons test [2.1s;3s] RL vs. UL *p* = 0.0025 ; One sample t-test against 0, [0.1 s ; 1 s] RL *p* = 0.0064, [0.1 s ; 1 s] UL *p* = 0.0131, [1.1 s ; 2 s] RL *p* = 0.0051, [1.1 s ; 2 s] UL *p* = 0.0191) (**bottom left**) Minimum Z-Score over [-0.9-0s] (One sample t-test against 0, RL *p* = 0.0659) (**bottom center**) Peak Z-Score over [0.1-3s] (One sample t-test against 0, RL *p* = 0.0005, UL *p* = 0.0003) (**bottom right**) Minimum Z-Score over [0.1-3s] (One sample t-test against 0, RL *p* = 0.0139). **j-k**, Reinforced lever during early vs. late sessions (**j**) Z-Score of ΔF/F at lever pressing (**k**), (**top**) Area Under the Curve (Two-Factor RM ANOVA, Training Effect F (1, 8) = 5.519 *p* = 0.0467, Time x Training F (3, 24) = 3.938 *p* = 0.0204, Tukey’s multiple comparisons test, [-0.9s;0s] Early vs. Late *p* = 0.0297, [1.1s;2s] Early vs. Late *p* = 0.0018, [2.1s;3s] Early vs. Late *p* = 0.0004 ; One sample t-test against 0, [-0.9 s ; 0 s] Early *p* = 0.0238, [0.1 s ; 1 s] Early *p* = 0.0052, [0.1 s ; 1 s] Late *p* = 0.0001, [1.1 s ; 2 s] Early *p* = 0.0039, [1.1 s ; 2 s] Late *p* = 0.0347, [2.1 s ; 3 s] Late *p* = 0.0426) (**bottom left**) Minimum Z-Score over [-0.9-0s] (Paired t-test *p* = 0.0391, One sample t-test against 0, UL *p* = 0.0232) (**bottom center**) Peak Z-Score over [0.1-3s] (One sample t-test against 0, Early *p* = 0.0003, Late *p*<0.0001) (**bottom right**) Minimum Z-Score over [0.1-3s] (One sample t-test against 0, Early *p* = 0.0589, Late *p* = 0.0048). **l-m**, Reinforced vs. Unreinforced lever over 8 sessions, n = 9 (**l**) Z-Score of ΔF/F at lever pressing (**m**), (**top**) Area Under the Curve (Two-Factor RM ANOVA, Time x Lever F (3, 24) = 16.20 *p*<0.0001, Tukey’s multiple comparisons test [0.1s;1s] RL vs. UL *p* = 0.0015, [2.1s;3s] RL vs. UL *p*<0.0001 ; One sample t-test against 0, [0.1 s ; 1 s] RL p<0.0001, [0.1 s ; 1 s] UL *p* = 0.0046, [2.1 s ; 3 s] RL *p* = 0.0014) (**bottom left**) Minimum Z-Score over [-0.9-0s] (Paired t-test *p* = 0.0353 ; One sample t-test against 0, RL *p* = 0.0026, UL *p* = 0.0228) (**bottom center**) Peak Z-Score over [0.1-3s] (Paired t-test *p* = 0.0021 ; One sample t-test against 0, RL *p*<0.0001, UL *p* = 0.0006) (**bottom right**) Minimum Z-Score over [0.1-3s] (Paired t-test *p* = 0.0025 ; One sample t-test against 0, RL *p* = 0.0002, UL *p* = 0.0013). **n-o**, Reinforced vs. Unreinforced lever during early sessions, n = 9 (**n**) Z-Score of ΔF/F at lever pressing (**o**), (**top**) Area Under the Curve (Two-Factor RM ANOVA, Lever Effect F (1, 8) = 9.697 *p* = 0.0144, Time x Lever F (3, 24) = 6.118 *p* = 0.0030, Tukey’s multiple comparisons test [0.1s;1s] RL vs. UL *p* = 0.0126, [1.1s;2s] RL vs. UL *p*<0.0001, One sample t-test against 0, [0.1 s ; 1 s] RL *p* = 0.0011, [0.1 s ; 1 s] UL *p* = 0.0076, [1.1 s ; 2 s] RL *p* = 0.0021) (**bottom left**) Minimum Z-Score over [-0.9-0s] (One sample t-test against 0, UL *p* = 0.0142) (**bottom center**) Peak Z-Score over [0.1-3s] (Paired t-test *p* = 0.0025 ; One sample t-test against 0, RL *p* = 0.0002, UL *p* = 0.0004) (**bottom right**) Minimum Z-Score over [0.1-3s] (One sample t-test against 0, RL *p* = 0.0126, UL *p* = 0.0011). **p-q**, Reinforced vs. Unreinforced lever during late sessions, n = 9 (**p**) Z-Score of ΔF/F at lever pressing (**q**), (**top**) Area Under the Curve (Two-Factor RM ANOVA, Time x Lever F (3, 24) = 8.455 *p* = 0.0005, Tukey’s multiple comparisons test, [-0.9s;0s] RL vs. UL *p* = 0.0016, [1.1s;2s] RL vs. UL *p* = 0.0257, [2.1s;3s] RL vs. UL *p*<0.0001 ; One sample t-test against 0, [-0.9 s ; 0 s] UL *p* = 0.0230, [0.1 s ; 1 s] RL *p* = 0.0038, [0.1 s ; 1 s] UL *p* = 0.0003, [1.1 s ; 2 s] UL *p* = 0.0135, [2.1 s ; 3 s] RL *p* = 0.0038) (**bottom left**) Minimum Z-Score over [-0.9-0s] (Paired t-test *p* = 0.0515 ; One sample t-test against 0, RL *p* = 0.0195) (**bottom center**) Peak Z-Score over [0.1-3s] (One sample t-test against 0, RL *p* = 0.0005, UL *p*<0.0001) (**bottom right**) Minimum Z-Score over [0.1-3s] (Paired t-test *p* = 0.0245 ; One sample t-test against 0, RL *p* = 0.0008). **r**, Correlation between AUC [0.1 ; 2s] (RL-UL) difference and lever preference index during early instrumental training (n=9; Pearson’s r(7) = 0.5514, p=0.1238). **s**, Correlation between AUC [0.1 ; 2s] (RL-UL) difference and lever preference index during early reversal learning (n=9; Pearson’s r(7) = 0.8560, p=0.0032). All datasets represent mean values +/- s.e.m. Two-Factor RM ANOVA and Paired t-test \**p*<0.05, \*\**p*<0.01, \*\*\**p*<0.001, \*\*\*\**p*<0.0001. One sample t-test against 0, $ *p*<0.05, $$ *p*<0.01, $$$ *p*<0.001, $$$$ *p*<0.0001. Detailed statistics are shown in Supplementary Table 1. CS, conditioned stimulus; dmPFC, dorsomedial prefrontal cortex; ITI, inter trial interval; LP, lever press; RL, reinforced lever; Rwd, reward; UL, unreinforced lever; US, unconditioned stimulus.

Altogether, these data support that dopamine transmission in the dmPFC might influence reversal learning by signaling changes in action-outcome association and/or actions value. In this regard, neurons of the dmPFC have been shown to support flexible action selection by predicting outcomes associated to a particular action^35^. However, to which extent dopamine orchestrates such activity to allow for encoding of novel action-outcome associations is unclear. The activity of dmPFC neurons is largely driven by glutamatergic signals, while dopamine modulates their actions. Therefore, heteromers formed by dopamine and NMDA receptors are key candidates to allow for the integration of glutamate and dopamine inputs onto dopaminoceptive neurons as they function as coincidence detectors of both signals, facilitating neuronal adaptations^26,27,36^. We therefore assessed how mPFC dopaminoceptive neurons encode action-outcome relationship and how D1/NMDA and D2/NMDA heteromers modulate such activity.

### The distinct activity patterns of D1R+ and D2R+ dmPFC neurons during reversal learning are shaped by D1/NMDA and D2/NMDA heteromerization

DmPFC dopaminoceptive neurons are greatly regulated by glutamate signaling while dopamine modulates their activity. We hypothesized that crosstalk between dopamine and glutamate transmissions could be mediated by D1/NMDA and D2/NMDA receptor heteromers. We quantified the expression of these complexes in prefrontal and striatal subregions through proximity-ligation assay. D1/NMDA and D2/NMDA levels across the mPFC were found in amounts comparable to that of the striatal subregions (Supplementary Fig.4a-c), despite the lower cortical expression of dopamine receptors^37,38^, suggesting a higher heteromerization density in the mPFC.

We recorded bulk calcium dynamics of D1R- or D2R-expressing neurons of the dmPFC through expression of the calcium sensor GCaMP8m, coupled with fiber photometry in animals subjected to Pavlovian and instrumental conditioning paradigms. To assess the implication of dopamine-glutamate crosstalk on the activity of dmPFC dopaminoceptive neurons, we used an interfering peptide strategy to prevent D1/NMDA or D2/NMDA heteromerization^26^. Activity of dopaminoceptive neurons was first characterized in control conditions, i.e. under unilateral expression of control peptides for either heteromer (Fig.3a-c). During Pavlovian conditioning, while reward consumption was accompanied by a strong decrease in activity of both subpopulations (Supplementary Fig.6a-b,e-f), CS onset triggered a slight but prolonged increase of calcium dynamics in both D1R+ and D2R+ neurons in early sessions, that sharpened with training (Fig.3d-g). In instrumental training, dmPFC D1R+ and D2R+ neurons displayed a strong response induced by lever pressing on the RL which corresponds with CS onset (Fig.3h-k). However, this response decreased with training, while the slight increase in activity during action engagement remained stable over sessions (Fig.3h-k). These data suggest that rise in activity of dopaminoceptive neurons during lever press-induced CS might constitute a reinforcement learning signal, diminishing over learning of stable action-outcome associations.

**Figure 3:**
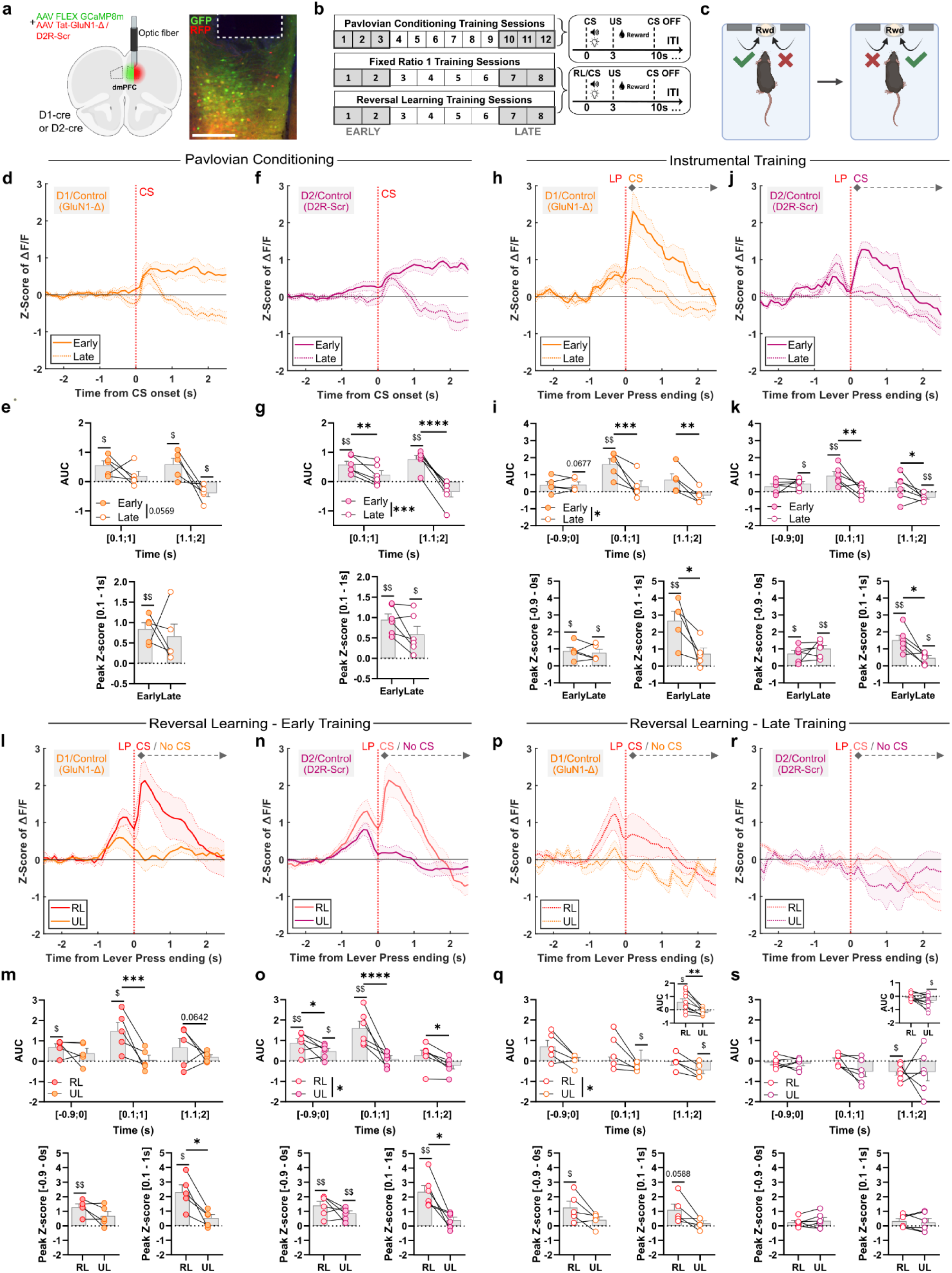
Recording of bulk activity of dmPFC dopaminoceptive neurons reveals a specific pattern of activity for D1R+ and D2R+ neurons during reversal learning. **a**, (**left**) Schematic representation of the viral vector strategy to record bulk calcium activity of D1R+ or D2R+ neurons in the dmPFC, (**right**) histological example of the expression of GCaMP8m (GFP) together with a control peptide (RFP), scale bar = 250 µm. **b**, Experimental design, the early stage corresponds to the first 3 sessions and late stage to the last 3 sessions for pavlovian conditioning, for instrumental training and reversal learning, the early stage corresponds to the first 2 sessions and the late stage to the last 2 sessions. CS-US sequence for pavlovian conditioning and RL/CS-US sequence for instrumental training. **c**, Schematic representation of instrumental training followed by reversal learning. **d-g**, Pavlovian conditioning (D1R+*_Control_*, *n* = 5 mice, D2R+*_Control_*, *n* = 6 mice) (**d**) Z-Score of ΔF/F for D1R+ neurons, (**e**) (**top**) Area Under the Curve (Two-Factor RM ANOVA, Training Effect F(1, 4) = 7.027 *p* = 0.0569 ; One sample t-test against 0, [0.1 s ; 1 s] Early *p* = 0.0198, [1.1 s ; 2 s] Early *p* = 0.0477, [1.1 s ; 2 s] Late *p* = 0.0347) ; (**bottom**) Peak Z-Score [-0.9s ; 0s] (One-sample t-test against 0, Early *p* = 0.00053) (**f**) Z-Score of ΔF/F for D2R+ neurons, (**g**) (**top**) Area Under the Curve (Two-Factor RM ANOVA, Training Effect F(1, 5) = 78.47 *p* = 0.0003, Time x Training F(1, 5) = 73.39 *p* = 0.0004, Šídák’s multiple comparisons test Early vs. Late [0.1 s ; 1 s] *p* = 0.0054, [1.1 s ; 2 s] *p* < 0.0001; One sample t-test against 0, [0.1 s ; 1 s] Early *p* = 0.0042, [1.1 s ; 2 s] Early *p* = 0.0023) ; (**bottom**) Peak Z-Score [0.1s ; 1s] (One sample t-test against 0, Early *p* = 0.0042, Late *p* = 0.0279). **h-k**, Instrumental training (D1R+*_Control_*, *n* = 5 mice, D2R+*_Control_*, *n* = 6 mice) (**h**) Z-Score of ΔF/F for D1R+ neurons, (**i**) (**top**) Area Under the Curve (Two-Factor RM ANOVA, Training Effect F(1, 4) = 17.05 *p* = 0.0145, Time x Training F(2, 8) = 8.499 *p* = 0.0105, Tukey’s multiple comparisons test, Early vs. Late [0.1 s ; 1 s] *p* = 0.0005, [1.1 s ; 2 s] *p* = 0.0039 ; One sample t-test against 0, [-0.9 s ; 0 s] Late *p* = 0.0677 [0.1 s ; 1 s] Early *p* = 0.0077), (**bottom left**) Peak Z-Score [- 0.9s ; 0s] (One-sample t-test against 0, Early *p* = 0.0169, Late *p* = 0.0216) and (**bottom right**) Peak Z- Score [0.1s ; 1s] (One-sample t-test against 0, Early *p* = 0.0089, Two-tailed Paired t-test *p* = 0.0388), (**j**) Z-Score of ΔF/F for D2R+ neurons, (**k**) (**top**) Area Under the Curve (Two-Factor RM ANOVA, Time x Training F(2, 10) =7.506 *p* = 0.0102, Tukey’s multiple comparisons test, Early vs. Late [0.1 s ; 1 s] *p* = 0.0013, [1.1 s ; 2 s] *p* = 0.0105 ; One sample t-test against 0, [-0.9 s ; 0 s] Late *p* = 0.0117, [0.1 s ; 1 s] Early *p* = 0.0087, [1.1 s ; 2 s] *p* = 0.0074), (**bottom left**) Peak Z-Score [-0.9s ; 0s] (One-sample t-test against 0, Early *p* = 0.0127, Late *p* = 0.0034) and (**bottom right**) Peak Z-Score [0.1s ; 1s] (One-sample t-test against 0, Early *p* = 0.0033, Late *p* = 0.0167, Two-tailed Paired t-test *p* = 0.0191). **l-o**, Reversal Learning early training (D1R+*_Control_*, *n* = 5 mice, D2R+*_Control_*, *n* = 6 mice) (**l**) Z-Score of ΔF/F for D1R+ neurons, (**m**) (**top**) Area Under the Curve (Two-Factor RM ANOVA, Time x Lever F(2, 8) = 7.442 *p* = 0.0149, Tukey’s multiple comparisons test, RL vs. UL [0.1 s ; 1 s] *p* = 0.0002, [1.1 s ; 2 s] *p* = 0.0642 ; One sample t-test against 0, [-0.9 s ; 0 s] RL *p* = 0.0154, [0.1 s ; 1 s] RL *p* = 0.0087, [0.1 s ; 1 s] UL *p* = 0.0250) ; (**bottom left**) Peak Z-Score [-0.9s ; 0s] (One-sample t-test against 0, RL *p* = 0.0051) and (**bottom right**) Peak Z-Score [0.1s ; 1s] (One-sample t-test against 0, RL *p* = 0.0105, Two-tailed Paired t-test *p* = 0.0148), (**n**) Z-Score of ΔF/F for D2R+ neurons, (**o**) (**top**) Area Under the Curve (Two-Factor RM ANOVA, Lever Effect F(1, 5) = 15.75 *p* = 0.0106, Time x Lever F(2, 10) = 11.45 *p* = 0.0149, Tukey’s multiple comparisons test, RL vs. UL [-0.9 s ; 0 s] *p* = 0.0460, [0.1 s ; 1 s] *p* < 0.0001, [1.1 s ; 2 s] *p* = 0.0179 ; One sample t-test against 0, [-0.9 s ; 0 s] RL *p* = 0.0084, [-0.9 s ; 0 s] UL *p* = 0.0195, [0.1 s ; 1 s] RL *p* = 0.0050) ; (**bottom left**) Peak Z-Score [-0.9s ; 0s] (One-sample t-test against 0, RL *p* = 0.0044, UL *p* = 0.0048) and (**bottom right**) Peak Z-Score [0.1s ; 1s] (One-sample t-test against 0, RL *p* = 0.0031, Two-tailed Paired t-test *p* = 0.0120). **p-s**, Reversal Learning late training (D1R+*_Control_*, *n* = 5 mice, D2R+*_Control_*, *n* = 6 mice) (**p**) Z-Score of ΔF/F for D1R+ neurons, (**q**) (**top**) Area Under the Curve (Two-Factor RM ANOVA, Lever Effect F (1, 4) = 8.690 *p* = 0.0421 ; One sample t-test against 0, [0.1 s ; 1 s] UL *p* = 0.0339, [1.1 s ; 2 s] UL *p* = 0.0454) ; (**top insert**) Area under the curve, pool of [-0.9 s ; 0 s] and [0.1 s ; 1 s] AUC (One-sample t-test against 0, RL *p* = 0.0329, Two-tailed Paired t-test *p* = 0.0040) ; (**bottom left**) Peak Z-Score [-0.9s ; 0s] (One-sample t-test against 0, RL *p* = 0.0434) and (**bottom right**) Peak Z-Score [0.1s ; 1s] (One-sample t-test against 0, RL *p* = 0.0588), (**r**) Z-Score of ΔF/F for D2R+ neurons, (**s**) (**top**) Area Under the Curve (One sample t-test against 0, [1.1 s ; 2 s] RL *p* = 0.0210) ; (**top insert**) Area under the curve, pool of [-0.9 s ; 0 s] and [0.1 s ; 1 s] AUC (One-sample t-test against 0, UL *p* = 0.0422) ; (**bottom left**) Peak Z-Score [- 0.9s ; 0s] and (**bottom right**) Peak Z-Score [0.1s ; 1s]. All datasets represent mean values +/- s.e.m. Two-Factor RM ANOVA and Paired t-test \**p*<0.05, \*\**p*<0.01, \*\*\**p*<0.001, \*\*\*\**p*<0.0001. One sample t-test against 0, $ *p*<0.05, $$ *p*<0.01. Detailed statistics are shown in Supplementary Table 1. CS, conditioned stimulus; dmPFC, dorsomedial prefrontal cortex; ITI, inter trial interval; LP, lever press; RL, reinforced lever; Rwd, reward; UL, unreinforced lever; US, unconditioned stimulus.

Strikingly, in the early phase of reversal learning, bulk activity of both D1R+ and D2R+ neurons clearly distinguished between the newly-reinforced and previously-reinforced levers. Indeed, a significant response associated with lever pressing engagement, together with a strong increase after pressing on the RL, e.g. during CS onset, while pressing on the UL led to virtually no activity for both neuron subtypes (Fig.3l-o). These data highlight a clear distinction in the activity of both D1R+ and D2R+ neurons of the dmPFC associated with acquisition of the newly- and previously-reinforced levers in a reversal learning task. With training, activity of D1R+ neurons associated with the RL was maintained selectively during action engagement, but not at RL-induced CS, while there was virtually no activity – and even inhibition - for the UL (Fig.3p-q). Strikingly, there was no remaining activity of D2R+ neurons associated with either RL or UL for late stages of reversal training (Fig.3r-s), which clearly differed from D2R+ activity for late stages of initial training that was maintained during action engagement associated with RL (Fig.3j-k). These findings show distinct pattern for both dopaminoceptive subpopulations of the dmPFC specifically during reversal learning, as opposed to initial action-outcome association. This supports the recruitment of dopamine signaling during reversal learning, exerting its activating and inhibiting effect through D1R and D2R, respectively. We next sought to unravel whether such an effect depended on D1/NMDA and D2/NMDA heteromers.

Upon blockade of dmPFC D1/NMDA or D2/NMDA heteromers through the expression of interfering peptides using the same protocol (Fig.4a-c), training-induced sharpening of CS-evoked activity of D1R+ and D2R+ neurons for Pavlovian conditioning (Fig.4d-g) and dampening for instrumental training (after RL pressing) was preserved (Fig.4h-k), although less stringent compared to controls (Fig.3h-k). This suggests that D1/NMDA and D2/NMDA heteromers are not required for learning-induced adaptations of dmPFC dopaminoceptive neurons. During the early phase of reversal learning, expression of interfering peptides preserved the differential patterns of activity between RL and UL for both D1R+ and D2R+ neurons (Fig.4l-o), although less pronounced compared to controls (Fig.3l-o). Of note, in animals where dmPFC D2/NMDA were blocked, D2R+ neurons still displayed an activity above baseline after pressing on the UL, contrary to control animals (Fig.3n-o). However, during late reversal training, blockade of D1/NMDA heteromers resulted in the absence of activity of D1R+ neurons related to either RL or UL (Fig.4p-q), which differed from control animals for RL (Fig.3p-q). Conversely, blockade of D2/NMDA heteromers translated in the absence of activity of D2R+ neurons related to the RL (Fig.4r-s), similar to control animals (Fig.3r-s), surprisingly accompanied by significant responding associated with engagement in pressing for the UL and a global activity higher for UL compared to RL (Fig.4r-s), contrary to controls (Fig.3r-s). Overall, reward consumption triggered a comparable decrease of activity during pavlovian conditioning, instrumental training (Supplementary Fig.6a-p) and reversal learning, with lower amplitude of inhibition with learning (Supplementary Fig.6q-x). Of note, in these recording conditions with unilateral blockade of D1/NMDA or D2/NMDA in the mPFC, there was no alteration in behavioral performance (Supplementary Fig.5a-h).

**Figure 4:**
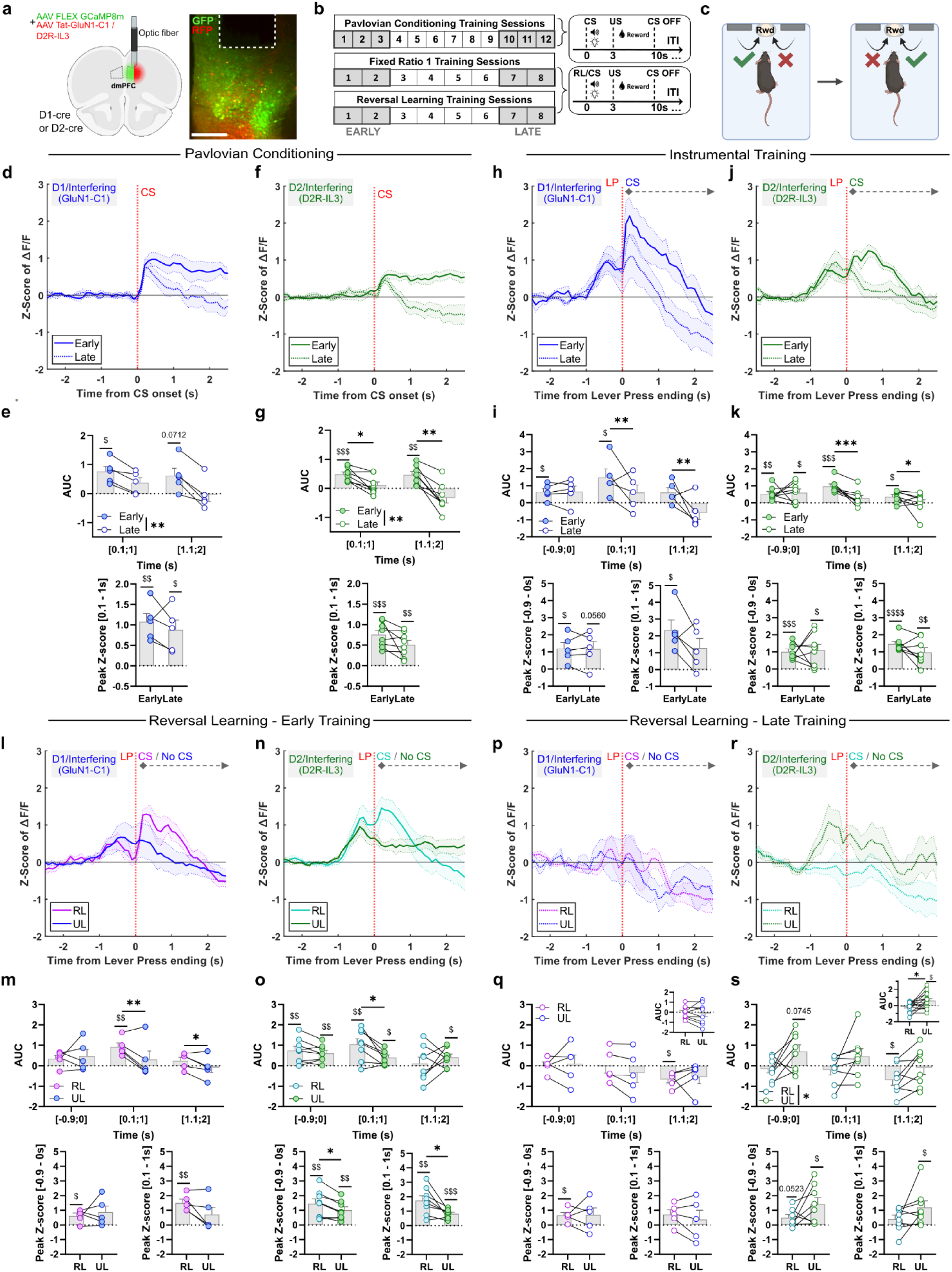
During reversal learning, dmPFC D1R+ and D2R+ neuronal signature is mediated by D1/NMDA and D2/NMDA heteromers, respectively. **a**, (**left**) Schematic representation of the viral vector strategy to record bulk calcium activity of D1R+ or D2R+ neurons in the dmPFC, (**right**) histological example of the expression of GCaMP8m (GFP) together with an interfering peptide (RFP), scale bar = 250 µm. **b**, Experimental design, the early stage corresponds to the first 3 sessions and late stage to the last 3 sessions for pavlovian conditioning, for instrumental training and reversal learning, the early stage corresponds to the first 2 sessions and the late stage to the last 2 sessions. CS-US sequence for pavlovian conditioning and RL/CS-US sequence for instrumental training. **c**, Schematic representation of instrumental training followed by reversal learning. **d-g**, Pavlovian conditioning (D1R+*_Interfering_*, *n* = 5 mice, D2R+*_Interfering_*, *n* = 8 mice) (**d**) Z-Score of ΔF/F for D1R+ neurons, (**e**) (**top**) Area Under the Curve (Two-Factor RM ANOVA, Training Effect F(1, 4) = 47.61 *p* = 0.0023 ; One sample t-test against 0, [0.1 s ; 1 s] Early *p* = 0.0141, [1.1 s ; 2 s] Early *p* = 0.0712) (**bottom**) Peak Z-Score [0.1s ; 1s] One sample t-test against 0, [0.1 s ; 1 s] Early *p* = 0.0198, [0.1 s ; 1 s] Late *p* = 0.0215), (**f**) Z-Score of ΔF/F for D2R+ neurons, (**g**) (**top**) Area Under the Curve (Two-Factor RM ANOVA, Training Effect F(1, 7) = 28.77 *p* = 0.0010, Time x Training F(1, 7) = 20.30 *p* = 0.0028, Šídák’s multiple comparisons test, Early vs. Late [0.1 s ; 1 s], *p* = 0.0211 [1.1 s ; 2 s] *p* = 0.0030 ; One sample t-test against 0, [0.1 s ; 1 s] Early *p* = 0.0006, [1.1 s ; 2 s] Early *p* = 0.0042) (**bottom**) Peak Z-Score [0.1s ; 1s] (One sample t-test against 0, Early *p* = 0.0002, Late *p* = 0.0019). **h-k**, Instrumental training (D1R+*_Interfering_*, *n* = 5 mice, D2R+*_Interfering_*, *n* = 8 mice) (**h**) Z-Score of ΔF/F for D1R+ neurons, (**i**) ) (**top**) Area Under the Curve (Two-Factor RM ANOVA, Time x Training F(2, 8) = 6.077 *p* = 0.0248, Tukey’s multiple comparisons test, Early vs. Late [0.1 s ; 1 s], *p* = 0.0082, [1.1 s ; 2 s] *p* = 0.0016 ; One sample t-test against 0, [-0.9s ; 0s] Early *p* = 0.0356, [0.1 s ; 1 s] Early *p* = 0.0393), (**bottom left**) Peak Z-Score [-0.9s ; 0s] (One-sample t-test against 0, Early *p* = 0.0295, Late *p* = 0.0560) and (**bottom right**) Peak Z-Score [0.1s ; 1s] (One-sample t-test against 0, Early *p* = 0.0174), (**j**) Z-Score of ΔF/F for D2R+ neurons, (**k**) (**top**) Area Under the Curve (Two-Factor RM ANOVA, Time x Training F(2, 14)=8.324 *p* = 0.0042, Tukey’s multiple comparisons test, Early vs. Late [0.1 s ; 1 s], *p* = 0.0001, [1.1 s ; 2 s] *p* = 0.0160 ; One sample t-test against 0, [-0.9s ; 0s] Early *p* = 0.0058, [-0.9s ; 0s] Late *p* = 0.0269, [0.1 s ; 1 s] Early *p* = 0.0002, [1.1 s ; 2 s] Early *p* = 0.0206), (**bottom left**) Peak Z-Score [-0.9s ; 0s] (One-sample t-test against 0, Early *p* = 0.0004, Late *p* = 0.0170) and (**bottom right**) Peak Z-Score [0.1s ; 1s] (One-sample t-test against 0, Early *p* < 0.0001, Late *p* = 0.0079). **l-o**, Reversal Learning early training (D1R+*_Interfering_*, *n* = 5 mice, D2R+*_Interfering_*, *n* = 8 mice) (**l**) Z-Score of ΔF/F for D1R+ neurons, (**m**) (**top**) Area Under the Curve (Two-Factor RM ANOVA, Time x Lever F(2, 8) = 8.469 *p* = 0.0106, Tukey’s multiple comparisons test, RL vs. UL [0.1 s ; 1 s], *p* = 0.0015, [1.1 s ; 2 s] *p* = 0.0380 ; One sample t-test against 0, [0.1 s ; 1 s] RL *p* = 0.0092), (**bottom left**) Peak Z-Score [-0.9s ; 0s] (One-sample t-test against 0, RL *p* = 0.0383) and (**bottom right**) Peak Z-Score [0.1s ; 1s] (One-sample t-test against 0, RL *p* = 0.0043), (**n**) Z-Score of ΔF/F for D2R+ neurons, (**o**) (**top**) Area Under the Curve (Two-Factor RM ANOVA, Time x Lever F(2, 14) = 4.004 *p* = 0.0422, Tukey’s multiple comparisons test, RL vs. UL [0.1 s ; 1 s], *p* = 0.0164 ; One sample t-test against 0, [-0.9s ; 0s] RL *p* = 0.0096, [-0.9s ; 0s] UL *p* = 0.0037, [0.1 s ; 1 s] RL *p* = 0.0059, [0.1 s ; 1 s] UL *p* = 0.0140, [1.1 s ; 2 s] UL *p* = 0.0352), (**bottom left**) Peak Z-Score [-0.9s ; 0s] (One-sample t-test against 0, RL *p* = 0.0033, UL *p* = 0.0030, Two-tailed Paired t-test *p* = 0.0117) and (**bottom right**) Peak Z-Score [0.1s ; 1s] (One-sample t-test against 0, RL *p* = 0.0013, UL *p* = 0.0002, Two-tailed Paired t-test *p* = 0.0184). **p-s**, Reversal Learning late training (D1R+*_Interfering_*, *n* = 5 mice, D2R+*_Interfering_*, *n* = 8 mice) (**p**) Z-Score of ΔF/F for D1R+ neurons, (**q**) (**top**) Area Under the Curve (One sample t-test against 0, [1.1 s ; 2 s] RL *p* = 0.0162) ; (**top insert**) Area under the curve, pool of [-0.9 s ; 0 s] and [0.1 s ; 1 s] AUC ; (**bottom left**) Peak Z-Score [-0.9s ; 0s] (One-sample t-test against 0, RL *p* = 0.0383 and (**bottom right**) Peak Z-Score [0.1s ; 1s], (**r**) Z-Score of ΔF/F for D2R+ neurons, (**s**) (**top**) Area Under the Curve (Two-Factor RM ANOVA, Lever Effect F(1, 7) = 9.748 *p* = 0.0168 ; One sample t-test against 0, [-0.9s ; 0s] UL *p* = 0.0745, [1.1 s ; 2 s] RL *p* = 0.0285) ; (**top insert**) Area under the curve, pool of [-0.9 s ; 0 s] and [0.1 s ; 1 s] AUC (One-sample t-test against 0, UL *p* = 0.0283, Two-tailed Paired t-test *p* = 0.0102) ; (**bottom left**) Peak Z-Score [-0.9s ; 0s] (One-sample t-test against 0, RL *p* = 0.0523, UL *p* = 0.0185) and (**bottom right**) Peak Z-Score [0.1s ; 1s] (One-sample t-test against 0, UL *p* = 0.0325). All datasets represent mean values +/- s.e.m. Two-Factor RM ANOVA and Paired t-test \**p*<0.05, \*\**p*<0.01, \*\*\**p*<0.001. One sample t-test against 0, $ *p*<0.05, $$ *p*<0.01, $$$ *p*<0.001, $$$$ *p*<0.0001. Detailed statistics are shown in Supplementary Table 1. CS, conditioned stimulus; dmPFC, dorsomedial prefrontal cortex; ITI, inter trial interval; LP, lever press; RL, reinforced lever; Rwd, reward; UL, unreinforced lever; US, unconditioned stimulus.

Altogether, these findings show that D1/NMDA and D2/NMDA heteromers are not required for the activity of dmPFC dopaminoceptive neurons associated with learning of the initial cue- or action-outcome association. However, on the one hand D1/NMDA heteromers are central for allowing the maintenance of activity of dmPFC D1R+ neurons associated with learning of the newly-reinforced action. On the other hand, D2/NMDA heteromers are necessary for repressing the activity of dmPFC D2R+ neurons associated with the previously-reinforced action.

### Interfering with D1/NMDA or D2/NMDA heteromerization in the dmPFC selectively impairs reversal of A-O associations

To assess the behavioral consequence of D1/NMDA or D2/NMDA heteromer blockade, either of interfering peptides were bilaterally expressed in the dmPFC (Fig.5a-b), and animals were subjected to the same behavioral procedures as described previously (Fig.1), to assess acquisition and expression of action-outcome association, as well as reversal of outcome identities (Fig.5c). Blocking dmPFC D1/NMDA or D2/NMDA did not impact instrumental responding (Fig.5d), nor action-outcome association and expression as all groups equally performed in the devaluation test (Fig.5e-f, Supplementary Fig.7a-b). However, despite comparable operant responding in reversal learning sessions (Fig.5g), blockade of either D1/NMDA or D2/NMDA heteromers impaired discrimination between the valued and devalued levers in the outcome devaluation test (Fig.5h-i). This effect was not due to a general motivational deficit or an unspecific satiety as all groups displayed similar consumption in prefeeding and in the post-extinction feeding test (Supplementary Fig.7c-d). Altogether this suggests that D1/NMDA and D2/NMDA heteromers control the ability to properly update action-outcome association without affecting initial goal-directed action.

**Figure 5:**
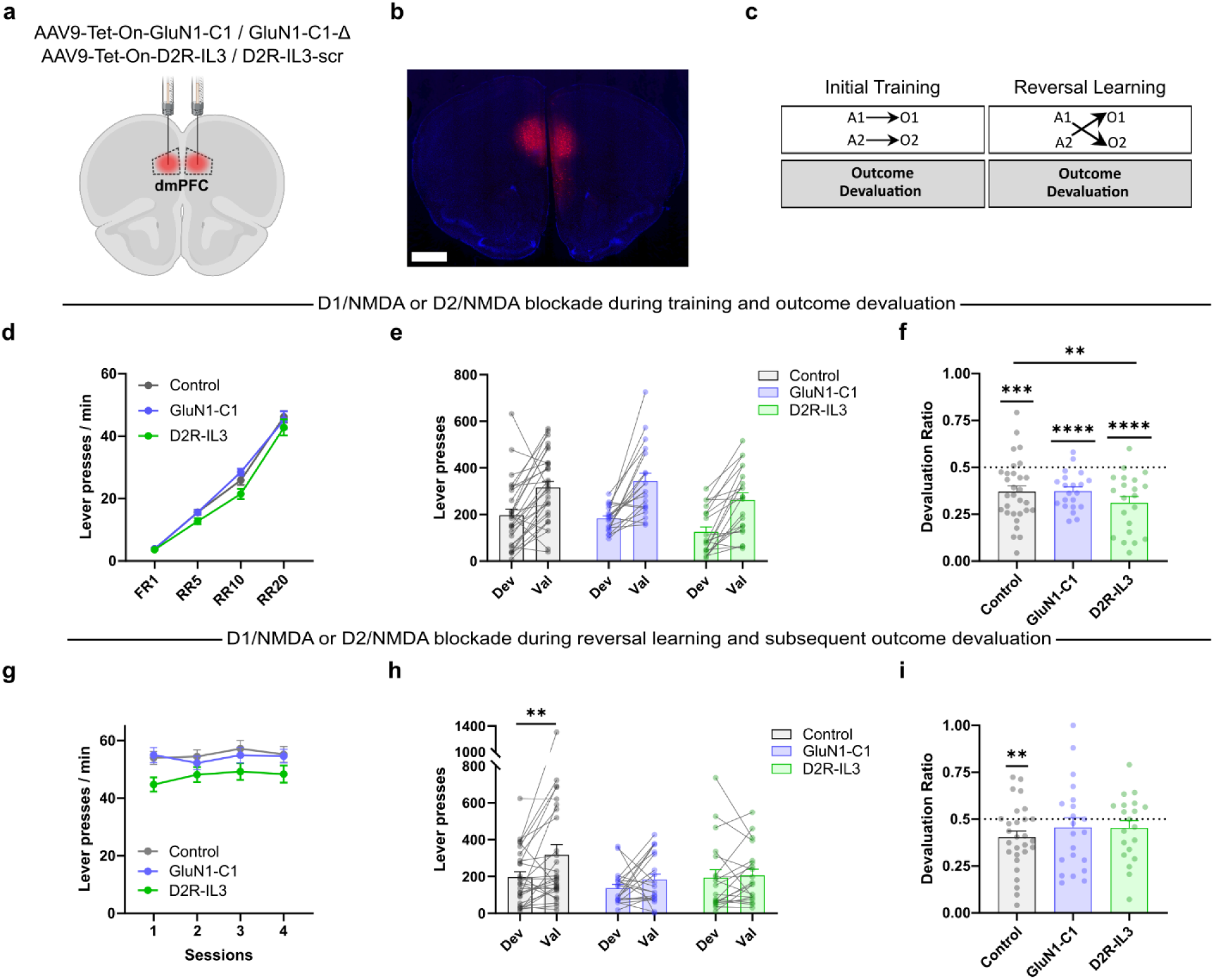
Blocking D1/NMDA or D2/NMDA heteromerization in the dmPFC selectively impairs reversal of action-outcome associations. **a**, Representation of the viral vector injections in the dmPFC. **b**, Histological example of the expression of the peptide, scale bar = 1 mm. **c**, Design of the behavioral procedure. **d**, Lever presses frequency during instrumental training (Control Peptides (n=30), GluN1-C1 (n=21), D2R-IL3 (n=21), Two-Factor RM ANOVA, Training effect F(3,207) = 542.8, *p*<0.0001, Peptide effect F(2,69) = 2.926, *p* = 0.0603). **e-f**, Extinction test during the outcome devaluation procedure (Control Peptides (n=30), GluN1-C1 (n=21), D2R-IL3 (n=21)) (**e**) amount of lever presses (Planned orthogonal contrast, within group effect F(1,69) = 75.444, *p*<0.0001) and (**f**) devaluation ratio (Planned orthogonal contrast, interaction F(2,69) = 3.128, *p* = 0.0501, simple effect analysis, Control vs D2R-IL3 F(2,69) = 5.660, *p* = 0.0053, GluN1-C1 vs D2R-IL3 F(2,69) = 2.711, *p* = 0.0736 ; One sample t-test against 0.5, Control *p*<0.001, GluN1-C1 *p*<0.0001, D2R-IL3 *p*<0.0001). **g**, Reversal learning training (Control Peptides (n=28), GluN1-C1 (n=21), D2R-IL3 (n=20), Two-Factor RM ANOVA, Training effect F(3,198) = 2.728, *p =* 0.0452, Peptide effect F(2,66) = 2.747, *p* = 0.0715). **h-i**, Extinction test during the outcome devaluation procedure (Control Peptides (n=28), GluN1-C1 (n=21), D2R-IL3 (n=20)) (**h**) amount of lever presses on the Dev and Val levers (Planned orthogonal contrast, interaction F(2,66) =3.939, *p* = 0.0242, simple effect analysis, Control F(1,66) = 11.664, *p* = 0.0011, GluN1-C1 F(1,66) = 1.337, *p* = 0.2517, D2R-IL3 F(1,66) = 0.085, *p* = 0.7715) and (**i**) devaluation ratio (One sample t-test against 0.5, Control *p* = 0.0072, GluN1-C1 *p* = 0.3965, D2R-IL3 *p* = 0.2426). All datasets represent mean values +/- s.e.m. \*\**p*<0.01, \*\*\**p*<0.001, \*\*\*\**p*<0.0001. A, action; Dev, devalued; dmPFC, dorsomedial prefrontal cortex; FR, fixed ratio; O, outcome; RR, random ratio; Val, valued. Detailed statistics are shown in Supplementary Table 1.

## DISCUSSION

The current findings bring evidence that dopamine transmission in the dmPFC participates in decision-making processes by signaling action-outcome association, in particular during changes in goal-directed actions. Integration of such dopamine signal by dopaminoceptive neurons is mediated by D1/NMDA and D2/NMDA heteromer, with activity of D1R-expressing neurons correlating with encoding of newly-reinforced actions, while D2R-expressing neurons become inhibited in relation with previously-reinforced actions. Consequently, blockade of either heteromers spares learning and expression of initial instrumental associations but selectively impairs adjustment to reversal of action-outcome associations. These data bring evidence that mPFC D1/NMDA and D2/NMDA heteromers mediate the effect of dopamine signaling for adaptation to changes in action-outcome relationships.

The findings that inhibition of mesocorticolimbic dopamine transmission is not critical for acquisition of action-outcome associations or their expression is supported by previous studies^9,15,39^. However, while some findings reveal that manipulations of dopamine transmission of the mPFC, particularly in its dorsal part (dmPFC), perturb adaptation to changes in action-outcome relationship probed through contingency degradation^15^, other studies found no effect in such adaptive behavior^16^. In this study, we provide evidence that dmPFC dopamine modulates adaptation to changes in action-outcome relationships by encoding reward prediction specifically when associated to an action, suggesting that it allows reinforcing action value. Indeed, we found virtually no change in dopamine signal at the CS during pavlovian conditioning. These results contrast with previous report showing significant signal in mPFC dopamine terminals in response to both reward and reward-predicting cue^40^. However, more recent findings show that reward-responding dopamine axons in the mPFC are not involved in reward prediction^32^. Such discrepancies might originate from the nature or the activation patterns of the mesocortical dopamine projections that might recruit different mechanisms, notably co-release of glutamate as previously shown^41–43,44^.

In accordance with a main role of dopaminergic projections in encoding changes in action causality, our dopamine recordings highlight a more discriminant signal between reinforced and non-reinforced actions following an inversion of action–outcome associations compared to initial contingencies. Such a difference not only translates in a higher increase in amplitude of dopamine release following the action – that correlates with correct lever preference in early training – but also through a significant signal inhibition (below baseline) before and after reinforced responses. Such a temporal sharpening of the dopamine response might represent more reliable signal for computation by dopaminoceptive neurons, enhancing discrimination between actions, and therefore optimal encoding of action value. This is particularly relevant in a reversal learning paradigm in which interference is high compared to initial instrumental learning. Although we cannot completely rule out that this effect is a consequence of prolonged instrumental training, such a main role of dmPFC dopamine transmission is in line with previous findings showing that dopamine efflux in the mPFC is elevated in early phases of reversal learning^45^. Moreover, phasic activation of mesocortical dopaminergic projections deviates behavior from previously learned associations^40^ and these projections are associated with the regulation of perseverative behaviors^43^.

Consistent with glutamate signals being one of the main drivers, activity of dmPFC dopaminoceptive neurons was mostly disconnected from dopamine dynamics. First, both dopaminoceptive subpopulations mainly displayed comparable activity patterns, while dopamine transmission is expected to exert excitatory and inhibitory effects through D1R and D2R, respectively. Moreover, while dmPFC dopamine displayed virtually no response to reward-predicting cues but significant increase at reward consumption, calcium dynamics in both D1R+ and D2R+ neurons were significantly increased in response to the CS in a pavlovian conditioning paradigm, but were inhibited during reward consumption. Those results are in line with studies showing that a proportion of dmPFC neurons are activated at CS onset while their activity decreases during reward consumption^46,47^. CS-induced activity was even more elevated when the outcome became contingent to an action, in accordance with dmPFC neurons being more sensitive to reward prediction in the situation of operant control^48^. This activity dampened with training, suggesting that dmPFC dopaminoceptive neurons become less recruited once the association between the action and the reward is established. This is in line with dmPFC neuronal ensembles narrowing over learning of an associative learning task^49^. Accordingly, activity of both D1R+ and D2R+ neurons recapitulated comparable patterns of activity for the newly-reinforced action when associations between action and outcome were reversed, i.e. when they have to be re-learned. However, in late phases of reversal training, only D1R+ neurons maintained activity related to the reinforced action, in comparison to D2R+ neurons that became virtually silent. This is in accordance with the well-established implication of dmPFC dopaminergic receptors in behavioral flexibility^15,18,19,50,51^. In addition, our results support previously-proposed complementary roles of D1R+ and D2R+ neurons in adaptations to changes in action-outcome relationships, the former being necessary to encode a new/updated association, while inhibition of the latter is necessary in order to disengage a formerly rewarded association that is no longer relevant^18,19,50,50,52^. These effects are likely to be mediated by facilitating and inhibitory action of dopamine through D1R and D2R, respectively. In accordance, blockade of D1/NMDA heteromers prevented the maintenance of activity of D1R+ neurons associated with late reversal training, while interfering with D2/NMDA complexes precluded inhibition of the response associated with the previously-reinforced action. These effects on dopaminoceptive neurons could account for selective deficits in reversal learning under each heteromer blockade. Our results are therefore in line with the dual-state theory of PFC dopamine that proposes that in a D1R-state, dopamine enhances the stability of prefrontal representations to sustain a relevant response, while a D2R-state allows representations to fluctuate and become less stable, thus enabling shifting between different strategies^53,54^.

In accordance with this view, our behavioral data show that dmPFC D1/NMDA and D2/NMDA heteromers are not necessary for adjustment of neuronal activity to new contingencies, but are specifically required to allow the maintenance of specific signature of dopaminoceptive neurons over reversal training. In other words, D1/NMDA and D2/NMDA heteromers seem to play a main role for the consolidation of updated associations. This is in accordance with the known role of these heteromers for neuronal adaptations. In the striatum, while D1/NMDA favor long term potentiation in D1-expressing medium spiny neurons (D1-MSNs)^27^, D2/NMDA tend to blunt D2-MSNs transmission^29^. Numerous studies have shown a facilitating effect of D1R activation on NMDA-dependent transmission in the PFC^55–57^, while D2R have the opposite effect^57,58^. It is therefore tempting to propose that the effects we found under heteromers blockade during reversal training, i.e. blunting of the activity of D1R+ neurons at the newly-reinforced lever, but maintenance of activity of D2R+ neurons for the previously-reinforced action, are a direct consequence of impaired plasticity mediated by D1/NMDA and D2/NMDA, respectively. Following this hypothesis, the inability to properly reverse associations would then be caused by a perturbation in the encoding of the reverse association in the case of D1/NMDA heteromer blockade, while being related to an impairment in repressing the previously-learned associations under D2/NMDA heteromer blockade.

These findings raise the question of the mPFC neuronal subpopulations involved, as well as putative downstream circuits. In the mPFC, D1R and D2R are expressed in mostly segregated populations with less than 15% coexpression in the dmPFC^17^. D1R mRNA is expressed in all layers but mainly in deeper layers (V and VI), while D2R mRNA can be found mostly in layer V^37^. While we cannot rule out that part of the recorded activity of either D1R+ or D2R+ neurons stems from GABAergic interneurons^37,59,60^, pyramidal projection neurons are likely to underlie the majority of the recorded signal^37^. These considerations also apply to the expression of D1/NMDA or D2/NMDA heteromers on the different subpopulations of dopaminoceptive neurons. Regarding downstream target regions, on the one hand, D1R+ neurons of the caudal dmPFC mainly project to the posterior dorsomedial striatum^59,61,62^, on the other hand, D2R+ neurons strongly project to the ventral striatum^60,61^ and to a lesser extent to the mediodorsal striatum^63^ , with both striatal subregions being implicated in updating of action-outcome associations in reversal learning^64–66^. Further studies will be necessary to unravel how these circuits cooperate to regulate reversal learning. Moreover, while our findings establish a role of dmPFC dopaminoceptive neurons for acquisition/consolidation of updated action-outcome associations through the recruitment of D1/NMDA and D2/NMDA heteromers, the roles of these neuronal populations in more dynamic adaptation to fast-changing situations remain to be established. Indeed, large body of literature emphasizes the role of the dmPFC to rule switching in working memory-based tasks that require on-line adjustment of action-outcome associations^30^, which has been shown to depend on dopamine transmission^18,19^.

Evidence demonstrated the implication of D1/NMDA and D2/NMDA heteromers in the ventral striatum in the context of addiction, particularly in psychostimulant-induced behavioral adaptations ^26^. A growing body of evidence points towards a concomitant dysregulation of both the glutamatergic and dopaminergic transmission in several psychiatric disorders^7,8,67^. Our findings, showing that these receptor complexes are main molecular platforms to mediate the crosstalk between glutamatergic and dopaminergic signals, therefore call for further investigation of the implication of those heteromers in psychiatric disorders and pave the way for potential future treatment strategies.

## Acknowledgements

We thank the Bordeaux Imaging Center (a service unit of the CNRS-INSERM and Bordeaux University, member of the national infrastructure France BioImaging) support by ANR-10-INBS-04 for imaging, Dr. Nicolas Heck for helping with PLA quantification (Sorbonne Université UMR 8265 – NeuroSU), Céline Ducroix-Crépy, Eva Bruchet and Jean-Christophe Helbling (INRAE UMR 1286) for genotyping, the staff from the animal facility of INRAE UMR 1286 for animal care and the CIRCE (Behavioural Engineering Center) facility of the Bordeaux Neurocampus. The authors wish to thank Dr. Shauna Parkes and Dr. Etienne Coutureau for their help with experimental design, statistical analyses and interpretation of the results.

## Authors contributions

AP, AC, RW, GF, AA, JB, PV and PT designed research. AP, AC, MCA, LH, FD, PC, AO, MFA, MC, MD, YH, AI, FA, JCV, RO, RT, AB, MR performed research. AP, AC, PT supervised research. AP, AC, RW, MCA, LH, FD, PC, AO, MC, MD, YH, AI, FA, JCV, RO, RT, AB, MR analyzed data. AP, FD, RW, MCA, GF, PV, JB and PT wrote the manuscript. All authors edited and approved the manuscript.

## MATERIAL AND METHODS

### Experimental models and subject details

Adult male mice aged 10 – 28 weeks were group housed (3-10 animals) in standard polypropylene cages and maintained in a temperature (21-23°C) and humidity-controlled facility under a 12:12 light-dark cycle (07:00 on) with *ad libitum* access to food and water. All animal care and experimental procedures were in accordance with the INRAE Quality Reference System and with French (Directive 87/148, Ministère de l’Agriculture et de la Pêche) and European (Directive 86/609/EEC) legislations. They followed ethical protocols approved by the Region Aquitaine Veterinary Services (Direction Départementale de la Protection des Animaux, approval ID: B33-063-920) and by the animal ethic committee of Bordeaux CEEA50. Every effort was made to minimize suffering and reduce the number of animals used.

The following mouse lines were used in this study : C57BL/6J were purchased from Janvier Laboratories (Robert Janvier, Le Genest St-Isle, France), D1-Cre (MMRC: 029178-UCD, strain code: Tg(Drd1-cre)FK150Gsat/Mmucd, GENSAT), D2-Cre (MMRC: 017263-UCD, strain code: Tg(Drd2-cre)ER44Gsat/Mmucd, GENSAT), DAT-cre (DAT^IREScre^, B6.SJL-Slc6a3tm1.1(cre)Bkmn/J, RRID:IMSR_JAX:006660).

### Stereotactic viral injections and implantation of optic fibers

Mice analgesia was induced with the subcutaneous injection of Buprenorphine (BUPRECARE, 0.1 mg/kg) 30 minutes prior to the surgical procedure. Animals were then anesthetized with isoflurane (induction at 5%, procedure at 1-2%) and then placed on a stereotactic frame (RWD Life Science) on a heating pad (37°C). A 10µl NanoFil Sub-micorliter syringe from WPI was used with a 34G Nanofil blunt needle. AAV used and structures targeted are specified below. AAV were injected at a 100 nl/min rate. The syringe was left in place for 5 minutes post injection to prevent backflow and was then slowly removed. At the end of the procedure, postoperative analgesia was ensured with a subcutaneous injection of non-steroidal anti-inflammatory drug Carprofene (CARPROX, 5 mg/kg). Mice were sutured and placed on a heating pad until full recovery from the procedure, and were monitored for 3 days post-surgery. Behavioral procedures begun 3 – 4 weeks after the surgeries. Every procedure was used to minimize bacterial contamination and enforce prophylaxis.

### Dopaminergic neurons chemogenetic inhibition

For chemogenetic inhibition of VTA dopamine neurons during instrumental learning, DAT-cre positive male mice were injected with AAV8-hSyn1-dlox-hM4D(Gi)_mCherry(rev)-dlox-WPRE-hGHp(A) (6.3 x 10^12^ vg/ml, v84, ETH Zurich Viral Vector Facility) or AAV8-hSyn1-dlox-mCherry(rev)-dlox-WPRE-hGHp(A) (5.9×10¹^2^ vg/ml, v116, ETH Zurich Viral Vector Facility). n_Gi_ = 10 and n_mCherry_ = 10. Mice were injected in the VTA at AP : -3.30 ; ML : +/- 0.50 ; DV : -4.30 from the surface of the brain, at a volume of 300nl/hemisphere. For the chemogenetic inhibition during outcome devaluation and reversal learning, DAT-cre positive or negative mice were injected with AAV8-hSyn1-dlox-hM4D(Gi)_mCherry(rev)-dlox-WPRE-hGHp(A) (6.3 x 10^12^ vg/ml, v84, Zurich Vector Facility) in the VTA at the same coordinates and volumes. n_Cre+_ = 11 and n_Cre-_= 11.

### In vivo fiber photometry recording of mPFC dopamine dynamics

For fiber photometry experiments, C57BL/6J male mice received a unilateral injection of AAV-9/2-hSyn1-GRAB(DA2m)-WPRE-hGHp(A) (7x10^12^ vg/ml, v677-9, ETH Zurich Viral Vector Facility) vector in the mPFC. The following coordinates from bregma and volumes were used for AAV injections in the mPFC (AP : +1.78 ; ML : +/-0.30 ; DV : -2.40 ; 500nl). The optic fiber (Fiber Optic Cannula with Ceramic Ferrule, O.D. 1.25 mm, 0.5 NA, 400 µm core diameter, L = 3 mm; R-FOC-BL400C-50NA, RWD) was placed 250 µm above the injection site and immobilized with an opaque dental cement (Super-Bond, Sun Medical; Meliodent, Kulzer).

### D1/NMDA and D2/NMDA heteromer blockade

C57BL/6J male mice were injected with viral vectors expressing interfering peptides, previously validated in ^26^. Briefly, AAV9-Tet-On-GluN1C1 bicistronically bears the fluorescent reporter protein RFP and the C1 cassette of the GluN1 subunit (_864_DRKSGRAEPDPKKKATFRAITSTLASSFKRRRSSKDT_900_) under doxycycline treatment. The related control virus AAV9-Tet-On-GluN1C1Δ bears a truncated version of C1 that is deleted from a stretch of nine positively charged amino acids (_890_SFKRRRSSK_898_), which are required for D1R-NMDAR interaction. The AAV9-Tet-On-D2R-IL3 encodes a sequence of the IL3 of the D2R (_225_TKRSSRAFRA_234_) interacting with GluN2B. The control AAV9-Tet-On-D2R-scr bears a scrambled sequence (KFARRTSASR) of the D2R-IL3 (full AAV sequences are available upon request). All Tet-On AAV were injected at 5x10^13^ vg/ml in PBS 1X Pluronic 0.001%. The same mPFC coordinates as for expression of dopamine sensor were used and the injection volume was 400nl/hemisphere. Results correspond to a pool of three different cohorts.

### In vivo fiber photometry recording of calcium dynamics of mPFC dopaminoceptive neurons upon blockade of D1/NMDA or D2/NMDA heteromers

For fiber photometry experiments, D1-cre male mice received a unilateral injection of a mix of AAV-DJ-hSyn1-dlox-jGCaMP8m(rev)-dlox-WPRE-SV40p(A) (6x10^12^ vg/ml, v628, ETH Zurich Viral Vector Facility) and AAV9-Tet-On-GluN1C1 or AAV9-Tet-On-GluN1C1Δ vector in the mPFC at the same volume and coordinates used previously for the fiber photometry recordings. D2-cre mice received a unilateral injection of a mix of AAV-DJ-hSyn1-DIO-GCaMP8m and AAV9-Tet-On-D2R-IL3 or AAV9-Tet-On-D2R-IL3-scr. The optic fiber implantation procedure was the same as described previously.

### Doxycycline induction of interfering peptide expression

Peptide expression induction was achieved through the concomitant I.P. injection of Doxycycline solution (9-Tert-Butyl-Doxycycline HCL 10 mg/kg, DMSO 5%, Tween 20 5%, in saline 0.9%) for 5 days as well as continuous supply of Doxycycline in drinking water until sacrifice (Doxycycline Hyclate 2 g/L, Sigma; 9-Tert-Butyl-Doxycycline HCL 80 mg/L, TebuBio ; Sucrose 1%, Sigma ; pH = 4.5 ; Evian). Peptide expression was maintained over the whole course of behavioral experiments, through the I.P. injection of the same doxycycline solution every two weeks.

### Fiber photometry setup

Two RWD R820 fiber photometry setups were used. A 470nm LED light stimulation coupled to a 405nm isobestic stimulation was delivered through a fiber-optic patch cord (O.D. 1.25 mm, Ceramic Ferrule, 400 µm core diameter, 0.5 NA; R-FC-L-N5-400-L1, RWD). Intensities were set to reach a 100 µW power for 470 nm LED and 40 µW for 405 nm isobestic LED. Activity was recorded at 40 Hz.

### Behavioral Procedures

Animals were food deprived and maintained at 85-90% of their baseline *ad libitum* weight. Mice were placed in sound and light isolated operant chambers (40x30x36 cm, IMETRONIC) equipped with two retractable levers along with food delivery systems at the center. For fiber photometry experiments, diluted sweet milk (17.5%, Nestlé) was delivered in a trough connected to a pump. For behavioral procedures, grain pellets and purified milk pellets (Bio-Serv / Phymep, 20 mg, isocaloric) were distributed through a food dispenser. Reward delivery was associated to the illumination of a LED located above the associated reinforced lever together with the emission of a sound stimulus (65 dB, 3000 Hz, 200 ms).

### Operant protocols for action-outcome specific associations

#### Magazine Training

Animals were trained to collect reward pellets for 3 days in two different sessions (30 minutes, 13 rewards, daily alternation of the Grain-Milk order). Grain or milk reward pellets were randomly delivered throughout the session simultaneously to the light and sound cues.

#### Instrumental Conditioning

There were 2 sessions per day during which mice had to press on the lever to receive the associated reward pellet. At first, mice were trained with a Fixed Ratio 1 for 4 days on average. Similarly to the magazine training, the light and sound cues were activated at lever pressing. Once the animals reached the criterion, they were trained in a random ratio (RR), RR5 (P_reward_ = 0.2 ; 2 days), RR10 (P_reward_ = 0.1 ; 2 days), RR20 (P_reward_ = 0.05 ; 4 days). Sessions stopped once the 20 pellets criterion was reached or after 30 minutes.

#### Initial general instrumental conditioning and specific action – outcome association

This protocol was used to evaluate the implication of dopaminergic projections in the specific association between action and outcomes. There were 2 sessions per day during which DAT-cre mice expressing cre-dependent inhibitory DREADD (Gi) or mCherry in the VTA had to press on the lever to receive one reward pellet (Grain or milk pellets randomized between animals). At first, mice were trained with a Fixed Ratio 1, where one pellet was delivered following lever pressing. Once the animals reached the criterion of 20 pellets in less than 10 minutes (4 days), they were trained in a random ratio (RR), RR5 (P_reward_ = 0.2 ; 2 days), RR10 (P_reward_ = 0.1 ; 2 days), RR20 (P_reward_ = 0.05 ; 4 days). Sessions stopped once the 20 pellets criterion was reached or after 30 minutes. Following the RR20 training, animals were trained for 8 sessions with the specific association Action 1 → Outcome 1 and Action 2 → Outcome 2 (A1→O1, A2→O2) under CNO injection, to inhibit VTA dopaminergic activity.

#### Sensory-Specific Satiety Outcome Devaluation

Animals were given *ad libitum* access to grain or milk reward pellets for 80 minutes in individual cages with bedding prior to the test, the food was weighed before and after feeding to evaluate consumption. They were then placed in the operant chamber for a choice test under extinction for 15 minutes with both levers available. Right after the test, animals were given access to both rewards successively for 15 minutes each, in individual cages with bedding and the food was weighed before and after feeding to evaluate consumption. At least 2 outcome devaluation procedures were performed where the prefeeding pellets were counterbalanced between sessions, separated by 4 days of training in RR20. Lever presses on the valued vs. devalued lever were compared and a devaluation ratio corresponding to 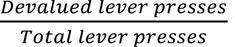 was computed to account for the potential lever pressing variability within each group, giving an index ranging from 0 (devaluation) to 1 (devaluation) that was compared to 0.5 (same amount of lever presses on both levers).

#### Reversal of outcome identities

Following the last outcome devaluation, animals were trained in RR20 for 3 days before switching to Reversal Learning. The association between right or left lever and the corresponding reward pellet was reversed. Animals were trained with this new protocol for 4 days before being subjected to a new Outcome Devaluation procedure. A minimal threshold value of 50 lever presses in total was applied to include animals in the analysis after each procedure. Hence, one DAT-cre animal (control mCherry, Figure 1) was removed from the devaluation procedure following specific acquisition of action – outcome association; one DAT-cre animal was removed (cre-, Figure 1) from the devaluation procedure following reversal learning; three WT animals (2 control peptides; 1 D2R-IL3 peptide, Figure 5) were removed from the devaluation procedure following reversal learning.

#### CNO treatment

For chemogenetic manipulations, CNO (2 mg/kg) was injected I.P. 30 minutes prior to any test. For specific A-O associations or for reversal learning, CNO was injected before the training sessions but not during the outcome devaluation procedure. For the manipulation during outcome devaluation, the CNO was injected 30 minutes before the beginning of the extinction test (end of the prefeeding).

#### Operant protocols used for fiber photometry recordings

Mice were first trained in Pavlovian conditioning for 10-12 days (15-20 minutes depending on cohorts) following a protocol used in^68^. During the session, the house LED was on. The emission of a light and sound stimulus (CS) predicted the delivery of a drop of diluted sweet milk (US). The CS was on for 10 seconds and the US occurred 3 seconds after the CS onset. Once the trough was empty and the CS was off, a new trial begun. The inter-trial interval (ITI) lasted 30 seconds.

Animals were then trained in Fixed Ratio 1 (FR1, 15-20 minutes depending on cohorts), during which both levers were out but only one was reinforced. Pressing on the non-reinforced lever led to no consequence. When the reinforced lever was pressed, both levers retracted and the CS started with the same CS-US sequence as pavlovian training. Levers were out again at the beginning of the new trial, after an ITI of 10 seconds. After FR1 training (8 sessions), animals were trained in reversal learning (8 sessions, 15-20 minutes depending on cohorts), in which reinforced and non-reinforced lever were switched. A lever preference ratio was computed 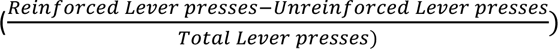, ranging from -1 (preference of the unreinforced lever) to 1 (preference of the reinforced lever).

#### Mouse tissue preparation

Mice received analgesia 30 min prior to the procedure through the subcutaneous injection of Buprenorphine (BUPRECARE, 0.1 mg/kg) before being deeply anesthetized with an I.P. injection of sodic pentobarbital (Euthasol, 300 mg/kg) and before being perfused with an ice cold 4% paraformaldehyde (PFA) solution in PBS 1X. Brains were postfixed overnight in a 4% PFA solution, then transferred to PBS 1X, before further use. Brains were sliced (40 µm thickness for viral expression verification; 30 µm for PLA quantification) using a vibratome (Leica) and slices were collected in a cryoprotectant solution containing 30% (v/v) ethylene glycol, 30% (v/v) glycerol, and 0.1 M PBS and stocked at -20°C until further use.

#### Histology and image acquisition

For interfering peptides and sensors expressions, slices were mounted with Vectashield Hardset mounting medium with DAPI (H-1500, VectorLab, Eurobio Scientific), before being scanned at 20X with a NanoZoomer slide scanner (Hamamatsu). The infections were then assessed in the targeted structures according to mCherry, RFP or GFP expression, respectively. For the chemogenetic manipulations and expression of interfering peptides, animals were removed if the viral expression was too restricted in the experimental groups, but all control animals were kept (Heteromer blockade, 1 GluN1-C1 and 1 D2R-IL3 removed). For fiber photometry experiments, animals were removed when photometry signal was weak, or if the fiber was misplaced (1 D1-cre animal (GluN1-C1 peptide) removed from GCaMP analysis and 1 WT animal (GRAB-DA2m) removed from cortical dopamine recording).

#### Proximity Ligation Assay

Male WT mice (10 weeks, n = 4 for D1/NMDA and n = 6 for D2/NMDA) were perfused with PFA and processed according to the procedure described. The PLA protocol is adapted from the one of the provider (Duolink - Merck). 3 slices (anterior PFC, AP = +2.22 ; posterior PFC, AP = +1.78 ; anterior striatum, AP = +1.10,^69^) per animal were used. Slices were sorted on ice and washed in TBS 1X (3 x 10 min). For D1/NMDA, slices were processed in 48-well plate. For D2/NMDA, slices were mounted on slides, left to dry and encircled with the hydrophobic PAP pen (Thermofischer, Epredia™ SuperFrost Ultra Plus™ GOLD, 11976299; ImmEdge Hydrophobic Barrier PAP pen (H-4000)). All washing and incubation steps were done under stirring. Except for TBS, FBS, Triton X-100, and primary antibodies, all reagents were provided in the Duolink Brightfield Detection kit (Merck). They were then quenched with hydrogen peroxide (D1/NMDA, floating slices: 5min ; D2/NMDA, mounted slices: 30 min with H2O2 (sigma) diluted into TBS1X (6.6 mL in 200 mL TBS 1X) protected from light) to suppress endogenous peroxidase activity, then washed with Buffer A (D1/NMDA, floating slices; Buffer A from the kit), or TBS-T (D2/NMDA, mounted slices, TBS-Triton X-100 0,2%) for 3 x 5 min. Slices were then incubated with blocking/permeabilization solution (D1/NMDA, floating slices: 3% FBS, 0.2% Triton X-100, TBS 1X for 2 hours; D2/NMDA, mounted slices: solution from the kit for 1 hour) at room temperature (RT), before being incubated with primary antibodies (D1/NMDA : Goat polyclonal anti-D1R 1/200 D1R-Go-Af 1000, Frontier Institute ; rabbit polyclonal anti-GluN1 1/200 ab17345, Abcam; D2/NMDA : Rabbit polyclonal anti-D2R 1/150 Millipore ABN462 ; Mouse monoclonal anti-NMDAR2B NT 1/150 Millipore MAB5782 ; in antibody diluent solution ; negative control: anti-GluN1 only ; anti-D2R only) overnight at 4°C. Slices were then washed with Buffer A (D1/NMDA, floating slices, 4 x 30 min) or TBS-T (D2/NMDA, mounted slices, 3 x 10 min), and incubated with secondary PLA probes (D1/NMDA : Duolink Anti-Goat Minus probe 1/5, Duolink anti-Rabbit Plus probe 1/5; D2/NMDA : Duolink Anti-Rabbit Plus probe 1/5, Duolink anti-Mouse Minus probe 1/5; positive control: Duolink anti-Mouse Minus probe 1/5, Duolink anti-Mouse Plus probe 1/5, in antibody diluent solution) for 1h (D2/NMDA, mounted slices) or 2h (D1/NMDA, floating slices) at 37°C. Slices were washed in buffer A (3 x 5 min) and then incubated in the ligation solution (ligation buffer 1/5, ligase 1/40, in mQ H_2_O) for 45 min at 37°C. Slices were washed in buffer A (3 x 5 min) and then incubated in the amplification solution (amplification buffer 1/5, polymerase 1/80, in mQ H_2_O) for 2h at 37°C. Slices were washed in buffer A (3 x 5 min) and then incubated in the detection solution (HRP probes 1/5, in mQ H_2_O) for 1h at RT. Slices were washed in buffer A (3 x 5 min) and incubated in the substrate development solution (3,3’-Diaminobenzidine, DAB substrates : A 1/70, B 1/100, C 1/100, D 1/50 in mQ H_2_O) for 10min at RT. For D1/NMDA, the floating slices were washed in buffer A (2 x 5 min) and mounted on slides before drying overnight. For D2/NMDA, mounted slices were washed in mQ H_2_O (2 x 5 min). Nuclear staining was then performed on the slides for 2 min and slides were rinsed with mQ H_2_O for 10 min. Slides were then dehydrated (95% EtOH 2x2min, 100% EtOH 2x2min, Xylene 2x5min) and mounted with a coverslip using DPX / Permount mounting medium.

#### Quantification of PLA signal

Slides were scanned at 40X over 8-21 stacks (step = 0.85-1µm) using a slide scanner (Nanozoomer Hamamatsu; Zeiss Axioscan). Nine subregions of the PFC and striatum were determined according to ^69^ (ACC, PL, IL, MO, DMS, DLS, Core, medShell, latShell). Six areas (when applicable) per region per hemisphere of 150 µm X 150 µm were determined using QuPath. Acquisitions were then processed in ImageJ (FIJI, National Institutes of Health (NIH)) using a custom macro to determine the best focus of each stack (Based on Extended Depth Field plugin from https://github.com/Biomedical-Imaging-Group/EDF-Extended-Depth-of-Field), and a custom macro adapted from the “Find Maxima” macro (Prominence = 14) to detect the PLA dots. Quantification was expressed in PLA points x 10^3^ / mm^2^.

#### Fiber photometry analysis

Fiber photometry recordings of dopamine release and D1- or D2-dopaminceptive neurons activity in the mPFC were analyzed using a MATLAB custom script. The isobestic signal was fitted (linear fitting) to the 470 nm signal and ΔF/F was calculated as (F 470 nm – F 405 nm fitted)/F 405 nm fitted for each time point. The z-score around each behavioral event was then calculated using a baseline of 1.5sec (from -2.5 sec to -1 sec before the event) as 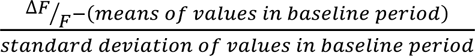. Area Under the Curve (AUC) were calculated in MATLAB using the *trapz* function. Peak z-score and minimum z-score were calculated as the maximum and minimum value in the specific interval, respectively.

#### Statistical analysis

For statistical analysis, GraphPad Prism (version 10.0.1, GraphPad Software Inc.) was used. All data are displayed as means +/-SEM. Data sets were tested for normality and subsequent test were performed accordingly. Unpaired two-tailed Student’s t-test or Mann-Whitney test were used for the comparison of two independent groups. For the analysis of more than two groups with one factor, one-way analysis of variance (ANOVA, parametric) or Kruskall-Wallis test (non-parametric), were used. When two factors were present, two-way repeated-measures ANOVA were performed. When applicable, the ANOVA were followed by Tukey’s multiple comparison (when comparing every mean with every other mean) or Šídák’s multiple comparison (when comparing a set of means), and the Kruskall-Wallis test by Dunn’s multiple comparison test. One-sample t-test were used to compare group means to a fixed value (*0.5* for the devaluation ratio; *0* for fiber photometry analysis of change in AUC and max or min z-score from baseline). Planned orthogonal contrasts were performed to analyze the data from choice tests under extinction in the outcome devaluation procedure, using PSY software (Kevin Bird, Dusan Hadzi-Pavlovic and Andrew Isaac, School of Psychology and School of Psychiatry, University of New South Wales). These contrasts tested for the main group effect (Control group versus treatment group(s)), the main devaluation effect (valued lever vs. devalued lever) and the interaction. Simple effects were then performed accordingly. Statistical significance was * *p* <0.05, ** *p* < 0.01, *** *p* < 0.001, **** p < 0.0001.

