## Supplementary Material for "A role for prefrontal dopamine signaling in control of goal-directed actions"

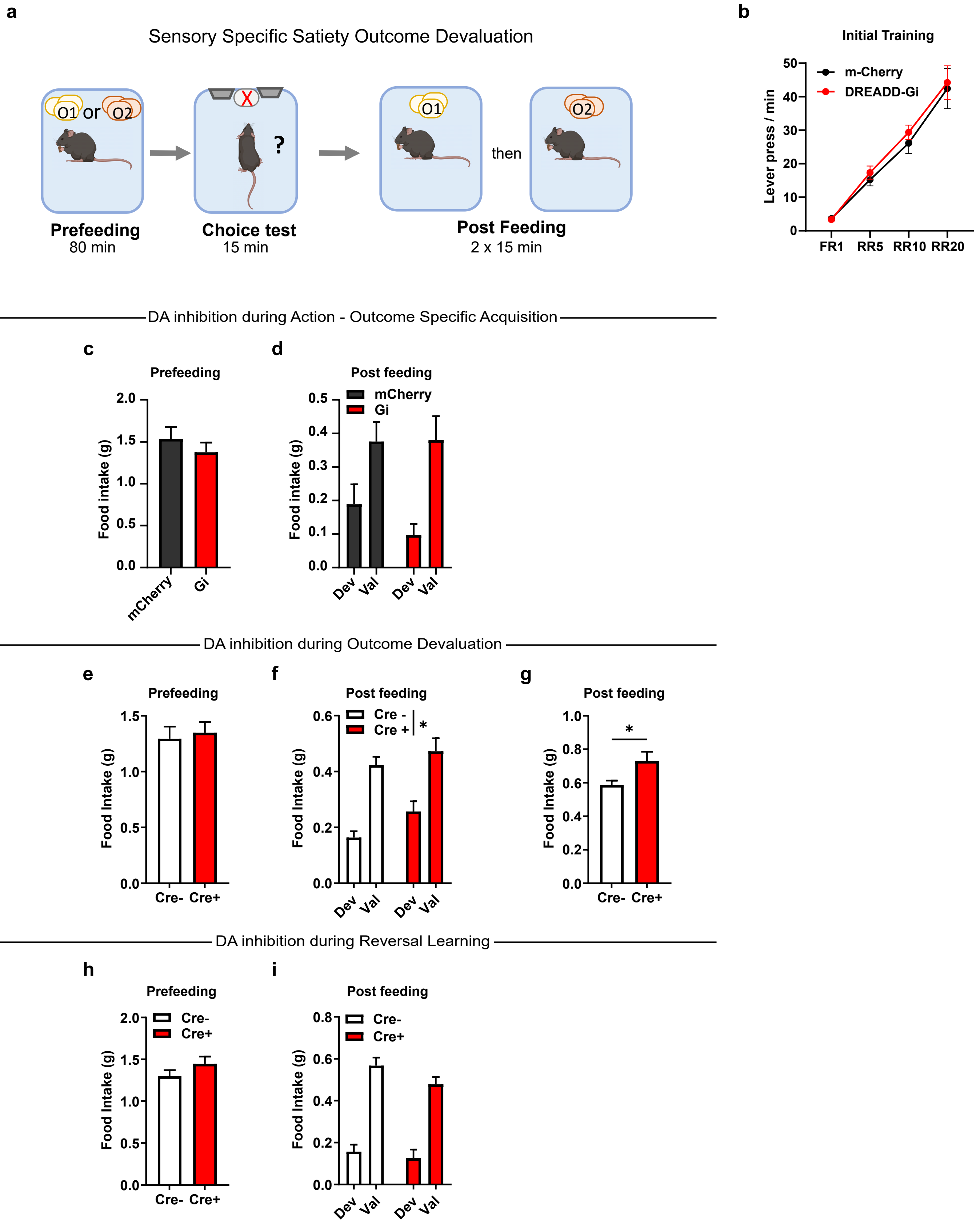

a
Sensory Specific Satiety Outcome Devaluation
O1 or O2
?
O1
then
O2
Prefeeding
80 min
Choice test
15 min
Post Feeding
2 x 15 min
b
Initial Training
m-Cherry
DREADD-Gi
Lever press / min
FR1 RR5 RR10 RR20
DA inhibition during Action - Outcome Specific Acquisition
c
Prefeeding
Food intake (g)
mCherry Gi
d
Post feeding
mCherry
Gi
Food intake (g)
Dev Val Dev Val
DA inhibition during Outcome Devaluation
e
Prefeeding
Food Intake (g)
Cre- Cre+
f
Post feeding
Cre -
Cre +
*
Food Intake (g)
Dev Val Dev Val
g
Post feeding
*
Food Intake (g)
Cre- Cre+
DA inhibition during Reversal Learning
h
Prefeeding
Cre-
Cre+
Food Intake (g)
Cre- Cre+
i
Post feeding
Cre-
Cre+
Food Intake (g)
Dev Val Dev Val

**<u>Supplementary Figure 1</u> : Inhibiting the VTA dopaminergic projections does not alter food devaluation.**

**a**, Sensory specific satiety outcome devaluation procedure schematic representation, prefeeding is followed by the choice test, then by a postfeeding test. **b**, Lever pressing frequency across operant schedule in DAT-cre mice expressing mCherry (n=10) or DREADD-Gi (n=10) before CNO administration. **c-d** Outcome devaluation procedure following specific action-outcome training, mCherry (n=9), DREADD-Gi (n=10) **c**, Food intake during prefeeding (g); **d**, Consumption of each reward following the outcome devaluation test. **e-g** Outcome devaluation procedure, Cre- (n=11), Cre+ (n=11) **e**, Food intake during prefeeding (g); **f**, Consumption of each reward following the outcome devaluation test (Two RM ANOVA, Group effect F (1, 20) = 5.299, p=0.0322); **g**, Total food following the outcome devaluation test (Welch's test, t=2.302, df=14,48, p=0.0367). **h-i** Outcome devaluation procedure following reversal learning, Cre- (n=10), Cre+ (n=11) **h**, Food intake during prefeeding (g); **i**, Consumption of each reward following the outcome devaluation test. All datasets represent mean values +/- s.e.m. *$p$<0.05. Dev, devalued; FR, fixed, ratio; O, outcome; RR, random ratio; Val, valued. Detailed statistics are shown in Supplementary Table 1.

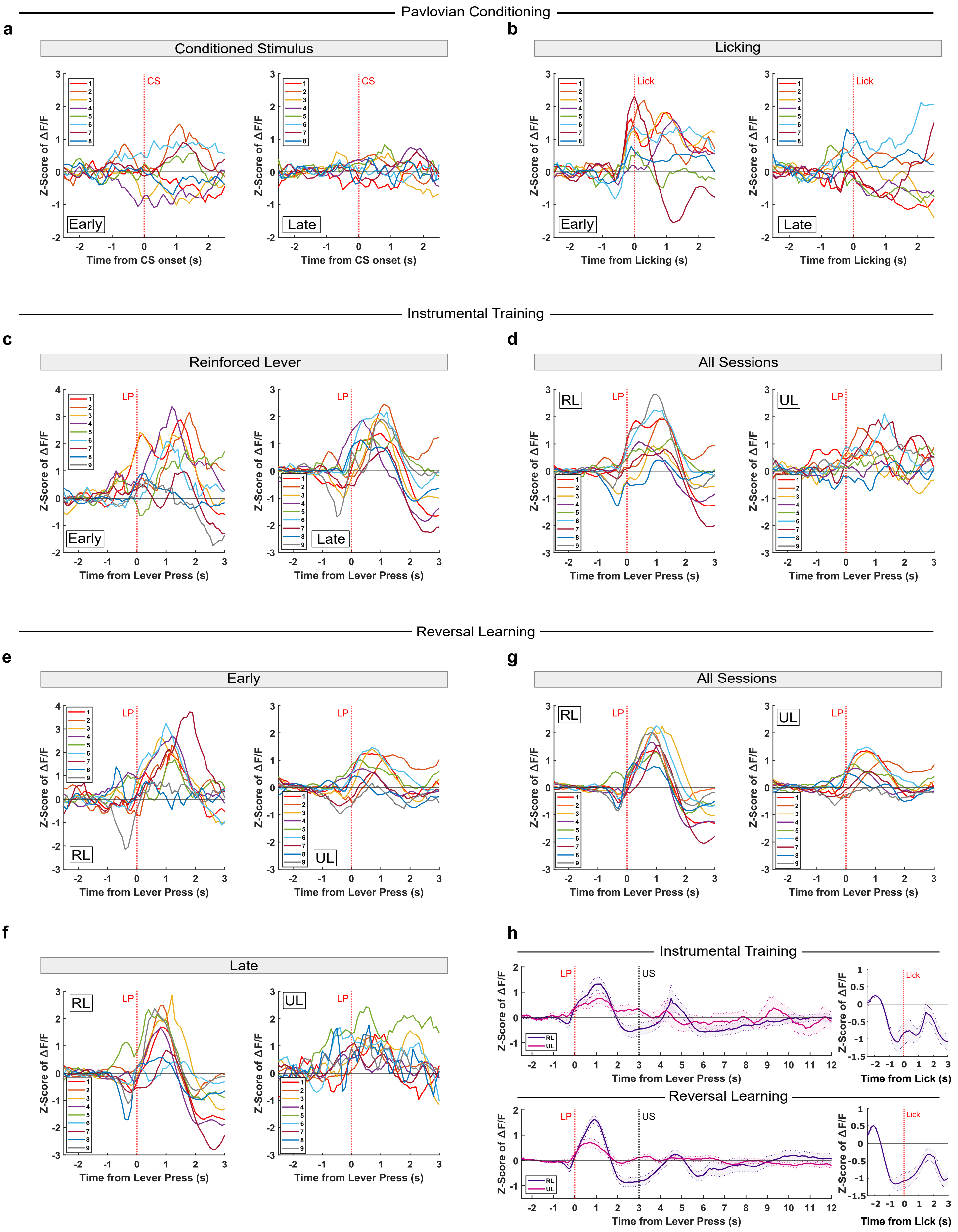

Pavlovian Conditioning
a
Conditioned Stimulus
b
Licking
Early
Late
CS
Lick
Z-Score of ΔF/F
Time from CS onset (s)
Time from Licking (s)
Instrumental Training
c
Reinforced Lever
d
All Sessions
RL
UL
LP
Time from Lever Press (s)
Reversal Learning
e
Early
g
All Sessions
f
Late
h
Instrumental Training
Reversal Learning
US
Time from Lick (s)

**<u>Supplementary Figure 2</u> : Individual traces of mPFC dopamine dynamics averaged for each animal across pavlovian and instrumental tasks**

**a-b** Pavlovian conditioning (n=8); **a**, Z-Score of ΔF/F at CS onset; **(left)** individual traces in early sessions, **(right)** individual traces in late sessions; **b**, Z-Score of ΔF/F at lick onset (n=8); **(left)** individual traces in early sessions, **(right)** individual traces in late sessions. **c-d** Instrumental training (n=9); **c**, Z-Score of ΔF/F at lever pressing on the reinforced lever; **(left)** individual traces in early sessions, **(right)** individual traces in late sessions; **d**, Z-Score of ΔF/F at lever pressing over all sessions, **(left)** individual traces for the reinforced lever, **(right)** individual traces for the unreinforced lever. **e-g** Reversal Learning (n=9); **e**, Z-Score of ΔF/F at lever pressing during early sessions, **(left)** individual traces for the reinforced lever, **(right)** individual traces for the unreinforced lever ; **f**, Z-Score of ΔF/F at lever pressing during late sessions, **(left)** individual traces for the reinforced lever, **(right)** individual traces for the unreinforced lever ; **g**, Z-Score of ΔF/F at lever pressing over all sessions, **(left)** individual traces for the reinforced lever, **(right)** individual traces for the unreinforced lever. **h**, **top**, instrumental training (n=9); **(left)** Z-Score of ΔF/F over a trial, from lever press onset on the reinforced or unreinforced lever, **(right)** Z-Score of ΔF/F at lick onset; **bottom**, reversal learning (n=9); **(left)** Z-Score of ΔF/F over a trial, from lever press onset on the reinforced or unreinforced lever, **(right)** Z-Score of ΔF/F at lick onset. All datasets represent mean values +/- s.e.m. *$p<0.05$. CS, Conditioned stimulus; LP, lever press; RL, reinforced lever, UL, unreinforced lever, US, Unconditioned stimulus. Detailed statistics are shown in Supplementary Table 1.

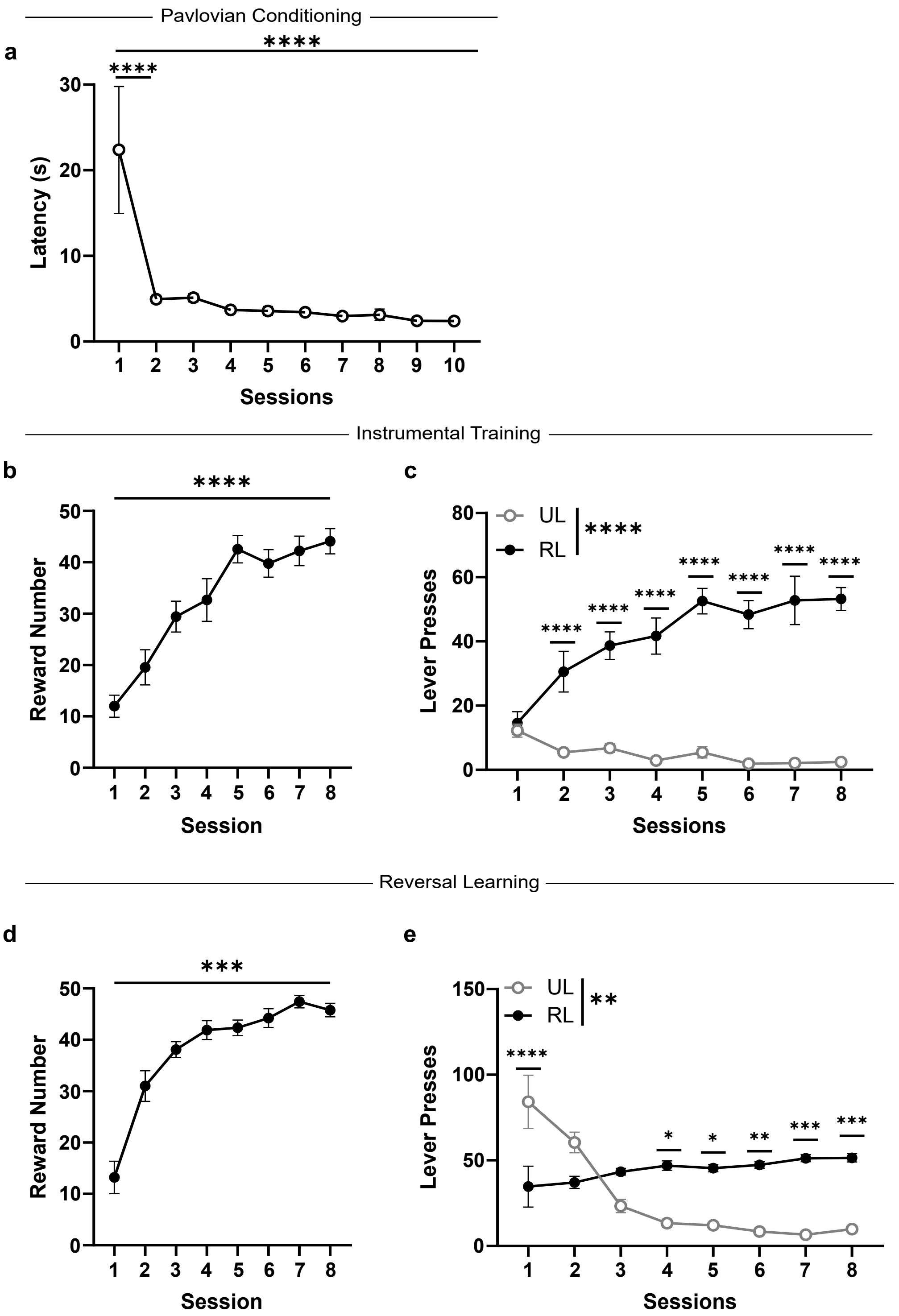

Pavlovian Conditioning
a
****
****
Latency (s)
0
10
20
30
1 2 3 4 5 6 7 8 9 10
Sessions
Instrumental Training
b
****
Reward Number
0
10
20
30
40
50
1 2 3 4 5 6 7 8
Session
c
UL
RL
****
****
****
****
****
****
****
****
Lever Presses
0
20
40
60
80
1 2 3 4 5 6 7 8
Sessions
Reversal Learning
d
***
Reward Number
0
10
20
30
40
50
1 2 3 4 5 6 7 8
Session
e
UL
RL
**
****
*
*
**
***
***
Lever Presses
0
50
100
150
1 2 3 4 5 6 7 8
Sessions

**<u>Supplementary Figure 3</u> : Behavioral performance during recording of dopamine dynamics in the mPFC**

**a**, Latency from reward delivery to reward consumption (s) during pavlovian conditioning (n=9; RM One-way ANOVA F(9, 72) = 6.385, p<0.0001, Tukey's multiple comparisons test, 1 vs. 2 p<0.0001, 1 vs. 10 p<0,0001). **b-c** Instrumental Training; **b**, Number of rewards obtained over training sessions (n=9; RM One-way ANOVA F(7, 56) = 63.70, p<0.0001, Dunnett's multiple comparisons test, 1 vs. 8 p<0.0001); **c**, Lever presses on reinforced versus unreinforced levers over training sessions (n=9; RM Two-way ANOVA, Lever Effect F (1, 8) = 66.81, p<0.0001, Session x Lever Interaction F (7, 56) = 24.20, p<0.0001, Šídák's multiple comparisons test, UL – RL, 1 p=0.9957, 2 p<0.0001, 3 p<0.0001, 4 p<0.0001, 5 p<0.0001, 6 p<0.0001, 7 p<0.0001, 8 p<0.0001). **d-e**, Reversal Learning; **d**, Number of rewards obtained over training sessions (n=9; RM One-way ANOVA F (7, 56) = 47.14, p<0.0001, Dunnett's multiple comparisons test, 1 vs. 8 p<0.0001); **e**, Lever presses on reinforced versus unreinforced levers over training sessions (n=9; RM Two-way ANOVA, Lever Effect F(1, 8) = 19.07, p=0.0024, Session x Lever Interaction F(7, 56)= 12.08, p<0.0001, Šídák's multiple comparisons test, UL – RL, 1 p<0.0001, 2 p=0.1716, 3 p=0.3386, 4 p=0.0114, 5 p=0.0121, 6 p=0.0022, 7 p=0.0003, 8 p=0.0009). All datasets represent mean values +/- s.e.m. *$p$<0.05, **$p$<0.01, ***$p$<0.001, ****$p$<0.0001. RL, reinforced lever, UL, unreinforced lever. Detailed statistics are shown in Supplementary Table 1.

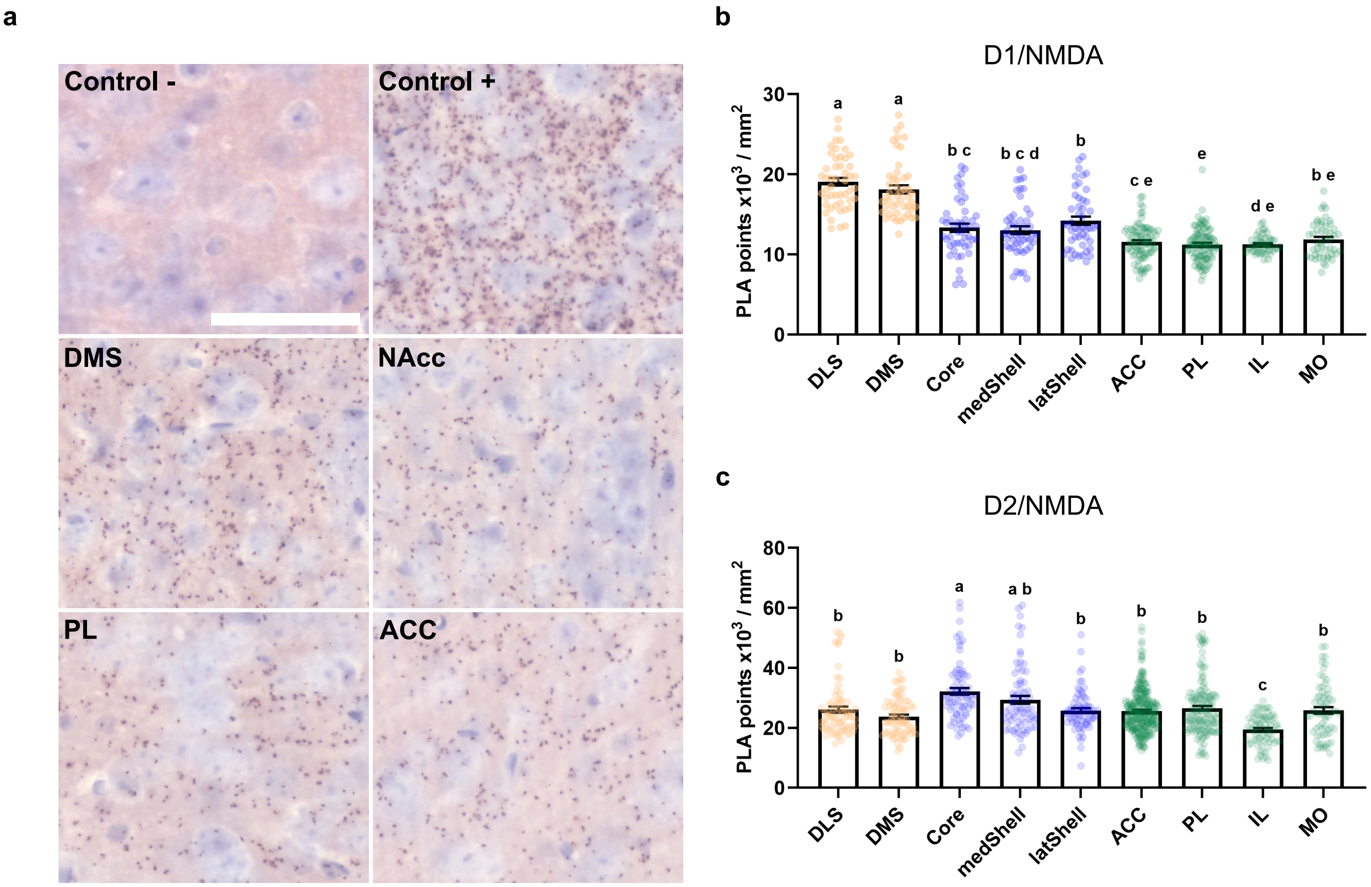
a
Control -
Control +
DMS
NAcc
PL
ACC
b
D1/NMDA
PLA points x10^3 / mm^2
0
10
20
30
a
a
b c
b c d
b
c e
e
d e
b e
DLS
DMS
Core
medShell
latShell
ACC
PL
IL
MO
c
D2/NMDA
PLA points x10^3 / mm^2
0
20
40
60
80
b
b
a
a b
b
b
b
c
b
DLS
DMS
Core
medShell
latShell
ACC
PL
IL
MO

**Supplementary Figure 4 : In situ quantification of striatal and medial prefrontal D1/NMDA and D2/NMDA complexes through proximity ligation assay**

**a**, Example of D2/NMDA detection using proximity ligation assay (PLA) in different subregions, from top left to bottom right: negative control (anti-D2R only), positive control (NMDAR-2B single recognition), dorsomedial striatum, nucleus accumbens core, prelimbic cortex, anterior cingulate cortex; scale bar = 50 µm. **b**, D1/NMDA quantification per striatal and cortical subregion (PLA points x $10^3$ / $mm^2$) ; (Kruskal-Wallis test, H(8) = 218.4, p<0.0001, Dunn's multiple comparisons test). **c**, D2/NMDA quantification per striatal and cortical subregion (PLA points x $10^3$ / $mm^2$) ; (Kruskal-Wallis test, H(8) = 99.49, p<0.0001, Dunn's multiple comparisons test). Detailed statistics are shown in Supplementary Table 1. ACC, anterior cingulate cortex ; Core, nucleus accumbens core ; DLS, dorsolateral striatum ; DMS, dorsomedial striatum ; IL, infralimbic cortex ; latShell, nucleus accumbens lateral shell ; medShell, nucleus accumbens medial shell ; MO, medial orbitofrontal cortex ; PL, prelimbic cortex ; PLA, proximity ligation assay.

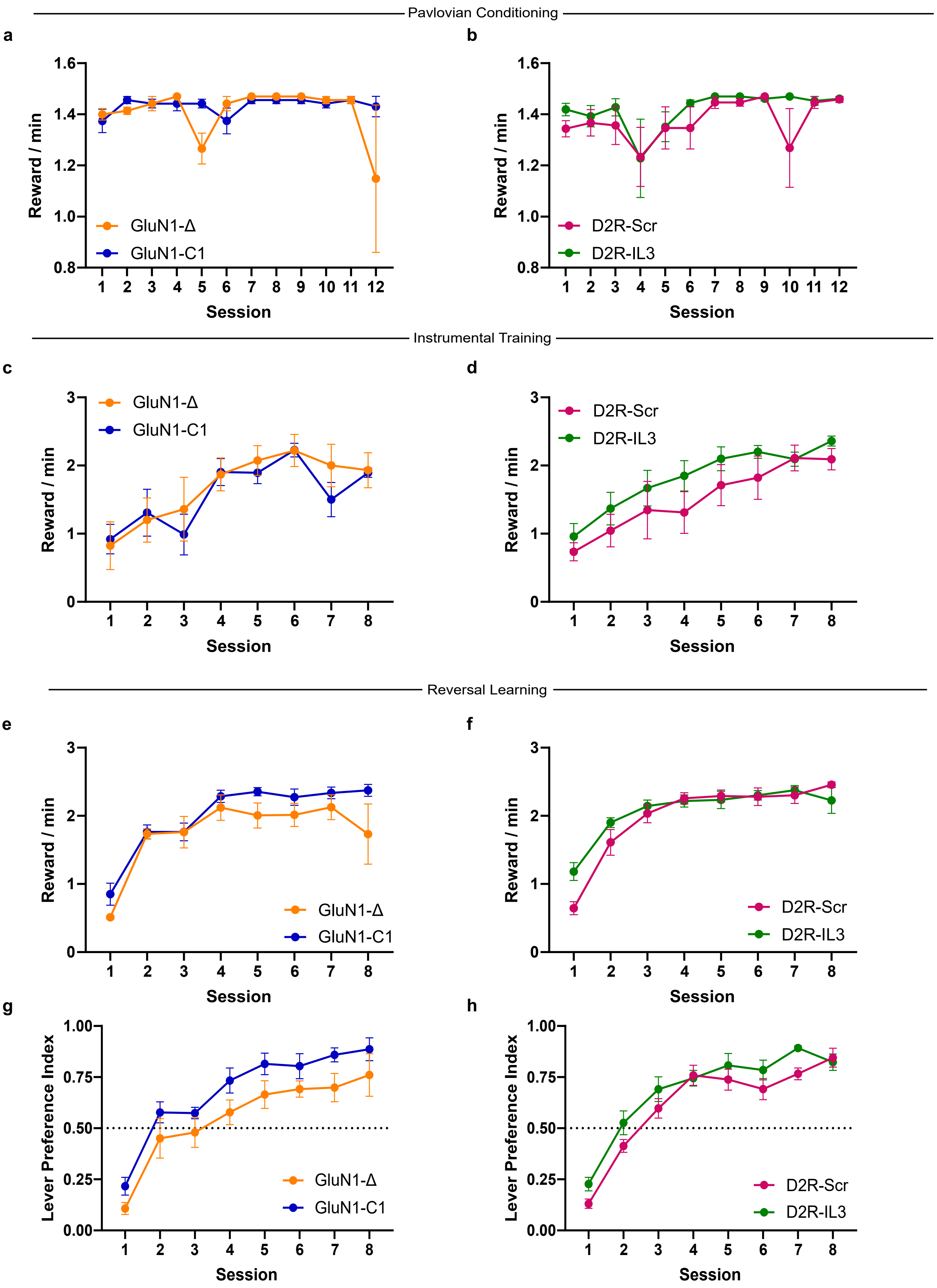

Pavlovian Conditioning
a
b
Reward / min
Session
GluN1-Δ
GluN1-C1
D2R-Scr
D2R-IL3
Instrumental Training
c
d
Reward / min
Session
GluN1-Δ
GluN1-C1
D2R-Scr
D2R-IL3
Reversal Learning
e
f
Reward / min
Session
GluN1-Δ
GluN1-C1
D2R-Scr
D2R-IL3
g
h
Lever Preference Index
Session
GluN1-Δ
GluN1-C1
D2R-Scr
D2R-IL3

<u>Supplementary Figure 5</u> **: Behavioral performance of mice expressing a control peptide or an interfering peptide blocking either D1/NMDA or D2/NMDA unilaterally in the mPFC during calcium dynamics recording of mPFC D1R+ or D2R+ dopaminoceptive neurons**

**a-b** Reward number per minute over pavlovian training sessions for, **a**, GluN1-Δ (n=5) or GluN1-C1 (n=5) groups, **b**, D2R-Scr (n=6) or D2R-IL3 (n=8) groups. **c-d** Reward number per minute over instrumental training sessions for, **c**, GluN1-Δ (n=5) or GluN1-C1 (n=5) groups, **d**, D2R-Scr (n=6) or D2R-IL3 (n=8) groups. **e-f** Reward number per minute over Reversal learning training sessions for, **e**, GluN1-Δ (n=5) or GluN1-C1 (n=5) groups, **f**, D2R-Scr (n=6) or D2R-IL3 (n=8) groups. **g-h** Lever preference index over Reversal learning training sessions for, **g**, GluN1-Δ (n=5) or GluN1-C1 (n=5) groups, **h**, D2R-Scr (n=6) or D2R-IL3 (n=8) groups. All datasets represent mean values +/- s.e.m. Detailed statistics are shown in Supplementary Table 1.

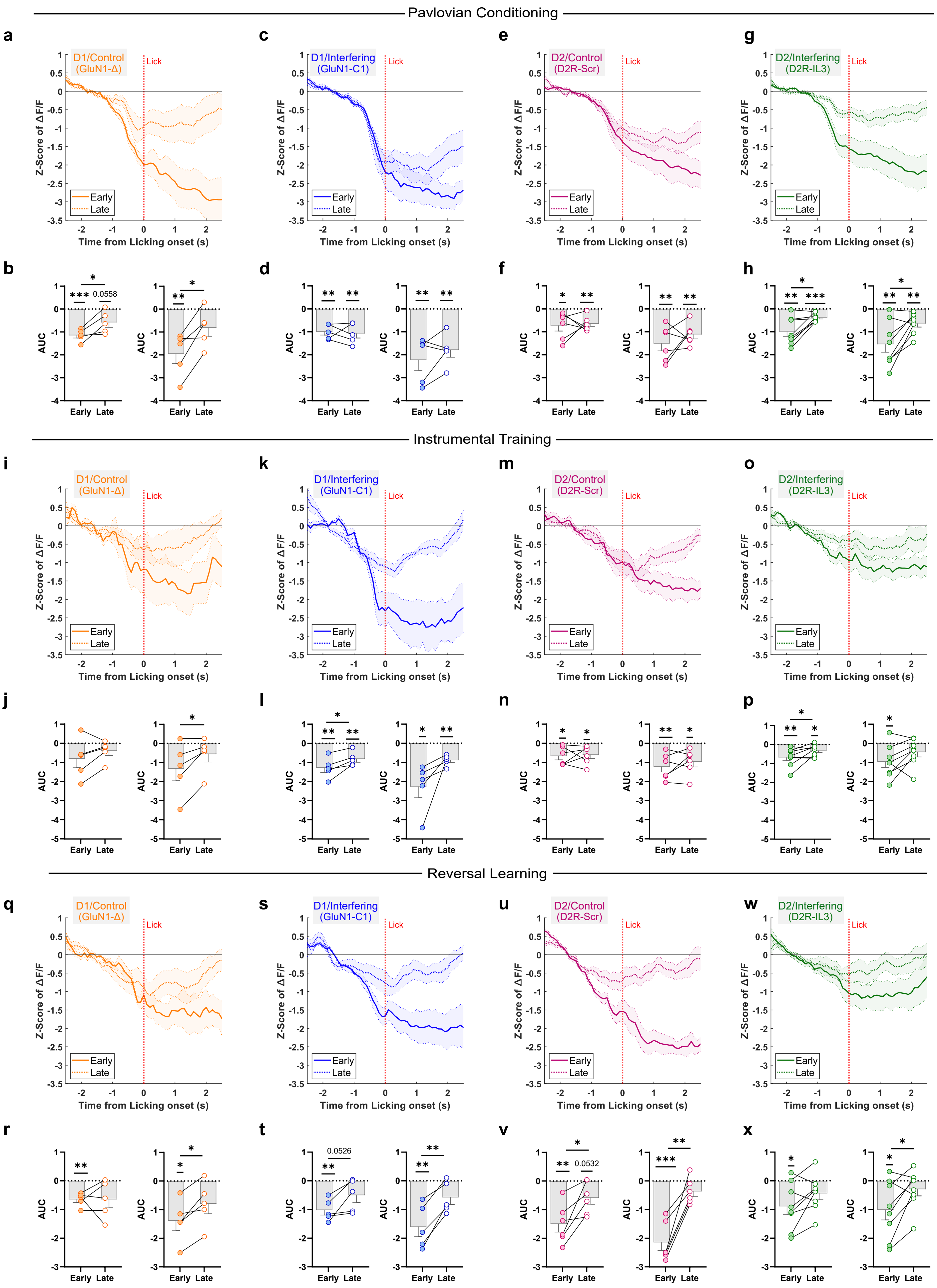

Pavlovian Conditioning
a
D1/Control (GluN1-Δ)
c
D1/Interfering (GluN1-C1)
e
D2/Control (D2R-Scr)
g
D2/Interfering (D2R-IL3)
Lick
Early
Late
Z-Score of ΔF/F
Time from Licking onset (s)
b
d
f
h
AUC
Early Late
0.0558
Instrumental Training
i
D1/Control (GluN1-Δ)
k
D1/Interfering (GluN1-C1)
m
D2/Control (D2R-Scr)
o
D2/Interfering (D2R-IL3)
j
l
n
p
Reversal Learning
q
D1/Control (GluN1-Δ)
s
D1/Interfering (GluN1-C1)
u
D2/Control (D2R-Scr)
w
D2/Interfering (D2R-IL3)
r
t
0.0526
v
0.0532
x

<u>Supplementary Figure 6</u> **: Recordings of calcium dynamics of mPFC D1R+ or D2R+ neurons at licking onset under control or interfering peptide condition during pavlovian conditioning, instrumental training and reversal learning**

**a-h**, Pavlovian conditioning. **a**, Z-Score of ΔF/F for D1R+ neurons, D1R+$_{Control}$ (*n* = 5), **b**, AUC (**left**) [-0.9s ; 0s] (One-sample t-test against 0, Early p = 0.0006, Late p = 0.0558, Two-tailed paired t-test , p = 0.0363) and (**right**) [0.1s ; 1s] (One-sample t-test against 0, Early p = 0.0097, Late p = 0.0211); **c**, Z-Score of ΔF/F for D1R+ neurons, D1R+$_{Interfering}$ (n=5), **d**, AUC (**left**) [-0.9s ; 0s] (One-sample t-test against 0, Early p = 0.0023, Late p = 0.0059) and (**right**) [0.1s ; 1s] (One-sample t-test against 0, Early p = 0.0075, Late p = 0.0046); **e**, Z-Score of ΔF/F for D2R+ neurons, D2R+$_{Control}$ (*n* = 6), **f**, AUC (**left**) [-0.9s ; 0s] (One-sample t-test against 0, Early p = 0.0311, Late p = 0.0043), (**right**) [0.1s ; 1s] (One-sample t-test against 0, Early p = 0.0049, Late p = 0.0024); **g**, Z-Score of ΔF/F for D2R+ neurons, D2R+$_{Interfering}$ (*n* = 8), **h**, AUC (**left**) [-0.9s ; 0s] (One-sample t-test against 0, Early p = 0.0021, Late p = 0.0007, Two-tailed paired t-test , p = 0.0113), (**right**) [0.1s ; 1s] (One-sample t-test against 0, Early p = 0.0029, Late p = 0.0049, Two-tailed paired t-test , p = 0.0166). **i-p**, Instrumental Training. **i**, Z-Score of ΔF/F for D1R+ neurons, D1R+$_{Control}$ (*n* = 5), **j**, AUC (**right**) [0.1s ; 1s] (Two-tailed paired t-test , p = 0.0430); **k**, Z-Score of ΔF/F for D1R+ neurons, D1R+$_{Interfering}$ (n=5), **l**, AUC (**left**) [-0.9s ; 0s] (One-sample t-test against 0, Early p = 0.0062, Late p = 0.0079, Two-tailed paired t-test , p = 0.0208) and (**right**) [0.1s ; 1s] (One-sample t-test against 0, Early p = 0.0159, Late p = 0.0023); **m**, Z-Score of ΔF/F for D2R+ neurons, D2R+$_{Control}$ (*n* = 6), **n**, AUC (**left**) [-0.9s ; 0s] (One-sample t-test against 0, Early p = 0.0187, Late p = 0.0247), (**right**) [0.1s ; 1s] (One-sample t-test against 0, Early p = 0.0071, Late p = 0.0169); **o**, Z-Score of ΔF/F for D2R+ neurons, D2R+$_{Interfering}$ (*n* = 8), **p**, AUC (**left**) [-0.9s ; 0s] (One-sample t-test against 0, Early p = 0.0050, Late p = 0.0171, Two-tailed paired t-test , p = 0.0493), (**right**) [0.1s ; 1s] (One-sample t-test against 0, Early p = 0.0155). **q-x**, Reversal Learning. **q**, Z-Score of ΔF/F for D1R+ neurons, D1R+$_{Control}$ (*n* = 5), **r**, AUC (**left**) [-0.9s ; 0s] (One-sample t-test against 0, Early p = 0.0037) and (**right**) [0.1s ; 1s] (One-sample t-test against 0, Early p = 0.0145, Two-tailed paired t-test, p = 0.0115); **s**, Z-Score of ΔF/F for D1R+ neurons, D1R+$_{Interfering}$ (n=5), **t**, AUC (**left**) [-0.9s ; 0s] (One-sample t-test against 0, Early p = 0.0039, Two-tailed paired t-test, p = 0.0526) and (**right**) [0.1s ; 1s] (One-sample t-test against 0, Early p = 0.0095, Two-tailed paired t-test, p = 0.0033); **u**, Z-Score of ΔF/F for D2R+ neurons, D2R+$_{Control}$ (*n* = 6), **v**, AUC (**left**) [-0.9s ; 0s] (One-sample t-test against 0, Early p = 0.0032, Late p = 0.0532, Two-tailed paired t-test, p = 0.0168), (**right**) [0.1s ; 1s] (One-sample t-test against 0, Early p = 0.0006, Two-tailed paired t-test, p = 0.0014); **w**, Z-Score of ΔF/F for D2R+ neurons, D2R+$_{Interfering}$ (*n* = 8), **x**, AUC (**left**) [-0.9s ; 0s] (One-sample t-test against 0, Early p = 0.0188), (**right**) [0.1s ; 1s] (One-sample t-test against 0, Early p = 0.0269, Two-tailed paired t-test, p = 0.0341). All datasets represent mean values +/- s.e.m. *$p$<0.05, **$p$<0.01, ***$p$<0.001. Detailed statistics are shown in Supplementary Table 1.

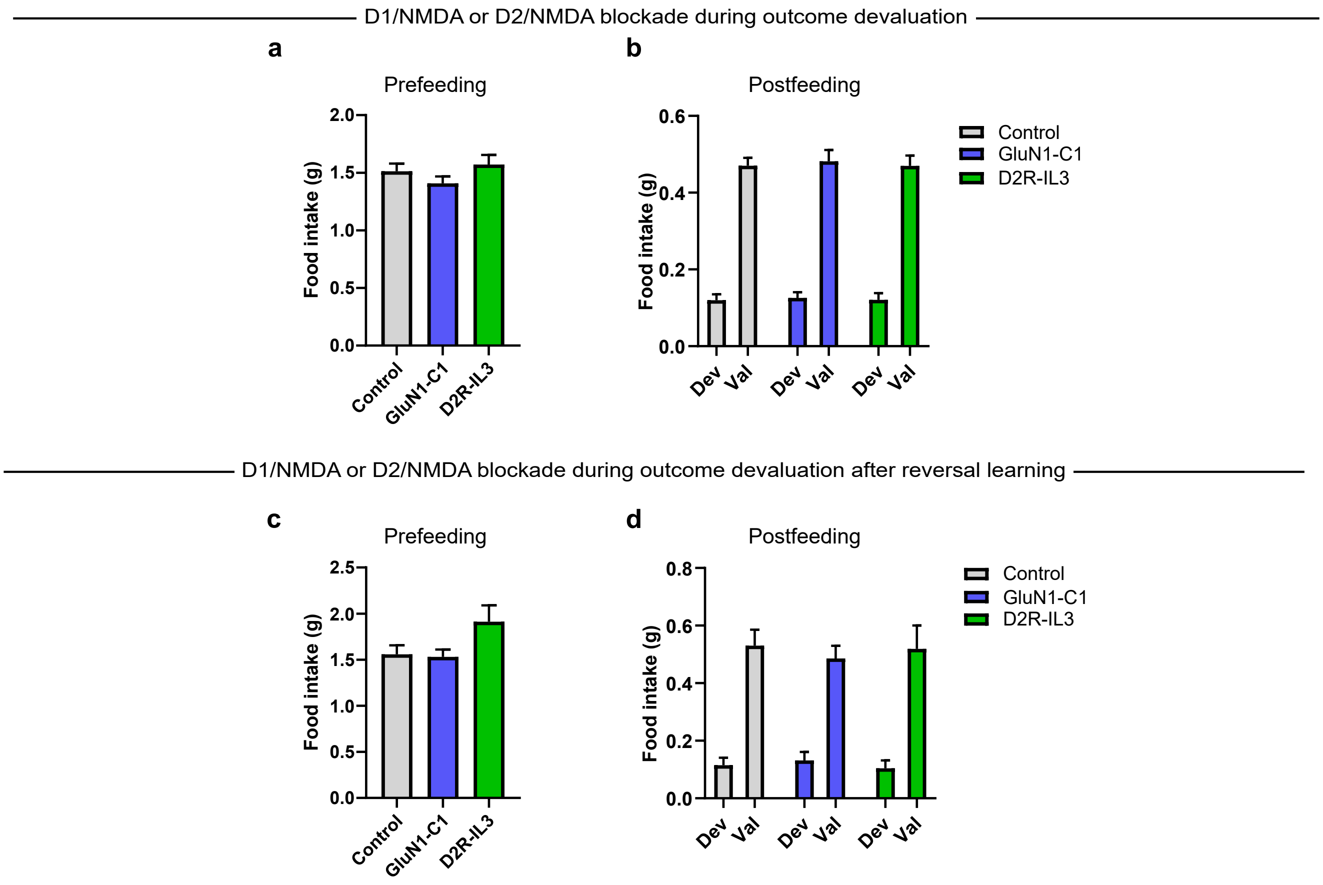
D1/NMDA or D2/NMDA blockade during outcome devaluation
a
Prefeeding
Food intake (g)
2.0
1.5
1.0
0.5
0.0
Control
GluN1-C1
D2R-IL3
b
Postfeeding
Food intake (g)
0.6
0.4
0.2
0.0
Dev
Val
Control
GluN1-C1
D2R-IL3
D1/NMDA or D2/NMDA blockade during outcome devaluation after reversal learning
c
Prefeeding
Food intake (g)
2.5
2.0
1.5
1.0
0.5
0.0
Control
GluN1-C1
D2R-IL3
d
Postfeeding
Food intake (g)
0.8
0.6
0.4
0.2
0.0
Dev
Val
Control
GluN1-C1
D2R-IL3

**Supplementary Figure 7 : Blocking mPFC D1/NMDA or D2/NMDA does not impact food consumption and devaluation**

**a-b** Outcome devaluation after initial learning of action-outcome associations; **a**, Food intake during prefeeding (g); **b**, Consumption of each reward following the outcome devaluation test. **c-d** Outcome devaluation after reversal learning; **c**, Food intake during prefeeding (g); **d**, Consumption of each reward following the outcome devaluation test. All datasets represent mean values +/- s.e.m. Dev, devalued; Val, valued. Detailed statistics are shown in Supplementary Table 1.

### Key Resource Table

| REAGENT or RESOURCE | SOURCE | IDENTIFIER |
|---|---|---|
| **Antibodies** | | |
| Goat polyclonal anti-D1R | Frontier Institute | D1R-Go-Af 1000 |
| Rabbit polyclonal anti-GluN1 | Abcam | ab17345 |
| Rabbit polyclonal anti-D2R | Millipore | ABN462 |
| Mouse monoclonal anti-NMDAR2B NT | Millipore | MAB5782 |
| Duolink® anti-Rabbit Plus | Merck | DUO92002 |
| Duolink® anti-Mouse Minus | Merck | DUO92004 |
| Duolink® anti-Mouse Plus | Merck | DUO92001 |
| Duolink® Anti-Goat Minus probe | Merck | DUO92006 |
| **Bacterial and virus strains** | | |
| AAV9-TRE-RFA-TetON3G-gluc1n1 | BiolMol MIRCen / CEA /Institut de biologie François Jacob | N/A |
| AAV9-TRE-RFA-TetON3G-gluc1n1d | BiolMol MIRCen / CEA /Institut de biologie François Jacob | N/A |
| AAV9-TRE-RFP-D2GluN2 | BiolMol MIRCen / CEA /Institut de biologie François Jacob | N/A |
| AAV9-TRE-RFP-D2GluN2.scramble2 | BiolMol MIRCen / CEA /Institut de biologie François Jacob | N/A |
| AAV8-hSyn1-dlox-hM4D(Gi)_mCherry(rev)-dlox-WPRE-hGHp(A) | ETH Zurich Viral Vector Facility | v84-8 |
| AAV8-hSyn1-dlox-mCherry(rev)-dlox-WPRE-hGHp(A) | ETH Zurich Viral Vector Facility | V116-8 |
| AAV-9/2-hSyn1-GRAB(DA2m)-WPRE-hGHp(A) | ETH Zurich Viral Vector Facility | v677-9 |
| AAV-DJ-hSyn1-dlox-jGCaMP8m(rev)-dlox-WPRE-SV40p(A) | ETH Zurich Viral Vector Facility | v628 |
| **Chemicals, peptides, and recombinant proteins** | | |
| Clozapine-N-Oxide BML-NS105-0025 | ENZO | CAS 34233-69-7 |
| 9-tert-butyl doxycycline HCL 117B-0801-1g | Tebu-bio / Echelon Biosciences | CAS 233585-94-9 |
| DOXYCYCLINE HYCLATE D9891-25G | Merck | CAS 24390-14-5 |
| Duolink® In Situ Wash Buffer, Brightfield | Merck | DUO82047 |
| **Critical commercial assays** | | |
| Duolink® Brightfield Detection kit | | DUO92012 |

| | | |
|---|---|---|
| **Experimental models: Organisms/strains** | | |
| C57BL/6J | Janvier Laboratories | CSAL (Orléans) - 1993 (F172) |
| Tg(Drd1-cre)FK150Gsat/Mmucd | GENSAT | MMRC: 029178-UCD |
| Tg(Drd2-cre)ER44Gsat/Mmucd | GENSAT | MMRC: 017263-UCD |
| $DAT^{IREScre}$ B6.SJL-Slc6a3tm1.1(cre)Bkmn/J | The Jackson Laboratory | RRID:IMSR_JAX:006660 |
| **Software and algorithms** | | |
| GraphPad-Prism 10 | GraphPad Software | RRID: SCR_002798 |
| MATLAB R2023b | MathWorks | RRID:SCR_001622 |
| PSY | Kevin Bird, Dusan Hadzi-Pavlovic, and Andrew Isaac © School of Psychology, University of New South Wales | N/A |
| Multichannel Fiber Photometry Software | RWD | v2.0.0.33169 |
| QuPath v.0.5.1 | Bankhead, Peter et al. “QuPath: Open source software for digital pathology image analysis.” Scientific reports vol. 7,1 16878. 4 Dec. 2017, doi:10.1038/s41598-017-17204-5 | RRID:SCR_018257 |
| FIJI (ImageJ) | Schindelin, Johannes et al. “Fiji: an open-source platform for biological-image analysis.” Nature methods vol. 9,7 676-82. 28 Jun. 2012, doi:10.1038/nmeth.2019 | RRID:SCR_002285 |
| NDP.view2 | Hamamatsu | RRID:SCR_025177 |
| **Other** | | |
| Dustless precision pellets, Grain based, 20mg | Phymep / Bio-Serv | F0163 |
| Dustless Precision Pellets 20mg, Rodent Purified diet | Phymep / Bio-Serv | F0071 |
| Sweetened Condensed Milk | Nestlé | N/A |

Supplementary Table 1

| **Data** | **Statistical test** | **n** | **Outcome measure** | **P value** |
|---|---|---|---|---|
| Figure 1 DAT VTA Gi cpt | | | | |
| Figure 1. E | Two-Factor RM ANOVA | Control mCherry = 10<br>DREADD Gi = 10 | Lever Presses / minute | DREADD Effect F(1,18) = 0.2827, p=0.6014<br>Training Effect F(7,126) = 1.627, p=0.1337<br>Interaction F (7, 126) = 0.6749, p=0.6930 |
| Figure 1. F | Planned Orthogonal Contrast<br><br>Two-Factor RM ANOVA | Control mCherry = 9<br>DREADD Gi = 10 | Lever presses | DREADD Effect F(1,17) = 0.554, p=0.4669<br>Devaluation Effect F(1,17) = 11.094, p=0.0040<br>Interaction F(1,17) = 0.246, p=0.6263 |
| Figure 1. G | One-sample t-test against 0.5<br><br>One-sample t-test against 0.5<br><br>Unpaired t-test (two-tailed) | Control mCherry = 9<br><br>DREADD Gi = 10 | Devaluation Ratio | t=3.354, df=8, p= 0.0100<br><br>t=3.929, df=9, p= 0.0035<br><br>t=1.159, df=17, p=0.2625 |
| Figure 1. I | Two-Factor RM ANOVA | Control DREADD Gi Cre- = 11<br>DREADD Gi Cre+ = 11 | Lever Presses / minute | DREADD Effect F(1,20) = 0.2827, p=0.6014<br>Training Effect F(3,60) = 261.0, p<0.0001<br>Interaction F (3, 60) = 0.2844, p=0.8364 |
| Figure 1. J | Planned Orthogonal Contrast<br><br>Two-Factor RM ANOVA | Control DREADD Gi Cre- = 11<br>DREADD Gi Cre+ = 11 | Lever presses | DREADD Effect F(1,20) = 11.541, p=0.0029<br>Devaluation Effect F(1, 20) = 19.436, p=0.0003<br>Interaction F(1, 20) = 1.458, p=0.2413 |
| Figure 1. K | One-sample t-test against 0.5<br><br>One-sample t-test against 0.5<br><br>Unpaired t-test (two-tailed) | Control DREADD Gi Cre- = 11<br><br>DREADD Gi Cre+ = 11 | Devaluation Ratio | t=3.533, df=10, p= 0.0054<br><br>t=2.212, df=10, p= 0.0514<br><br>t=0.03175, df=20, p= 0.9750 |
| Figure 1. M | Two-Factor RM ANOVA | Control DREADD Gi Cre- = 11<br>DREADD Gi Cre+ = 11 | Lever Presses / minute | DREADD Effect F(1,20) = 1.246, p=0.2775<br>Training Effect F(3,60) = 5.130, p=0.0032<br>Interaction F (3, 60) = 0.1289, p=0.9426 |
| Figure 1. N | Planned Orthogonal Contrast<br><br>Two-Factor RM ANOVA | Control DREADD Gi Cre- = 10<br>DREADD Gi Cre+ = 11 | Lever presses | DREADD Effect F(1,19) = 0.054, p=0.8187<br>Devaluation Effect F(1, 19) = 2.767, p=0.1126<br>Interaction F(1, 19) = 2.513, p=0.1294 |
| Figure 1. O | One-sample t-test against 0.5 | DREADD Gi Cre- = 10 | Devaluation Ratio | t=2.771, df=9, p= 0.0217 |

| | | | | |
|---|---|---|---|---|
| | One-sample t-test against 0.5<br><br>Unpaired t-test (two-tailed) | DREADD Gi Cre+ = 11 | | t=1.020, df=10, p= 0.3316<br><br><br>t=2.562, df=19, p= 0.0191 |
| Figure 2 Fiber DA mPFC | | | | |
| Figure 2. E | Two-Factor RM ANOVA<br><br>One sample t-test against 0<br><br>One sample t-test against 0 | Early = 8<br>Late = 8 | AUC | Time Effect F (1, 7) = 0.1817, p=0.6827<br>Session Effect F (1, 7) = 0.1764, p=0.6871<br>Time x Session F (1, 7) = 0.5338, p=0.4887<br><br>[0.1 s ; 1 s] Early<br>t=0.2343, df=7, p=0.8214<br>[0.1 s ; 1 s] Late<br>t=1.166, df=7, p=0.2817<br><br>[1.1 s ; 2 s] Early<br>t=0.2357, df=7, p=0.8204<br>[1.1 s ; 2 s] Late<br>t=0.8621, df=7, p=0.4172 |
| Figure 2. G (left) | Two-Factor RM ANOVA | Early = 8<br>Late = 8 | AUC | Time Effect F (2, 14) = 0.6544, p=0.5350<br>Training Effect F (1, 7) = 7.976, p=0.0256<br>Time x Training F (2, 14) = 4.127, p=0.0390<br><br>*Tukey's multiple comparisons test*<br>[-0.9;0]<br>Early vs. Late p=0.5112<br>[0.1;1]<br>Early vs. Late p=0.0004<br>[1.1;2]<br>Early vs. Late p=0.0047<br><br>Early<br>[-0.9;0] vs. [0.1;1] p=0.0291<br>[-0.9;0] vs. [1.1;2] p=0.3360<br>[0.1;1] vs. [1.1;2] p=0.3482<br>Late<br>[-0.9;0] vs. [0.1;1] p=0.5427<br>[-0.9;0] vs. [1.1;2] p=0.4636<br>[0.1;1] vs. [1.1;2] p=0.9895 |
| Figure 2. G (left) | One sample t-test against 0 | Early = 8<br>Late = 8 | AUC | [-0.9 s ; 0 s] Early<br>t=2.754, df=7, p=0.0283<br>[-0.9 s ; 0 s] Late<br>t=1.330, df=7, p=0.2253 |
| Figure 2. G (left) | One sample t-test against 0 | Early = 8<br>Late = 8 | AUC | [0.1 s ; 1 s] Early<br>t=4.034, df=7, p=0.0050<br>[0.1 s ; 1 s] Late<br>t=0.2853, df=7, p=0.7837 |
| Figure 2. G (left) | One sample t-test against 0 | Early = 8<br>Late = 8 | AUC | [1.1 s ; 2 s] Early<br>t=1.975, df=7, p=0.0888<br>[1.1 s ; 2 s] Late<br>t=0.3481, df=7, p=0.7380 |
| Figure 2. G (right) | One sample t-test against 0<br><br>One sample t-test against 0<br><br>Paired t-test (two-tailed) | Early = 8<br>Late = 8 | Peak Z-score | [-0.9 s ; 2 s] Early<br>t=7.243, df=7, p=0.0002<br><br>[-0.9 s ; 2 s] Late<br>t=2.994, df=7, p=0.0201<br><br>Early vs. Late<br>t=2.374, df=7, p=0.0493 |

| | | | | |
|---|---|---|---|---|
| Figure 2. I (top) | Two-Factor RM ANOVA | RL = 9<br>UL = 9 | AUC | Time Effect F (3, 24) = 19.86, p<0.0001<br>Lever Effect F (1, 8) = 0.5217, p=0.4907<br>Time x Lever F (3, 24) = 4.966, p=0.0080<br><br>*Tukey's multiple comparisons test*<br>[-0.9;0]<br>RL vs. UL p=0.4145<br>[0.1;1]<br>RL vs. UL p=0.1203<br>[1.1;2]<br>RL vs. UL p=0.3598<br>[2.1;3]<br>RL vs. UL p=0.0025<br><br>RL<br>[-0.9;0] vs. [0.1;1] p=0.0013<br>[-0.9;0] vs. [1.1;2] p=0.0079<br>[-0.9;0] vs. [2.1;3] p=0.2976<br>[0.1;1] vs. [1.1;2] p=0.8742<br>[0.1;1] vs. [2.1;3] p<0.0001<br>[1.1;2] vs. [2.1;3] p<0.0001<br>UL<br>[-0.9;0] vs. [0.1;1] p=0.2634<br>[-0.9;0] vs. [1.1;2] p=0.2968<br>[-0.9;0] vs. [2.1;3] p=0.8753<br>[0.1;1] vs. [1.1;2] p=0.9998<br>[0.1;1] vs. [2.1;3] p=0.6765<br>[1.1;2] vs. [2.1;3] p=0.7216 |
| Figure 2. I (top) | One sample t-test against 0 | RL = 9<br>UL = 9 | AUC | [-0.9 s ; 0 s] RL<br>t=0.5322, df=8, p=0.6090<br>[-0.9 s ; 0 s] UL<br>t=1.291, df=8, p=0.2329<br>[0.1 s ; 1 s] RL<br>t=3.662, df=8, p=0.0064<br>[0.1 s ; 1 s] UL<br>t=3.176, df=8, p=0.0131<br>[1.1 s ; 2 s] RL<br>t=3.814, df=8, p=0.0051<br>[1.1 s ; 2 s] UL<br>t=2.926, df=8, p=0.0191<br>[2.1 s ; 3 s] RL<br>t=2.017, df=8, p=0.0784<br>[2.1 s ; 3 s] UL<br>t=1.842, df=8, p=0.1028 |
| Figure 2. I (bottom left) | One sample t-test against 0<br><br>One sample t-test against 0<br><br>Paired t-test (two-tailed) | RL = 9<br>UL = 9 | Min Z-score | [-0.9 s ; 0 s] RL<br>t=2.129, df=8, p=0.0659<br><br>[-0.9 s ; 0 s] UL<br>t=1.294, df=8, p=0.2317<br><br>RL vs. UL<br>t=1.431, df=8, p=0.1902 |
| Figure 2. I (bottom center) | One sample t-test against 0<br><br>One sample t-test against 0<br><br>Paired t-test | RL = 9<br>UL = 9 | Peak Z-score | [0.1 s ; 3 s] RL<br>t=5.684, df=8, p=0.0005<br><br>[0.1 s ; 3 s] UL<br>t=5.981, df=8, p=0.0003<br><br>RL vs. UL |

| | | | | |
|---|---|---|---|---|
| | (two-tailed) | | | t=1.249, df=8, p=0.2469 |
| Figure 2. I (bottom right) | One sample t-test against 0<br><br>One sample t-test against 0<br><br>Paired t-test (two-tailed) | RL = 9<br>UL = 9 | Min Z-score | [0.1 s ; 3 s] RL<br>t=3.136, df=8, p=0.0139<br><br>[0.1 s ; 3 s] UL<br>t=1.830, df=8, p=0.1046<br><br>RL vs. UL<br>t=1.502, df=8, p=0.1715 |
| Figure 2. K (top) | Two-Factor RM ANOVA | RL = 9<br>UL = 9 | AUC | Time Effect F (3, 24) = 15.90, p<0.0001<br>Training Effect F (1, 8) = 5.519, p=0.0467<br>Time x Training F (3, 24) = 3.938, p=0.0204<br><br>*Tukey's multiple comparisons test*<br><br>[-0.9;0]<br>Early vs. Late p=0.0297<br>[0.1;1]<br>Early vs. Late p=0.7314<br>[1.1;2]<br>Early vs. Late p=0.0018<br>[2.1;3]<br>Early vs. Late p=0.0004<br><br>Early<br>[-0.9;0] vs. [0.1;1] p=0.0607<br>[-0.9;0] vs. [1.1;2] p=0.0010<br>[-0.9;0] vs. [2.1;3] p=0.7949<br>[0.1;1] vs. [1.1;2] p=0.3289<br>[0.1;1] vs. [2.1;3] p=0.0076<br>[1.1;2] vs. [2.1;3] p=0.0001<br>Late<br>[-0.9;0] vs. [0.1;1] p=0.0001<br>[-0.9;0] vs. [1.1;2] p=0.0188<br>[-0.9;0] vs. [2.1;3] p=0.0514<br>[0.1;1] vs. [1.1;2] p=0.1734<br>[0.1;1] vs. [2.1;3] p<0.0001<br>[1.1;2] vs. [2.1;3] p<0.0001 |
| Figure 2. K (top) | One sample t-test against 0 | RL = 9<br>UL = 9 | AUC | [-0.9 s ; 0 s] Early<br>t=2.783, df=8, p=0.0238<br>[-0.9 s ; 0 s] Late<br>t=0.9540, df=8, p=0.3680<br>[0.1 s ; 1 s] Early<br>t=3.799, df=8 , p=0.0052<br>[0.1 s ; 1 s] Late<br>t=6.833, df=8, p=0.0001<br>[1.1 s ; 2 s] Early<br>t=4.008, df=8, p=0.0039<br>[1.1 s ; 2 s] Late<br>t=2.540, df=8, p=0.0347<br>[2.1 s ; 3 s] Early<br>t=0.5045, df=8, p=0.6275<br>[2.1 s ; 3 s] Late<br>t=2.409, df=8, p=0.0426 |
| Figure 2. K (bottom left) | One sample t-test against 0<br><br>One sample t-test against 0 | RL = 9<br>UL = 9 | Min Z-score | [-0.9 s ; 0 s] RL<br>t=0.2008, df=8, p=0.8459<br><br>[-0.9 s ; 0 s] UL<br>t=2.800, df=8, p=0.0232 |

| | | | | |
|---|---|---|---|---|
| | Paired t-test (two-tailed) | | | RL vs. UL<br>t=2.464, df=8, p=0.0391 |
| Figure 2. K (bottom center) | One sample t-test against 0<br><br>One sample t-test against 0<br><br>Paired t-test (two-tailed) | RL = 9<br>UL = 9 | Peak Z-score | [0.1 s ; 3 s] RL<br>t=6.003, df=8, p=0.0003<br><br>[0.1 s ; 3 s] UL<br>t=9.749, df=8, p<0.0001<br><br>RL vs. UL<br>t=1.237, df=8, p=0.2510 |
| Figure 2. K (bottom right) | One sample t-test against 0<br><br>One sample t-test against 0<br><br>Paired t-test (two-tailed) | RL = 9<br>UL = 9 | Min Z-score | [0.1 s ; 3 s] RL<br>t=2.201, df=8, p=0.0589<br><br>[0.1 s ; 3 s] UL<br>t=3.864, df=8, p=0.0048<br><br>RL vs. UL<br>t=1.492, df=8, p=0.1740 |
| Figure 2. M (top) | Two-Factor RM ANOVA | RL = 9<br>UL = 9 | AUC | Time Effect F (3, 24) = 39.96, p<0.0001<br>Lever Effect F (1, 8) = 0.5959, p=0.4623<br>Time x Lever F (3, 24) = 16.20, p<0.0001<br><br>*Tukey's multiple comparisons test*<br>[-0.9;0]<br>RL vs. UL p=0.3839<br>[0.1;1]<br>RL vs. UL p=0.0015<br>[1.1;2]<br>RL vs. UL p=0.3089<br>[2.1;3]<br>RL vs. UL p<0.0001<br><br>RL<br>[-0.9;0] vs. [0.1;1] p<0.0001<br>[-0.9;0] vs. [1.1;2] p=0.0072<br>[-0.9;0] vs. [2.1;3] p=0.0002<br>[0.1;1] vs. [1.1;2] p<0.0001<br>[0.1;1] vs. [2.1;3] p<0.0001<br>[1.1;2] vs. [2.1;3] p<0.0001<br>UL<br>[-0.9;0] vs. [0.1;1] p=0.0009<br>[-0.9;0] vs. [1.1;2] p=0.3561<br>[-0.9;0] vs. [2.1;3] p>0.9999<br>[0.1;1] vs. [1.1;2] p=0.0463<br>[0.1;1] vs. [2.1;3] p=0.0008<br>[1.1;2] vs. [2.1;3] p=0.3517 |
| Figure 2. M (top) | One sample t-test against 0 | RL = 9<br>UL = 9 | AUC | [-0.9 s ; 0 s] RL<br>t=1.779, df=8, p=0.1130<br>[-0.9 s ; 0 s] UL<br>t=0.08946, df=8, p=0.9309<br>[0.1 s ; 1 s] RL<br>t=9.813, df=8, p<0.0001<br>[0.1 s ; 1 s] UL<br>t=3.884, df=8, p=0.0046<br>[1.1 s ; 2 s] RL<br>t=1.979, df=8, p=0.0832<br>[1.1 s ; 2 s] UL<br>t=1.840, df=8, p=0.1031 |

| | | | | |
|---|---|---|---|---|
| | | | | [2.1 s ; 3 s] RL<br>t=4.772, df=8, p=0.0014<br>[2.1 s ; 3 s] UL<br>t=0.09581, df=8, p=0.9260 |
| Figure 2. M (bottom left) | One sample t-test against 0<br><br>One sample t-test against 0<br><br>Paired t-test (two-tailed) | RL = 9<br>UL = 9 | Min Z-score | [-0.9 s ; 0 s] RL<br>t=4.309, df=8, p=0.0026<br><br>[-0.9 s ; 0 s] UL<br>t=2.812, df=8, p=0.0228<br><br>RL vs. UL<br>t=2.529, df=8, p=0.0353 |
| Figure 2. M (bottom center) | One sample t-test against 0<br><br>One sample t-test against 0<br><br>Paired t-test (two-tailed) | RL = 9<br>UL = 9 | Peak Z-score | [0.1 s ; 3 s] RL<br>t=10.89, df=8, p<0.0001<br><br>[0.1 s ; 3 s] UL<br>t=5.500, df=8, p=0.0006<br><br>RL vs. UL<br>t=4.447, df=8, p=0.0021 |
| Figure 2. M (bottom right) | One sample t-test against 0<br><br>One sample t-test against 0<br><br>Paired t-test (two-tailed) | RL = 9<br>UL = 9 | Min Z-score | [0.1 s ; 3 s] RL<br>t=6.581, df=8, p=0.0002<br><br>[0.1 s ; 3 s] UL<br>t=4.847, df=8, p=0.0013<br><br>RL vs. UL<br>t=4.345, df=8, p=0.0025 |
| Figure 2. O (top) | Two-Factor RM ANOVA | RL = 9<br>UL = 9 | AUC | Time Effect F (3, 24) = 10.41, p=0.0001<br>Lever Effect F (1, 8) = 9.697, p=0.0144<br>Time x Lever F (3, 24) = 6.118, p=0.0030<br><br>*Tukey's multiple comparisons test*<br>[-0.9;0]<br>RL vs. UL p=0.5433<br>[0.1;1]<br>RL vs. UL p=0.0126<br>[1.1;2]<br>RL vs. UL p<0.0001<br>[2.1;3]<br>RL vs. UL p=0.2644<br><br>RL<br>[-0.9;0] vs. [0.1;1] p<0.0001<br>[-0.9;0] vs. [1.1;2] p<0.0001<br>[-0.9;0] vs. [2.1;3] p=0.8904<br>[0.1;1] vs. [1.1;2] p=0.7123<br>[0.1;1] vs. [2.1;3] p=0.0001<br>[1.1;2] vs. [2.1;3] p<0.0001<br>UL<br>[-0.9;0] vs. [0.1;1] p=0.0038<br>[-0.9;0] vs. [1.1;2] p=0.4363<br>[-0.9;0] vs. [2.1;3] p=0.9976<br>[0.1;1] vs. [1.1;2] p=0.1161<br>[0.1;1] vs. [2.1;3] p=0.0060<br>[1.1;2] vs. [2.1;3] p=0.5469 |
| Figure 2. O (top) | One sample t-test against 0 | RL = 9<br>UL = 9 | AUC | [-0.9 s ; 0 s] RL<br>t=0.01613, df=8, p=0.9875<br>[-0.9 s ; 0 s] UL |

| | | | | |
|---|---|---|---|---|
| | | | | t=0.9693, df=8, p=0.3608<br>[0.1 s ; 1 s] RL<br>t=4.947, df=8, p=0.0011<br>[0.1 s ; 1 s] UL<br>t=3.542, df=8, p=0.0076<br>[1.1 s ; 2 s] RL<br>t=4.466, df=8, p=0.0021<br>[1.1 s ; 2 s] UL<br>t=0.9789, df=8, p=0.3563<br>[2.1 s ; 3 s] RL<br>t=0.6949, df=8, p=0.5068<br>[2.1 s ; 3 s] UL<br>t=0.6161, df=8, p=0.5550 |
| Figure 2. O (bottom left) | One sample t-test against 0<br><br>One sample t-test against 0<br><br>Paired t-test (two-tailed) | RL = 9<br>UL = 9 | Min Z-score | [-0.9 s ; 0 s] RL<br>t=1.823, df=8, p=0.1058<br><br>[-0.9 s ; 0 s] UL<br>t=3.122, df=8, p=0.0142<br><br>RL vs. UL<br>t=0.5833, df=8, p=0.5758 |
| Figure 2. O (bottom center) | One sample t-test against 0<br><br>One sample t-test against 0<br><br>Paired t-test (two-tailed) | RL = 9<br>UL = 9 | Peak Z-score | [0.1 s ; 3 s] RL<br>t=6.498, df=8, p=0.0002<br><br>[0.1 s ; 3 s] UL<br>t=5.844, df=8, p=0.0004<br><br>RL vs. UL<br>t=4.335, df=8, p=0.0025 |
| Figure 2. O (bottom right) | One sample t-test against 0<br><br>One sample t-test against 0<br><br>Paired t-test (two-tailed) | RL = 9<br>UL = 9 | Min Z-score | [0.1 s ; 3 s] RL<br>t=3.202, df=8, p=0.0126<br><br>[0.1 s ; 3 s] UL<br>t=4.947, df=8, p=0.0011<br><br>RL vs. UL<br>t=0.3742, df=8, p=0.7180 |
| Figure 2. Q (top) | Two-Factor RM ANOVA | RL = 9<br>UL = 9 | AUC | Time Effect F (3, 24) = 27.27, p<0.0001<br>Lever Effect F (1, 8) = 20.36, p=0.0020<br>Time x Lever F (3, 24) = 8.455, p=0.0005<br><br>*Tukey's multiple comparisons test*<br>[-0.9;0]<br>RL vs. UL p=0.0016<br>[0.1;1]<br>RL vs. UL p=0.8558<br>[1.1;2]<br>RL vs. UL p=0.0257<br>[2.1;3]<br>RL vs. UL p<0.0001<br><br>RL<br>[-0.9;0] vs. [0.1;1] p<0.0001<br>[-0.9;0] vs. [1.1;2] p=0.4227<br>[-0.9;0] vs. [2.1;3] p=0.0016<br>[0.1;1] vs. [1.1;2] p=0.0007<br>[0.1;1] vs. [2.1;3] p<0.0001<br>[1.1;2] vs. [2.1;3] p<0.0001<br>UL |

| | | | | |
|---|---|---|---|---|
| | | | | [-0.9;0] vs. [0.1;1] p=0.1083<br>[-0.9;0] vs. [1.1;2] p=0.9816<br>[-0.9;0] vs. [2.1;3] p=0.7728<br>[0.1;1] vs. [1.1;2] p=0.2141<br>[0.1;1] vs. [2.1;3] p=0.0136<br>[1.1;2] vs. [2.1;3] p=0.5503 |
| Figure 2. Q (top) | One sample t-test against 0 | RL = 9<br>UL = 9 | AUC | [-0.9 s ; 0 s] RL<br>t=1.407, df=8, p=0.1970<br>[-0.9 s ; 0 s] UL<br>t=2.806, df=8, p=0.0230<br>[0.1 s ; 1 s] RL<br>t=4.033, df=8, p=0.0038<br>[0.1 s ; 1 s] UL<br>t=6.221, df=8, p=0.0003<br>[1.1 s ; 2 s] RL<br>t=0.5132, df=8, p=0.6216<br>[1.1 s ; 2 s] UL<br>t=3.154, df=8, p=0.0135<br>[2.1 s ; 3 s] RL<br>t=4.036, df=8, p=0.0038<br>[2.1 s ; 3 s] UL<br>t=1.459, df=8, p=0.1826 |
| Figure 2. Q (bottom left) | One sample t-test against 0<br><br>One sample t-test against 0<br><br>Paired t-test (two-tailed) | RL = 9<br>UL = 9 | Min Z-score | [-0.9 s ; 0 s] RL<br>t=2.913, df=8, p=0.0195<br><br>[-0.9 s ; 0 s] UL<br>t=0.8807, df=8, p=0.4042<br><br>RL vs. UL<br>t=2.287, df=8, p=0.0515 |
| Figure 2. Q (bottom center) | One sample t-test against 0<br><br>One sample t-test against 0<br><br>Paired t-test (two-tailed) | RL = 9<br>UL = 9 | Peak Z-score | [0.1 s ; 3 s] RL<br>t=5.697, df=8, p=0.0005<br><br>[0.1 s ; 3 s] UL<br>t=8.778, df=8, p<0.0001<br><br>RL vs. UL<br>t=0.8849, df=8, p=0.4020 |
| Figure 2. Q (bottom right) | One sample t-test against 0<br><br>One sample t-test against 0<br><br>Paired t-test (two-tailed) | RL = 9<br>UL = 9 | Min Z-score | [0.1 s ; 3 s] RL<br>t=5.224, df=8, p=0.0008<br><br>[0.1 s ; 3 s] UL<br>t=1.503, df=8, p=0.1712<br><br>RL vs. UL<br>t=2.764, df=8, p=0.0245 |
| Figure 3 Fiber Peptide mPFC CTRL | | | | |
| Figure 3.E (top) | Two-Factor RM ANOVA | GluN1-Δ Control peptide Early / Late = 5 | AUC | Time Effect F(1, 4) = 7.732, p=0.0498<br>Training Effect F(1, 4) = 7.027, p=0.0569<br>Time x Training F(1, 4) = 6.514, p=0.0631 |
| Figure 3.E (top) | One-sample t-test against 0 | GluN1-Δ Control peptide Early / Late = 5 | AUC | [0.1 s ; 1 s] Early<br>t=3.760, df=4, p=0.0198<br>[0.1 s ; 1 s] Late<br>t=1.191, df=4, p=0.2996<br>[1.1 s ; 2 s] Early<br>t=2.823, df=4, p=0.0477<br>[1.1 s ; 2 s] Late<br>t=3.144, df=4, p=0.0347 |

| | | | | |
|---|---|---|---|---|
| Figure 3.E (bottom) | One-sample t-test against 0<br><br>One-sample t-test against 0<br><br>Paired t-test (two-tailed) | GluN1-Δ Control peptide Early / Late = 5 | Peak Z-score | [0.1 s ; 1 s] Early<br>t=5.513, df=4, p= 0.0053<br><br>[0.1 s ; 1 s] Late<br>t=2.240, df=4, p= 0.0887<br><br>[0.1 s ; 1 s]<br>t=0.4407, df=4, p= 0.6822 |
| Figure 3.G (top) | Two-Factor RM ANOVA | D2R-Scr Control peptide Early / Late = 6 | AUC | Time Effect F 1, 5) = 2.474, p=0.1766<br>Training Effect F(1, 5) = 78.47, p=0.0003<br>Time x Training F(1, 5) = 73.39, p=0.0004<br><br>*Šídák's multiple comparisons test*<br><br>[0.1;1]<br>Early vs. Late p=0.0054<br>[1.1;2]<br>Early vs. Late p<0.0001<br>Early<br>[0.1;1] vs. [1.1;2] p=0.0716<br>Late<br>[0.1;1] vs. [1.1;2] p=0.0005 |
| Figure 3.G (top) | One-sample t-test against 0 | D2R-Scr Control peptide Early / Late = 6 | AUC | [0.1 s ; 1 s] Early<br>t=4.973, df=5, p=0.0042<br>[0.1 s ; 1 s] Late<br>t=1.594, df=5, p=0.1719<br>[1.1 s ; 2 s] Early<br>t=5.701, df=5, p=0.0023<br>[1.1 s ; 2 s] Late<br>t=2.111, df=5, p=0.0885 |
| Figure 3.G (bottom) | One-sample t-test against 0<br><br>One-sample t-test against 0<br><br>Paired t-test (two-tailed) | D2R-Scr Control peptide Early / Late = 6 | Peak Z-score | [0.1 s ; 1 s] Early<br>t=6.731, df=5, p=0.0011<br><br>[0.1 s ; 1 s] Late<br>t=3.065, df=5, p=0.0279<br><br>[0.1 s ; 1 s]<br>t=2.228, df=5, p=0.0764 |
| Figure 3.I (top) | Two-Factor RM ANOVA | GluN1-Δ Control peptide Early / Late = 5 | AUC | Time Effect F(2, 8) = 7.568, p=0.0143<br>Training Effect F(1, 4) = 17.05, p=0.0145<br>Time x Training F(2, 8) = 8.499, p=0.0105<br><br>*Tukey's multiple comparisons test*<br><br>[-0.9;0]<br>Early vs. Late p=0.9470<br>[0.1;1]<br>Early vs. Late p=0.0005<br>[1.1;2]<br>Early vs. Late p=0.0039<br><br>Early<br>[-0.9;0] vs. [0.1;1] p=0.0021<br>[-0.9;0] vs. [1.1;2] p=0.4213<br>[0.1;1] vs. [1.1;2] p=0.0114<br>Late<br>[-0.9;0] vs. [0.1;1] p=0.9026<br>[-0.9;0] vs. [1.1;2] p=0.0578<br>[0.1;1] vs. [1.1;2] p=0.1093 |

| | | | | |
|---|---|---|---|---|
| Figure 3.I (top) | One-sample t-test against 0 | GluN1-Δ Control peptide Early / Late = 5 | AUC | [-0.9 s ; 0 s] Early<br>t=1.999, df=4, p=0.1162<br>[-0.9 s ; 0 s] Late<br>t=2.487, df=4, p=0.0677<br>[0.1 s ; 1 s] Early<br>t=4.956, df=4, p=0.0077<br>[0.1 s ; 1 s] Late<br>t=0.9711, df=4, p=0.3865<br>[1.1 s ; 2 s] Early<br>t=2.097, df=4, p=0.1040<br>[1.1 s ; 2 s] Late<br>t=1.234, df=4, p=0.2846 |
| Figure 3.I (bottom left) | One-sample t-test against 0<br><br>One-sample t-test against 0<br><br>Paired t-test (two-tailed) | GluN1-Δ Control peptide Early / Late = 5 | Peak Z-score | [-0.9 s ; 0 s] Early<br>t=3.944, df=4, p= 0.0169<br><br>[-0.9 s ; 0 s] Late<br>t=3.660, df=4, p= 0.0216<br><br>[-0.9 s ; 0 s]<br>t=0.5063, df=4, p= 0.6393 |
| Figure 3.I (bottom right) | One-sample t-test against 0<br><br>One-sample t-test against 0<br><br>Paired t-test (two-tailed) | GluN1-Δ Control peptide Early / Late = 5 | Peak Z-score | [0.1 s ; 1 s] Early<br>t=4.759, df=4, p= 0.0089<br><br>[0.1 s ; 1 s] Late<br>t=2.104, df=4, p= 0.1032<br><br>[0.1 s ; 1 s]<br>t=3.030, df=4, p= 0.0388 |
| Figure 3.K (top) | Two-Factor RM ANOVA | D2R-Scr Control peptide Early / Late = 6 | AUC | Time Effect F(2, 10) = 6.915, p=0.0130<br>Training Effect F(1, 5) = 5.086, p=0.0738<br>Time x Training F(2, 10) =7.506, p=0.0102<br><br>*Tukey's multiple comparisons test*<br><br>[-0.9;0]<br>Early vs. Late p=0.4302<br>[0.1;1]<br>Early vs. Late p=0.0013<br>[1.1;2]<br>Early vs. Late p=0.0105<br><br>Early<br>[-0.9;0] vs. [0.1;1] p=0.0231<br>[-0.9;0] vs. [1.1;2] p=0.9535<br>[0.1;1] vs. [1.1;2] p=0.0143<br>Late<br>[-0.9;0] vs. [0.1;1] p=0.1519<br>[-0.9;0] vs. [1.1;2] p=0.0043<br>[0.1;1] vs. [1.1;2] p=0.1167 |
| Figure 3.K (top) | One-sample t-test against 0 | D2R-Scr Control peptide Early / Late = 6 | AUC | [-0.9 s ; 0 s] Early<br>t=2.088, df=5, p=0.0912<br>[-0.9 s ; 0 s] Late<br>t=3.872, df=5, p=0.0117<br>[0.1 s ; 1 s] Early<br>t=4.172, df=5, p=0.0087<br>[0.1 s ; 1 s] Late<br>t=0.4764, df=5, p=0.6539<br>[1.1 s ; 2 s] Early<br>t=0.8322, df=5, p=0.4432<br>[1.1 s ; 2 s] Late |

| | | | | |
|---|---|---|---|---|
| | | | | t=4.339, df=5, p= 0.0074 |
| Figure 3.K (bottom left) | One-sample t-test against 0<br><br>One-sample t-test against 0<br><br>Paired t-test (two-tailed) | D2R-Scr Control peptide Early / Late = 6 | Peak Z-score | [-0.9 s ; 0 s] Early<br>t=3.792, df=5, p=0.0127<br><br>[-0.9 s ; 0 s] Late<br>t=5.226, df=5, p=0.0034<br><br>[-0.9 s ; 0 s]<br>t=1,184, df=5, p=0,2895 |
| Figure 3.K (bottom right) | One-sample t-test against 0<br><br>One-sample t-test against 0<br><br>Paired t-test (two-tailed) | D2R-Scr Control peptide Early / Late = 6 | Peak Z-score | [0.1 s ; 1 s] Early<br>t=5.255, df=5, p=0.0033<br><br>[0.1 s ; 1 s] Late<br>t=3.532, df=5, p=0.0167<br><br>[0.1 s ; 1 s]<br>t=3.408, df=5, p=0.0191 |
| Figure 3.M (top) | Two-Factor RM ANOVA | GluN1-Δ Control peptide RL = 5 | AUC | Time Effect F(2, 8) = 1.799, p=0.2264<br>Lever Effect F(1, 4) = 4.230, p=0.1089<br>Time x Lever F(2, 8) = 7.442, p=0.0149<br><br>*Tukey's multiple comparisons test*<br><br>[-0.9;0]<br>RL vs. UL p=0.2117<br>[0.1;1]<br>RL vs. UL p=0.0002<br>[1.1;2]<br>RL vs. UL p=0.0642<br><br>RL<br>[-0.9;0] vs. [0.1;1] p=0.0174<br>[-0.9;0] vs. [1.1;2] p=0.9988<br>[0.1;1] vs. [1.1;2] p=0.0162<br>UL<br>[-0.9;0] vs. [0.1;1] p=0.3490<br>[-0.9;0] vs. [1.1;2] p=0.6925<br>[0.1;1] vs. [1.1;2] p=0.8004 |
| Figure 3.M (top) | One-sample t-test against 0 | GluN1-Δ Control peptide UL = 5 | AUC | [-0.9 s ; 0 s] RL<br>t=4.058, df=4, p=0.0154<br>[-0.9 s ; 0 s] UL<br>t=1.631, df=4, p=0.1781<br>[0.1 s ; 1 s] RL<br>t=3.496, df=4, p=0.0087<br>[0.1 s ; 1 s] UL<br>t=0.2778, df=4, p=0.0250<br>[1.1 s ; 2 s] RL<br>t=1.550, df=4, p=0.1960<br>[1.1 s ; 2 s] UL<br>t=1.851, df=4, p= 0.1378 |
| Figure 3.M (bottom left) | One-sample t-test against 0<br><br>One-sample t-test against 0<br><br>Paired t-test (two-tailed) | GluN1-Δ Control peptide RL / UL = 5 | Peak Z-score | [-0.9 s ; 0 s] RL<br>t=5.554, df=4, p= 0.0051<br><br>[-0.9 s ; 0 s] UL<br>t=2.199, df=4, p= 0.0928<br><br>[-0.9 s ; 0 s]<br>t=2.042, df=4, p= 0.1107 |

| | | | | |
|---|---|---|---|---|
| Figure 3.M (bottom right) | One-sample t-test against 0<br><br>One-sample t-test against 0<br><br>Paired t-test (two-tailed) | GluN1-Δ Control peptide RL / UL = 5 | Peak Z-score | [0.1 s ; 1 s] RL<br>t=4.542, df=4, p= 0.0105<br><br>[0.1 s ; 1 s] UL<br>t=2.114, df=4, p= 0.1020<br><br>[0.1 s ; 1 s]<br>t=4.106, df=4, p= 0.0148 |
| Figure 3.O (top) | Two-Factor RM ANOVA | D2R-Scr Control peptide RL / UL = 6 | AUC | Time Effect F(2, 10) = 39.15, p<0.0001<br>Lever Effect F(1, 5) = 15.75, p=0.0106<br>Time x Lever F(2, 10) = 11.45, p=0.0026<br><br>*Tukey's multiple comparisons test*<br><br>[-0.9;0]<br>RL vs. UL p=0.0460<br>[0.1;1]<br>RL vs. UL p<0.0001<br>[1.1;2]<br>RL vs. UL p=0.0179<br><br>RL<br>[-0.9;0] vs. [0.1;1] p=0.0057<br>[-0.9;0] vs. [1.1;2] p=0.0169<br>[0.1;1] vs. [1.1;2] p<0.0001<br>UL<br>[-0.9;0] vs. [0.1;1] p=0.1525<br>[-0.9;0] vs. [1.1;2] p=0.0070<br>[0.1;1] vs. [1.1;2] p=0.1862 |
| Figure 3.O (top) | One-sample t-test against 0 | D2R-Scr Control peptide RL / UL = 6 | AUC | [-0.9 s ; 0 s] RL<br>t=4.208, df=5, p=0.0084<br>[-0.9 s ; 0 s] UL<br>t=3.390, df=5, p=0.0195<br>[0.1 s ; 1 s] RL<br>t=4.772, df=5, p=0.0050<br>[0.1 s ; 1 s] UL<br>t=0.7470, df=5, p=0.4887<br>[1.1 s ; 2 s] RL<br>t=1.096, df=5, p=0.3229<br>[1.1 s ; 2 s] UL<br>t=1.372, df=5, p= 0.2285 |
| Figure 3.O (bottom left) | One-sample t-test against 0<br><br>One-sample t-test against 0<br><br>Paired t-test (two-tailed) | D2R-Scr Control peptide RL / UL = 6 | Peak Z-score | [-0.9 s ; 0 s] RL<br>t=4.919, df=5, p=0.0044<br><br>[-0.9 s ; 0 s] UL<br>t=4.815, df=5, p= 0.0048<br><br>[-0.9 s ; 0 s]<br>t=1.643, df=5, p= 0.1612 |
| Figure 3.O (bottom right) | One-sample t-test against 0<br><br>One-sample t-test against 0<br><br>Paired t-test (two-tailed) | D2R-Scr Control peptide RL / UL = 6 | Peak Z-score | [0.1 s ; 1 s] RL<br>t=5.351, df=5, p= 0.0031<br><br>[0.1 s ; 1 s] UL<br>t=1.862, df=5, p= 0.1216<br><br>[0.1 s ; 1 s]<br>t=3.852, df=5, p= 0.0120 |
| Figure 3.Q (top) | Two-Factor RM ANOVA | GluN1-Δ Control peptide RL / UL = 5 | AUC | Time Effect F (2, 8) = 4.362, p=0.0524<br>Lever Effect F (1, 4) = 8.690, p=0.0421<br>Time x Lever F (2, 8) = 1.260, p=0.3343 |

| | | | | |
|---|---|---|---|---|
| Figure 3.Q (top) | One-sample t-test against 0 | GluN1-Δ Control peptide RL / UL = 5 | AUC | [-0.9 s ; 0 s] RL<br>t=2.284, df=4, p=0.0845<br>[-0.9 s ; 0 s] UL<br>t=0.2191, df=4, p=0.8373<br>[0.1 s ; 1 s] RL<br>t=1.250, df=4, p=0.2793<br>[0.1 s ; 1 s] UL<br>t=3.169, df=4, p=0.0339<br>[1.1 s ; 2 s] RL<br>t=0.2717, df=4, p=0.7993<br>[1.1 s ; 2 s] UL<br>t=2.872, df=4, p= 0.0454 |
| Figure 3.Q (top insert) | One-sample t-test against 0<br><br>One-sample t-test against 0<br><br>Paired t-test (two-tailed) | GluN1-Δ Control peptide RL / UL = 10<br><br>(Pool of two AUC intervals) | AUC | RL<br>t=2.518, df=9, p=0.0329<br><br>UL<br>t=1.356, df=9, p=0.2081<br><br>RL vs. UL<br>t=3.832, df=9, p=0.0040 |
| Figure 3.Q (bottom left) | One-sample t-test against 0<br><br>One-sample t-test against 0<br><br>Paired t-test (two-tailed) | GluN1-Δ Control peptide RL / UL = 5 | Peak Z-score | [-0.9 s ; 0 s] RL<br>t=2.917, df=4, p= 0.0434<br><br>[-0.9 s ; 0 s] UL<br>t=1.928, df=4, p= 0.1261<br><br>[-0.9 s ; 0 s]<br>t=2.358, df=4, p= 0.0778 |
| Figure 3.Q (bottom right) | One-sample t-test against 0<br><br>One-sample t-test against 0<br><br>Paired t-test (two-tailed) | GluN1-Δ Control peptide RL / UL = 5 | Peak Z-score | [0.1 s ; 1 s] RL<br>t=2.620, df=4, p= 0.0588<br><br>[0.1 s ; 1 s] UL<br>t=1.051, df=4, p= 0.3526<br><br>[0.1 s ; 1 s]<br>t=2.455, df=4, p= 0.0701 |
| Figure 3.S (top) | Two-Factor RM ANOVA | D2R-Scr Control peptide RL / UL = 6 | AUC | Time Effect F(2, 10) = 3.600, p=0.0664<br>Lever Effect F(1, 5) = 0.4951, p=0.5131<br>Time x Lever F(2, 10) = 1.828, p=0.2106 |
| Figure 3.S (top) | One-sample t-test against 0 | D2R-Scr Control peptide RL / UL = 6 | AUC | [-0.9 s ; 0 s] RL<br>t=0.8506, df=5, p=0.4338<br>[-0.9 s ; 0 s] UL<br>t=1.135, df=5, p=0.3080<br>[0.1 s ; 1 s] RL<br>t=0.1968, df=5, p=0.8517<br>[0.1 s ; 1 s] UL<br>t=2.144, df=5, p=0.0849<br>[1.1 s ; 2 s] RL<br>t=3.320, df=5, p=0.0210<br>[1.1 s ; 2 s] UL<br>t=1.262, df=5, p= 0.2627 |
| Figure 3.S (top insert) | One-sample t-test against 0<br><br>One-sample t-test against 0<br><br>Paired t-test (two-tailed) | D2R-Scr Control peptide RL / UL = 12<br><br>(Pool of two AUC intervals) | AUC | RL<br>t=0.4107, df=11, p=0.6892<br><br>UL<br>t=2,.298, df=11, p=0.0422<br><br>RL vs. UL<br>t=1.979, df=11, p=0.0734 |

| | | | | |
|---|---|---|---|---|
| Figure 3.S (bottom left) | One-sample t-test against 0<br><br>One-sample t-test against 0<br><br>Paired t-test (two-tailed) | D2R-Scr Control peptide RL / UL = 6 | Peak Z-score | [-0.9 s ; 0 s] RL<br>t=1.816, df=5, p= 0.1291<br><br>[-0.9 s ; 0 s] UL<br>t=1.541, df=5, p= 0.1840<br><br>[-0.9 s ; 0 s]<br>t=0.6602, df=5, p= 0.5383 |
| Figure 3.S (bottom right) | One-sample t-test against 0<br><br>One-sample t-test against 0<br><br>Paired t-test (two-tailed) | D2R-Scr Control peptide RL / UL = 6 | Peak Z-score | [0.1 s ; 1 s] RL<br>t=1.755, df=5, p= 0.1396<br><br>[0.1 s ; 1 s] UL<br>t=0.8489, df=5, p= 0.4347<br><br>[0.1 s ; 1 s]<br>t=0.5409, df=5, p= 0.6118 |
| Figure 4 Fiber Peptide mPFC Interfering | | | | |
| Figure 4.E (top) | Two-Factor RM ANOVA | GluN1-C1 Control peptide Early / Late = 5 | AUC | Time Effect F(1, 4) = 7.427, p=0.0527<br>Training Effect F(1, 4) = 47.61, p=0.0023<br>Time x Training F(1, 4) = 5.247, p=0.0838 |
| Figure 4.E (top) | One-sample t-test against 0 | GluN1-C1 Control peptide Early / Late = 5 | AUC | [0.1 s ; 1 s] Early<br>t=4.160, df=4, p=0.0141<br>[0.1 s ; 1 s] Late<br>t=2.225, df=4, p=0.0902<br>[1.1 s ; 2 s] Early<br>t=2.440, df=4, p=0.0712<br>[1.1 s ; 2 s] Late<br>t=0.1659, df=4, p=0.8763 |
| Figure 4.E (bottom) | One-sample t-test against 0<br><br>One-sample t-test against 0<br><br>Paired t-test (two-tailed) | GluN1-C1 Control peptide Early / Late = 5 | Peak Z-score | [0.1 s ; 1 s] Early<br>t=5.238, df=4, p= 0.0063<br><br>[0.1 s ; 1 s] Late<br>t=3.666, df=4, p= 0.0215<br><br>[0.1 s ; 1 s]<br>t=0.9838, df=4, p= 0.3809 |
| Figure 4.G (top) | Two-Factor RM ANOVA | D2R- IL3 Control peptide Early = 8 | AUC | Time Effect F(1, 7) = 15.64, p=0.0055<br>Training Effect F(1, 7) = 28.77, p=0.0010<br>Time x Training F(1, 7) = 20.30, p=0.0028<br><br>*Šídák's multiple comparisons test*<br><br>[0.1;1]<br>Early vs. Late p=0.0211<br>[1.1;2]<br>Early vs. Late p=0.0030<br>Early<br>[0.1;1] vs. [1.1;2] p=0.3204<br>Late<br>[0.1;1] vs. [1.1;2] p=0.0162 |
| Figure 4.G (top) | One-sample t-test against 0 | D2R- IL3 Control peptide Late = 8 | AUC | [0.1 s ; 1 s] Early<br>t=5.978, df=7, p= 0.0006<br>[0.1 s ; 1 s] Late<br>t=0.9190, df=7, p= 0.3886<br>[1.1 s ; 2 s] Early<br>t=4.167, df=7, p= 0.0042<br>[1.1 s ; 2 s] Late<br>t=1.967, df=7, p= 0.0899 |

| | | | | |
|---|---|---|---|---|
| Figure 4.G (bottom) | One-sample t-test against 0<br><br>One-sample t-test against 0<br><br>Paired t-test (two-tailed) | D2R- IL3 Control peptide Early / Late = 8 | Peak Z-score | [0.1 s ; 1 s] Early<br>t=6.979, df=7, p= 0.0002<br><br>[0.1 s ; 1 s] Late<br>t=4.814, df=7, p= 0.0019<br><br>[0.1 s ; 1 s]<br>t=2.097, df=7, p= 0.0742 |
| Figure 4.I (top) | Two-Factor RM ANOVA | GluN1-C1 Control peptide Early / Late = 5 | AUC | Time Effect F(2, 8) = 5.312, p=0.0340<br>Training Effect F(1, 4) = 3.371, p=0.1402<br>Time x Training F(2, 8) = 6.077, p=0.0248<br><br>*Tukey's multiple comparisons test*<br><br>[-0.9;0]<br>Early vs. Late p=0.9563<br>[0.1;1]<br>Early vs. Late p=0.0082<br>[1.1;2]<br>Early vs. Late p=0.0016<br><br>Early<br>[-0.9;0] vs. [0.1;1] p=0.0255<br>[-0.9;0] vs. [1.1;2] p=0.9864<br>[0.1;1] vs. [1.1;2] p=0.0204<br>Late<br>[-0.9;0] vs. [0.1;1] p=0.9727<br>[-0.9;0] vs. [1.1;2] p=0.0030<br>[0.1;1] vs. [1.1;2] p=0.0040 |
| Figure 4.I (top) | One-sample t-test against 0 | GluN1-C1 Control peptide Early / Late = 5 | AUC | [-0.9 s ; 0 s] Early<br>t=3.118, df=4, p=0.0356<br>[-0.9 s ; 0 s] Late<br>t=2.035, df=4, p=0.1116<br>[0.1 s ; 1 s] Early<br>t=3.017, df=4, p=0.0393<br>[0.1 s ; 1 s] Late<br>t=1.331, df=4, p=0.2540<br>[1.1 s ; 2 s] Early<br>t=2.116, df=4, p=0.1018<br>[1.1 s ; 2 s] Late<br>t=1.491, df=4, p= 0.2103 |
| Figure 4.I (bottom left) | One-sample t-test against 0<br><br>One-sample t-test against 0<br><br>Paired t-test (two-tailed) | GluN1-C1 Control peptide Early / Late = 5 | Peak Z-score | [-0.9 s ; 0 s] Early<br>t=3.315, df=4, p= 0.0295<br><br>[-0.9 s ; 0 s] Late<br>t=2.666, df=4, p= 0.0560<br><br>[-0.9 s ; 0 s]<br>t=0.1031, df=4, p= 0.9228 |
| Figure 4.I (bottom right) | One-sample t-test against 0<br><br>One-sample t-test against 0<br><br>Paired t-test (two-tailed) | GluN1-C1 Control peptide Early / Late = 5 | Peak Z-score | [0.1 s ; 1 s] Early<br>t=3.911, df=4, p= 0.0174<br><br>[0.1 s ; 1 s] Late<br>t=2.241, df=4, p= 0.0886<br><br>[0.1 s ; 1 s]<br>t=1.799, df=4, p= 0.1465 |
| Figure 4.K (top) | Two-Factor RM ANOVA | D2R- IL3 Control peptide Early / Late = 8 | AUC | Time Effect F(2, 14) = 6.114, p=0.0123<br>Training Effect F(1, 7) = 2.637, p=0.1484<br>Time x Training F(2, 14)=8.324, p=0.0042 |

| | | | | |
|---|---|---|---|---|
| | | | | *Tukey's multiple comparisons test*<br><br>[-0.9;0]<br>Early vs. Late p=0.6539<br>[0.1;1]<br>Early vs. Late p=0.0001<br>[1.1;2]<br>Early vs. Late p=0.0160<br><br>Early<br>[-0.9;0] vs. [0.1;1] p=0.0128<br>[-0.9;0] vs. [1.1;2] p=0.4103<br>[0.1;1] vs. [1.1;2] p=0.0010<br>Late<br>[-0.9;0] vs. [0.1;1] p=0.0710<br>[-0.9;0] vs. [1.1;2] p=0.0013<br>[0.1;1] vs. [1.1;2] p=0.1285 |
| Figure 4.K (top) | One-sample t-test against 0 | D2R- IL3 Control peptide Early / Late = 8 | AUC | [-0.9 s ; 0 s] Early<br>t=3.907, df=7, p=0.0058<br>[-0.9 s ; 0 s] Late<br>t=2.790, df=7, p=0.0269<br>[0.1 s ; 1 s] Early<br>t=7.192, df=7, p=0.0002<br>[0.1 s ; 1 s] Late<br>t=1.661, df=7, p=0.1406<br>[1.1 s ; 2 s] Early<br>t=2.976, df=7, p=0.0206<br>[1.1 s ; 2 s] Late<br>t=0.005791, df=7, p= 0.9955 |
| Figure 4.K (bottom left) | One-sample t-test against 0<br><br>One-sample t-test against 0<br><br>Paired t-test (two-tailed) | D2R- IL3 Control peptide Early / Late = 8 | Peak Z-score | [-0.9 s ; 0 s] Early<br>t=6.370, df=7, p= 0.0004<br><br>[-0.9 s ; 0 s] Late<br>t=3.113, df=7, p= 0.0170<br><br>[-0.9 s ; 0 s]<br>t=0.2048, df=7, p= 0.8436 |
| Figure 4.K (bottom right) | One-sample t-test against 0<br><br>One-sample t-test against 0<br><br>Paired t-test (two-tailed) | D2R- IL3 Control peptide Early / Late = 8 | Peak Z-score | [0.1 s ; 1 s] Early<br>t=9.671, df=7, p <0.0001<br><br>[0.1 s ; 1 s] Late<br>t=3.675, df=7, p= 0.0079<br><br>[0.1 s ; 1 s]<br>t=1.958, df=7, p= 0.0911 |
| Figure 4.M (top) | Two-Factor RM ANOVA | GluN1-C1 Control peptide RL / UL = 5 | AUC | Time Effect F(2, 8) = 3.665, p=0.0742<br>Lever Effect F(1, 4) = 1.573, p=0.2780<br>Time x Lever F(2, 8) = 8.469, p=0.0106<br><br>*Tukey's multiple comparisons test*<br><br>[-0.9;0]<br>RL vs. UL p=0.3155<br>[0.1;1]<br>RL vs. UL p=0.0015<br>[1.1;2]<br>RL vs. UL p=0.0380<br><br>RL |

| | | | | |
|---|---|---|---|---|
| | | | | [-0.9;0] vs. [0.1;1] p=0.0054<br>[-0.9;0] vs. [1.1;2] p=0.7339<br>[0.1;1] vs. [1.1;2] p=0.0021<br>UL<br>[-0.9;0] vs. [0.1;1] p=0.4214<br>[-0.9;0] vs. [1.1;2] p=0.0064<br>[0.1;1] vs. [1.1;2] p=0.0411 |
| Figure 4.M (top) | One-sample t-test against 0 | GluN1-C1 Control peptide RL / UL = 5 | AUC | [-0.9 s ; 0 s] RL<br>t=2.011, df=4, p=0.1146<br>[-0.9 s ; 0 s] UL<br>t=1.455, df=4, p=0.2194<br>[0.1 s ; 1 s] RL<br>t=4.715, df=4, p=0.0092<br>[0.1 s ; 1 s] UL<br>t=0.7353, df=4, p=0.5029<br>[1.1 s ; 2 s] RL<br>t=1.394, df=4, p=0.2357<br>[1.1 s ; 2 s] UL<br>t=0.3488, df=4, p= 0.7448 |
| Figure 4.M (bottom left) | One-sample t-test against 0<br><br>One-sample t-test against 0<br><br>Paired t-test (two-tailed) | GluN1-C1 Control peptide RL / UL = 5 | Peak Z-score | [-0.9 s ; 0 s] RL<br>t=3.044, df=4, p= 0.0383<br><br>[-0.9 s ; 0 s] UL<br>t=2.048, df=4, p= 0.1100<br><br>[-0.9 s ; 0 s]<br>t=0.6026, df=4, p= 0.5793 |
| Figure 4.M (bottom right) | One-sample t-test against 0<br><br>One-sample t-test against 0<br>Paired t-test (two-tailed) | GluN1-C1 Control peptide RL / UL = 5 | Peak Z-score | [0.1 s ; 1 s] RL<br>t=5.835, df=4, p= 0.0043<br><br>[0.1 s ; 1 s] UL<br>t=1.407, df=4, p= 0.2323<br><br>[0.1 s ; 1 s]<br>t=2.335, df=4, p= 0.0798 |
| Figure 4.O (top) | Two-Factor RM ANOVA | D2R-IL3 Control peptide RL = 8 | AUC | Time Effect F(2, 14) = 4.815, p=0.0256<br>Lever Effect F(1, 7) = 1.592, p=0.2474<br>Time x Lever F(2, 14) = 4.004, p=0.0422<br><br>*Tukey's multiple comparisons test*<br><br>[-0.9;0]<br>RL vs. UL p=0.5563<br>[0.1;1]<br>RL vs. UL p=0.0164<br>[1.1;2]<br>RL vs. UL p=0.2241<br><br>RL<br>[-0.9;0] vs. [0.1;1] p=0.4333<br>[-0.9;0] vs. [1.1;2] p=0.0452<br>[0.1;1] vs. [1.1;2] p=0.0039<br>UL<br>[-0.9;0] vs. [0.1;1] p=0.6773<br>[-0.9;0] vs. [1.1;2] p=0.7111<br>[0.1;1] vs. [1.1;2] p=0.9982 |
| Figure 4.O (top) | One-sample t-test against 0 | D2R- IL3 Control peptide UL = 8 | AUC | [-0.9 s ; 0 s] RL<br>t=3.530, df=7, p=0.0096<br>[-0.9 s ; 0 s] UL<br>t=4.261, df=7, p=0.0037 |

| | | | | |
|---|---|---|---|---|
| | | | | [0.1 s ; 1 s] RL<br>t=3.903, df=7, p=0.0059<br>[0.1 s ; 1 s] UL<br>t=3.254, df=7, p=0.0140<br>[1.1 s ; 2 s] RL<br>t=0.3703, df=7, p=0.7221<br>[1.1 s ; 2 s] UL<br>t=2.604, df=7, p= 0.0352 |
| Figure 4.O (bottom left) | One-sample t-test against 0<br><br>One-sample t-test against 0<br><br>Paired t-test (two-tailed) | D2R- IL3 Control peptide RL / UL = 8 | Peak Z-score | [-0.9 s ; 0 s] RL<br>t=4.362, df=7, p= 0.0033<br><br>[-0.9 s ; 0 s] UL<br>t=4.429, df=7, p= 0.0030<br><br>[-0.9 s ; 0 s]<br>t=3.383, df=7, p= 0.0117 |
| Figure 4.O (bottom right) | One-sample t-test against 0<br><br>One-sample t-test against 0<br><br>Paired t-test (two-tailed) | D2R- IL3 Control peptide RL / UL = 8 | Peak Z-score | [0.1 s ; 1 s] RL<br>t=5.146, df=7, p= 0.0013<br><br>[0.1 s ; 1 s] UL<br>t=7.247, df=7, p= 0.0002<br><br>[0.1 s ; 1 s]<br>t=3.056, df=7, p= 0.0184 |
| Figure 4.Q (top) | Two-Factor RM ANOVA | GluN1-C1 Control peptide RL / UL = 5 | AUC | Time Effect F(2, 8) = 6.300, p=0.0227<br>Lever Effect F(1, 4) = 0.03752, p=0.8559<br>Time x Lever F(2, 8) = 1.378, p=0.3060 |
| Figure 4.Q (top) | One-sample t-test against 0 | GluN1-C1 Control peptide RL / UL = 5 | AUC | [-0.9 s ; 0 s] RL<br>t=0.1241, df=4, p=0.9073<br>[-0.9 s ; 0 s] UL<br>t=0.2328, df=4, p=0.8273<br>[0.1 s ; 1 s] RL<br>t=0.03377, df=4, p=0.9747<br>[0.1 s ; 1 s] UL<br>t=0.7374, df=4, p=0.5018<br>[1.1 s ; 2 s] RL<br>t=3.992, df=4, p=0.0162<br>[1.1 s ; 2 s] UL<br>t=1.722, df=4, p=0.1601 |
| Figure 4.Q (top insert) | One-sample t-test against 0<br><br>One-sample t-test against 0<br><br>Paired t-test (two-tailed) | GluN1-C1 Control peptide RL / UL = 10 | AUC | RL<br>t=0.02388, df=9, p= 0.9815<br><br>UL<br>t=0.4067, df=9, p= 0.6937<br><br>RL vs. UL<br>t=0.6899, df=9, p= 0.5077 |
| Figure 4.Q (bottom left) | One-sample t-test against 0<br><br>One-sample t-test against 0<br><br>Paired t-test (two-tailed) | GluN1-C1 Control peptide RL / UL = 5 | Peak Z-score | [-0.9 s ; 0 s] RL<br>t=3.044, df=4, p= 0.0383<br><br>[-0.9 s ; 0 s] UL<br>t=1.443, df=4, p= 0.2224<br><br>[-0.9 s ; 0 s]<br>t=0.1148, df=4, p= 0.9141 |
| Figure 4.Q (bottom right) | One-sample t-test against 0<br><br>One-sample t-test against 0 | GluN1-C1 Control peptide RL / UL = 5 | Peak Z-score | [0.1 s ; 1 s] RL<br>t=2.114, df=4, p= 0.1020<br><br>[0.1 s ; 1 s] UL<br>t=0.5837, df=4, p= 0.5908 |

| | | | | |
|---|---|---|---|---|
| | Paired t-test (two-tailed) | | | [0.1 s ; 1 s]<br>t=1.060, df=4, p= 0.3487 |
| Figure 4.S (top) | Two-Factor RM ANOVA | D2R- IL3 Control peptide RL / UL = 8 | AUC | Time Effect F(2, 14) = 8.812, p=0.0033<br>Lever Effect F(1, 7) = 9.748, p=0.0168<br>Time x Lever F(2, 14) = 0.1847, p=0.8333 |
| Figure 4.S (top) | One-sample t-test against 0 | D2R- IL3 Control peptide RL / UL = 8 | AUC | [-0.9 s ; 0 s] RL<br>t=0.8607, df=7, p=0.4179<br>[-0.9 s ; 0 s] UL<br>t=2.094, df=7, p=0.0745<br>[0.1 s ; 1 s] RL<br>t=0.8229, df=7, p=0.4377<br>[0.1 s ; 1 s] UL<br>t=1.288, df=7, p=0.2388<br>[1.1 s ; 2 s] RL<br>t=2.750, df=7, p=0.0285<br>[1.1 s ; 2 s] UL<br>t=0.3162, df=7, p= 0.7611 |
| Figure 4.S (top insert) | One-sample t-test against 0<br><br>One-sample t-test against 0<br><br>Paired t-test (two-tailed) | D2R- IL3 Control peptide RL / UL = 16 | AUC | RL<br>t=1.218, df=15, p= 0.2421<br><br>UL<br>t=2.427, df=15, p= 0.0283<br><br>RL vs. UL<br>t=2.937, df=15, p= 0.0102 |
| Figure 4.S (bottom left) | One-sample t-test against 0<br><br>One-sample t-test against 0<br><br>Paired t-test (two-tailed) | D2R- IL3 Control peptide RL / UL = 8 | Peak Z-score | [-0.9 s ; 0 s] RL<br>t=2.334, df=7, p= 0.0523<br><br>[-0.9 s ; 0 s] UL<br>t=3.052, df=7, p= 0.0185<br><br>[-0.9 s ; 0 s]<br>t=1.859, df=7, p= 0.1054 |
| Figure 4.S (bottom right) | One-sample t-test against 0<br><br>One-sample t-test against 0<br><br>Paired t-test (two-tailed) | D2R- IL3 Control peptide RL / UL = 8 | Peak Z-score | [0.1 s ; 1 s] RL<br>t=1.367, df=7, p= 0.2140<br><br>[0.1 s ; 1 s] UL<br>t=2.660, df=7, p= 0.0325<br><br>[0.1 s ; 1 s]<br>t=1.710, df=7, p= 0.1310 |
| Figure 5 Peptide mPFC cpt | | | | |
| Figure 5.D | Two-Factor RM ANOVA | Control Peptide = 30<br>C1 peptide = 21<br>IL3 peptide = 21 | Lever Presses / minute | Peptide Effect F(2,69) = 2.926, p=0.0603<br>Training Effect F(3,207) = 542.8, p<0.0001<br>Interaction F(6,207) = 1.263, p=0.2761 |
| Figure 5.E | Planned Orthogonal Contrast<br><br>Two-Factor RM ANOVA | Control Peptide = 30<br>C1 peptide = 21<br>IL3 peptide = 21 | Lever presses | Peptide effect F(2,69) = 1.117, p=0.3331<br>Devaluation effect F(1,69) = 75.444, p<0.0001<br>Interaction F(2,69) = 0.890, p=0.4153 |
| Figure 5.F | Planned Orthogonal Contrast<br><br>Single-Factor ANOVA | Control Peptide = 30<br>C1 peptide = 21<br>IL3 peptide = 21 | Devaluation Ratio | Interaction F(2,69) = 3.128, p=0.0501<br>Multiple comparison<br>Control vs GluN1-C1 F(2,69) = 0.352, p=0.7045<br>Control vs D2R-IL3 F(2,69) = 5.660, p=0.0053 |

| | | | | |
|---|---|---|---|---|
| | | | | GluN1-C1 vs D2R-IL3 F(2,69) = 2.711, p=0.0736 |
| Figure 5.F | One-sample t-test against 0.5 | Control Peptide = 30 | Devaluation Ratio | t=4.172, df=29, p=0.0003 |
| Figure 5.F | One-sample t-test against 0.5 | C1 peptide = 21 | Devaluation Ratio | t=5.820, df=20, p<0.0001 |
| Figure 5.F | One-sample t-test against 0.5 | IL3 peptide = 21 | Devaluation Ratio | t=5,511, df=20, p<0.0001 |
| Figure 5.G | Two-Factor RM ANOVA | Control Peptide = 28<br>C1 peptide = 21<br>IL3 peptide = 20 | Lever Presses / minute | Peptide Effect F(2,66) = 2.747, p=0.0715<br>Training Effect F(3,198) = 2.728, p=0.0452<br>Interaction F(6,198) = 1.176, p=0.3204 |
| Figure 5.H | Planned Orthogonal Contrast<br><br>Two-Factor RM ANOVA | Control Peptide = 28<br>C1 peptide = 21<br>IL3 peptide = 20 | Lever presses | Peptide effect F(2,66) =3.773, p=0.0281<br>Devaluation effect F(1,66) =6.955, p=0.0104<br>Interaction F(2,66) =3.939, p=0.0242<br>Within group multiple comparison<br>Control F(1,66) = 11.664, p=0.0011,<br>GluN1-C1 F(1,66) = 1.337, p=0.2517,<br>D2R-IL3 F(1,66) = 0.085, p=0.7715 |
| Figure 5.I | Planned Orthogonal Contrast<br><br>Single-Factor ANOVA | Control Peptide = 28<br>C1 peptide = 21<br>IL3 peptide = 20 | Devaluation Ratio | Interaction F(2,66) =0.336, p=0.7158 |
| Figure 5.I | One-sample t-test against 0.5 | Control Peptide = 28 | Devaluation Ratio | t=2.908, df=27, p=0.0072 |
| Figure 5.I | One-sample t-test against 0.5 | C1 peptide = 21 | Devaluation Ratio | t=0.8665, df=20, p=0.3965 |
| Figure 5.I | One-sample t-test against 0.5 | IL3 peptide = 20 | Devaluation Ratio | t=1.206, df=19, p=0.2426 |
| Figure 5.D | Two-Factor RM ANOVA | Control Peptide = 30<br>C1 peptide = 21<br>IL3 peptide = 21 | Lever Presses / minute | Peptide Effect F(2,69) = 2.926, p=0.0603<br>Training Effect F(3,207) = 542.8, p<0.0001<br>Interaction F(6,207) = 1.263, p=0.2761 |
| Figure 5.E | Planned Orthogonal Contrast<br><br>Two-Factor RM ANOVA | Control Peptide = 30<br>C1 peptide = 21<br>IL3 peptide = 21 | Lever presses | Peptide effect F(2,69) = 1.117, p=0.3331<br>Devaluation effect F(1,69) = 75.444, p<0.0001<br>Interaction F(2,69) = 0.890, p=0.4153 |
| Figure 5.F | Planned Orthogonal Contrast<br><br>Single-Factor ANOVA | Control Peptide = 30<br>C1 peptide = 21<br>IL3 peptide = 21 | Devaluation Ratio | Interaction F(2,69) = 3.128, p=0.0501<br>Multiple comparison<br>Control vs GluN1-C1 F(2,69) = 0.352, p=0.7045<br>Control vs D2R-IL3 F(2,69) = 5.660, p=0.0053<br>GluN1-C1 vs D2R-IL3 F(2,69) = 2.711, p=0.0736 |
| Figure 5.F | One-sample t-test against 0.5 | Control Peptide = 30 | Devaluation Ratio | t=4.172, df=29, p=0.0003 |
| Figure 5.F | One-sample t-test against 0.5 | C1 peptide = 21 | Devaluation Ratio | t=5.820, df=20, p<0.0001 |
| Figure 5.F | One-sample t-test against 0.5 | IL3 peptide = 21 | Devaluation Ratio | t=5,511, df=20, p<0.0001 |

| | | | | |
|---|---|---|---|---|
| Figure 5.G | Two-Factor RM ANOVA | Control Peptide = 28<br>C1 peptide = 21<br>IL3 peptide = 20 | Lever Presses / minute | Peptide Effect F(2,66) = 2.747, p=0.0715<br>Training Effect F(3,198) = 2.728, p=0.0452<br>Interaction F(6,198) = 1.176, p=0.3204 |
| Figure 5.H | Planned Orthogonal Contrast<br><br>Two-Factor RM ANOVA | Control Peptide = 28<br>C1 peptide = 21<br>IL3 peptide = 20 | Lever presses | Peptide effect F(2,66) =3.773, p=0.0281<br>Devaluation effect F(1,66) =6.955, p=0.0104<br>Interaction F(2,66) =3.939, p=0.0242<br>Within group multiple comparison<br>Control F(1,66) = 11.664, p=0.0011,<br>GluN1-C1 F(1,66) = 1.337, p=0.2517,<br>D2R-IL3 F(1,66) = 0.085, p=0.7715 |
| Figure 5.I | Planned Orthogonal Contrast<br><br>Single-Factor ANOVA | Control Peptide = 28<br>C1 peptide = 21<br>IL3 peptide = 20 | Devaluation Ratio | Interaction F(2,66) =0.336, p=0.7158 |
| Figure 5.I | One-sample t-test against 0.5 | Control Peptide = 28 | Devaluation Ratio | t=2.908, df=27, p=0.0072 |
| Figure 5.I | One-sample t-test against 0.5 | C1 peptide = 21 | Devaluation Ratio | t=0.8665, df=20, p=0.3965 |
| Figure 5.I | One-sample t-test against 0.5 | IL3 peptide = 20 | Devaluation Ratio | t=1.206, df=19, p=0.2426 |
| Supplementary Figure 1 - DAT Gi VTA | | | | |
| Suppl. Fig. 1.B | Two-way RM ANOVA | mCherry = 10<br>DREADD-Gi = 10 | Lever presses / min | Training Effect F(3, 54) = 102.6, p<0,0001<br>Group Effect F(1, 18) = 0.2437, p=0.6275<br>Training x Group F(3, 54) = 0.1860, p=0.9055 |
| Suppl. Fig. 1.C | Unpaired t-test | mCherry = 9<br>DREADD-Gi = 10 | Food Intake | t=0.8623, df=17, p=0.4005 |
| Suppl. Fig. 1.D | Two-way RM ANOVA | mCherry = 9<br>DREADD-Gi = 10 | Food Intake | Devaluation Effect F(1, 17) = 14.88, p=0.0013<br>Group Effect F(1, 17) = 0.6737, p=0.4231<br>Devaluation x Group F(1, 17) = 0.6262, p=0.4397 |
| Suppl. Fig. 1.E | Unpaired t-test | Cre- = 11<br>Cre + = 11 | Food Intake | t=0.3720, df=20, p=0.7138 |
| Suppl. Fig. 1.F | Two-way RM ANOVA | Cre- = 11<br>Cre + = 11 | Food Intake | Devaluation Effect F(1, 20) = 37.74, p<0.0001<br>Group Effect F(1, 20) = 5.299, p=0.0322<br>Devaluation x Group F(1, 20) = 0.3203, p=0.5777 |
| Suppl. Fig. 1.G | Welch's t-test (Populations have different SDs) | Cre- = 11<br>Cre + = 11 | Food Intake | t=2.302, df=14,48, p=0.0367 |
| Suppl. Fig. 1.H | Unpaired t-test | Cre- = 10<br>Cre + = 11 | Food Intake | t=1,291, df=19, p=0.2123 |
| Suppl. Fig. 1.I | Two-way RM ANOVA | Cre- = 10<br>Cre + = 11 | Food Intake | Devaluation Effect F(1, 19)=103.8, p<0.0001<br>Group Effect F(1, 19) = 2.600, p=0.1233<br>Devaluation x Group F(1, 19) = 0.5850, p=0.4537 |
| Supplementary Figure 2B – GRAB DA CPT | | | | |
| Suppl. Fig. 2B.A | RM One-way ANOVA | n=9 | Latency to consumption | F(9, 72) = 6.385, p<0.0001<br>*Tukey's multiple comparisons test*<br>1 vs. 2 p<0.0001<br>1 vs. 10 p<0.0001 |

| | | | | |
|---|---|---|---|---|
| | | | | 2 vs. 10 p = 0.9989 |
| Suppl. Fig. 2B.B | RM One-way ANOVA | n=9 | Reward number | F(7, 56) = 63.70, p<0.0001<br>*Dunnett's multiple comparisons test*<br>1 vs. 8 p<0.0001 |
| Suppl. Fig. 2B.C | Two-way RM ANOVA | n=9 | Lever presses | Session Effect F (7, 56) = 12.74 , p<0.0001<br>Lever Effect F (1, 8) = 66.81, p<0.0001<br>Session x Lever F (7, 56) = 24.20, p<0.0001<br>*Šídák's multiple comparisons test*<br>UL - RL<br>1 p=0.9957<br>2 p<0.0001<br>3 p<0.0001<br>4 p<0.0001<br>5 p<0.0001<br>6 p<0.0001<br>7 p<0.0001<br>8 p<0.0001 |
| Suppl. Fig. 2B.D | Pearson's correlation coefficient<br><br>Simple linear regression | n=9 | Correlation between AUC (RL-UL) difference and lever preference index | r(7) = 0.5514, p=0.1238<br><br>$r^2$ = 0.3041, F (1, 7) = 3.058, p = 0.1238 |
| Suppl. Fig. 2B.E | RM One-way ANOVA | n=9 | Reward number | F (7, 56) = 47.14, p<0.0001<br>*Dunnett's multiple comparisons test*<br>1 vs. 8, p<0.0001 |
| Suppl. Fig. 2B.F | Two-way RM ANOVA | n=9 | Lever presses | Session Effect F(7, 56) = 30.90, p<0.0001<br>Lever Effect F(1, 8) = 19.07, p=0.0024<br>Session x Lever F(7, 56)= 12.08, p<0.0001<br>*Šídák's multiple comparisons test*<br>UL – RL<br>1 p<0.0001<br>2 p=0.1716<br>3 p=0.3386<br>4 p=0.0114<br>5 p=0.0121<br>6 p=0.0022<br>7 p=0.0003<br>8 p=0.0009 |
| Suppl. Fig. 2B.G | Pearson's correlation coefficient<br><br>Simple linear regression | n=9 | Correlation between AUC (RL-UL) difference and lever preference index | r(7) = 0.8560, p=0.0032<br><br>$r^2$ = 0.7327, F (1, 7) = 19.19, p = 0.0032 |
| Supplementary Figure 3A – PLA | | | | |
| Suppl. Fig. 3A.B | Kruskal-Wallis test | n = 517 fields / 4 mice | D1/NMDA PLA points x $10^3$ / $mm^2$ | H(8) = 218.4, p<0.0001<br><br>*Dunn's multiple comparisons test*<br><br>DLS vs. DMS p>0.9999 |

| | | | | |
|---|---|---|---|---|
| | | | | DLS vs. Core p<0.0001<br>DLS vs. medShell p<0.0001<br>DLS vs. latShell p<0.0001<br>DLS vs. ACC p<0.0001<br>DLS vs. PL p<0.0001<br>DLS vs. IL p<0.0001<br>DLS vs. MO p<0.0001<br>DMS vs. Core p<0.0001<br>DMS vs. medShell p<0.0001<br>DMS vs. latShell p=0.0004<br>DMS vs. ACC p<0.0001<br>DMS vs. PL p<0.0001<br>DMS vs. IL p<0.0001<br>DMS vs. MO p<0.0001<br>Core vs. medShell p>0.9999<br>Core vs. latShell p>0.9999<br>Core vs. ACC p=0.1021<br>Core vs. PL p=0.0048<br>Core vs. IL p=0.0263<br>Core vs. MO p=0.8736<br>medShell vs. latShell p>0.9999<br>medShell vs. ACC p=0.4466<br>medShell vs. PL p=0.0322<br>medShell vs. IL p=0.1149<br>medShell vs. MO p>0.9999<br>latShell vs. ACC p=0.0037<br>latShell vs. PL p<0.0001<br>latShell vs. IL p=0.0011<br>latShell vs. MO p=0.0819<br>ACC vs. PL p>0.9999<br>ACC vs. IL p>0.9999<br>ACC vs. MO p>0.9999<br>PL vs. IL p>0.9999<br>PL vs. MO p>0.9999<br>IL vs. MO p>0.9999 |
| Suppl. Fig. 3A.C | Kruskal-Wallis test | n = 918 fields / 6 mice | D2/NMDA PLA points x $10^3$ / $mm^2$ | $H(8) = 99.49$, $p<0.0001$<br><br>*Dunn's multiple comparisons test*<br><br>DLS vs. DMS p>0.9999<br>DLS vs. Core p<0.0001<br>DLS vs. medShell p>0.9999<br>DLS vs. latShell p>0.9999<br>DLS vs. ACC p>0.9999<br>DLS vs. PL p>0.9999<br>DLS vs. IL p<0.0001<br>DLS vs. MO p>0.9999<br>DMS vs. Core p<0.0001<br>DMS vs. medShell p=0.0609<br>DMS vs. latShell p>0.9999<br>DMS vs. ACC p>0.9999<br>DMS vs. PL p>0.9999<br>DMS vs. IL p=0.0113 |

| | | | | |
|---|---|---|---|---|
| | | | | DMS vs. MO p>0.9999<br>Core vs. medShell p=0.2933<br>Core vs. latShell p=0.0035<br>Core vs. ACC p<0.0001<br>Core vs. PL p=0.0001<br>Core vs. IL p<0.0001<br>Core vs. MO p=0.0005<br>medShell vs. latShell p>0.9999<br>medShell vs. ACC p>0.9999<br>medShell vs. PL p>0.9999<br>medShell vs. IL p<0.0001<br>medShell vs. MO p>0.9999<br>latShell vs. ACC p>0.9999<br>latShell vs. PL p>0.9999<br>latShell vs. IL p<0.0001<br>latShell vs. MO p>0.9999<br>ACC vs. PL p>0.9999<br>ACC vs. IL p<0.0001<br>ACC vs. MO p>0.9999<br>PL vs. IL p<0.0001<br>PL vs. MO p>0.9999<br>IL vs. MO p<0.0001 |
| Supplementary Figure 3B – D1D2 GCAMP Peptide CPT | | | | |
| Suppl. Fig. 3B.A | Two-way RM ANOVA | GluN1-Δ = 5 / GluN1-C1 = 5 | Reward / min | Session Effect F(11, 88) = 1.409,p=0.1833<br>Peptide Effect F(1, 8) = 1.369, p=0.2757<br>Session x Peptide F(11, 88)=1.201, p=0.2981 |
| Suppl. Fig. 3B.B | Two-way RM ANOVA | D2R-Scr = 6 / D2R-IL3 = 8 | Reward / min | Session Effect F(11, 132) = 2.862,p=0.0021<br>Peptide Effect F(1, 12) = 1.328, p=0.2716<br>Session x Peptide F (11, 132) = 0.5762, p=0.8454 |
| Suppl. Fig. 3B.C | Two-way RM ANOVA | GluN1-Δ = 5 / GluN1-C1 = 5 | Reward / min | Session Effect F(7, 56) = 11.42,p<0.0001<br>Peptide Effect F(1, 8) = 0.1432, p=0.7150<br>Session x Peptide F (7, 56) = 0.6472, p=0.7150 |
| Suppl. Fig. 3B.D | Two-way RM ANOVA | D2R-Scr = 6 / D2R-IL3 = 8 | Reward / min | Session Effect F(7, 84) = 21.30, p<0.0001<br>Peptide Effect F(1, 12) = 1.475, p=0.2479<br>Session x Peptide F(7, 84)=0.5982, p=0.7557 |
| Suppl. Fig. 3B.E | Two-way RM ANOVA | GluN1-Δ = 5 / GluN1-C1 = 5 | Reward / min | Session Effect F(7, 56) = 23.37,p<0.0001<br>Peptide Effect F(1, 8) = 3.229, p=0.1101<br>Session x Peptide F (7, 56) = 0.8945, p=0.5172 |
| Suppl. Fig. 3B.F | Two-way RM ANOVA | D2R-Scr = 6 / D2R-IL3 = 8 | Reward / min | Session Effect F(7, 84) = 21.30, p<0.0001<br>Peptide Effect F(1, 12) = 1.475, p=0.2479<br>Session x Peptide F(7, 84)=0.5982, p=0.7557 |
| Suppl. Fig. 3B.G | Two-way RM ANOVA | GluN1-Δ = 5 / GluN1-C1 = 5 | Lever Preference Index | Session F (7, 56) = 47.43, p<0.0001<br>Peptide F (1, 8) = 4.025, p=0.0797<br>Session x Peptide F (7, 56) = 0.1439, p=0.9941 |
| Suppl. Fig. 3B.H | Two-way RM ANOVA | D2R-Scr = 6 / D2R-IL3 = 8 | Lever Preference Index | Session F (7, 84) = 64.57, p<0.0001<br>Peptide F (1, 12) = 3.238, p=0.0971<br>Session x Peptide F (7, 84) = 1.019, p=0.4240 |
| Supplementary Figure 4 – D1D2 GCAMP Peptide Licks | | | | |
| Suppl. Fig. 4.B | One sample t-test | GluN1-Δ = 5 | AUC | [-0.9 s ; 0 s] Early<br>t=9.672, df=4, p=0.0006<br><br>[-0.9 s ; 0 s] Late |

| | | | | |
|---|---|---|---|---|
| | One sample t-test<br><br>Paired t-test (two tailed) | | | t=2.670, df=4, p=0.0558<br><br>[-0.9 s ; 0 s] Early vs. Late<br>t=3.098, df=4, p=0.0363 |
| Suppl. Fig. 4.B | One sample t-test<br><br>One sample t-test<br><br>Paired t-test (two tailed) | GluN1-Δ = 5 | AUC | [0.1 s ; 1 s] Early<br>t=4.647, df=4, p=0.0097<br><br>[0.1 s ; 1 s] Late<br>t=2.206, df=4 p=0.0920<br><br>[0.1 s ; 1 s] Early vs. Late<br>t=3.686, df=4, p=0.0211 |
| Suppl. Fig. 4.D | One sample t-test<br><br>One sample t-test<br><br>Paired t-test (two tailed) | GluN1-C1 = 5 | AUC | [-0.9 s ; 0 s] Early<br>t=6.926, df=4, p=0.0023<br><br>[-0.9 s ; 0 s] Late<br>t=5.351, df=4, p=0.0059<br><br>[-0.9 s ; 0 s] Early vs. Late<br>t=0.3583, df=4, p=0.7383 |
| Suppl. Fig. 4.D | One sample t-test<br><br>One sample t-test<br><br>Paired t-test (two tailed) | GluN1-C1 = 5 | AUC | [0.1 s ; 1 s] Early<br>t=4.991, df=4, p=0.0075<br><br>[0.1 s ; 1 s] Late<br>t=5.711, df=4, p=0.0046<br><br>[0.1 s ; 1 s] Early vs. Late<br>t=1.356, df=4, p=0.2466 |
| Suppl. Fig. 4.F | One sample t-test<br><br>One sample t-test<br><br>Paired t-test (two tailed) | D2R-Scr = 6 | AUC | [-0.9 s ; 0 s] Early<br>t=2.971, df=5, p=0.0311<br><br>[-0.9 s ; 0 s] Late<br>t=4.946, df=5, p=0.0043<br><br>[-0.9 s ; 0 s] Early vs. Late<br>t=0.2469, df=5, p=0.8148 |
| Suppl. Fig. 4.F | One sample t-test<br><br>One sample t-test<br><br>Paired t-test (two tailed) | D2R-Scr = 6 | AUC | [0.1 s ; 1 s] Early<br>t=4.797, df=5, p=0.0049<br><br>[0.1 s ; 1 s] Late<br>t=5.659, df=5, p=0.0024<br><br>[0.1 s ; 1 s] Early vs. Late<br>t=1.090, df=5, p=0.3254 |
| Suppl. Fig. 4.H | One sample t-test<br><br>One sample t-test<br><br>Paired t-test (two tailed) | D2R-IL3 = 8 | AUC | [-0.9 s ; 0 s] Early<br>t=4.740, df=7, p=0.0021<br><br>[-0.9 s ; 0 s] Late<br>t=5.765, df=7, p=0.0007<br><br>[-0.9 s ; 0 s] Early vs. Late<br>t=3.410, df=7, p=0.0113 |
| Suppl. Fig. 4.H | One sample t-test<br><br>One sample t-test | D2R-IL3 = 8 | AUC | [0.1 s ; 1 s] Early<br>t=4.456, df=7, p=0.0029<br><br>[0.1 s ; 1 s] Late<br>t=4.049, df=7, p=0.0049<br><br>[0.1 s ; 1 s] Early vs. Late |

| | | | | |
|---|---|---|---|---|
| | Paired t-test (two tailed) | | | t=3.129, df=7, p=0.0166 |
| Suppl. Fig. 4.J | One sample t-test<br><br>One sample t-test<br><br>Paired t-test (two tailed) | GluN1-Δ = 5 | AUC | [-0.9 s ; 0 s] Early<br>t=1.719, df=4, p=0.1608<br><br>[-0.9 s ; 0 s] Late<br>t=1.705, df=4, p=0.1634<br><br>[-0.9 s ; 0 s] Early vs. Late<br>t=1.494, df=4, p=0.2094 |
| Suppl. Fig. 4.J | One sample t-test<br><br>One sample t-test<br><br>Paired t-test (two tailed) | GluN1-Δ = 5 | AUC | [0.1 s ; 1 s] Early<br>t=2.144, df=4, p=0.0986<br><br>[0.1 s ; 1 s] Late<br>t=1.390, df=4, p=0.2368<br><br>[0.1 s ; 1 s] Early vs. Late<br>t=2.925, df=4, p=0.0430 |
| Suppl. Fig. 4.L | One sample t-test<br><br>One sample t-test<br><br>Paired t-test (two tailed) | GluN1-C1 = 5 | AUC | [-0.9 s ; 0 s] Early<br>t=5.278, df=4, p=0.0062<br><br>[-0.9 s ; 0 s] Late<br>t=4.929, df=4, p=0.0079<br><br>[-0.9 s ; 0 s] Early vs. Late<br>t=3.702, df=4, p=0.0208 |
| Suppl. Fig. 4.L | One sample t-test<br><br>One sample t-test<br><br>Paired t-test (two tailed) | GluN1-C1 = 5 | AUC | [0.1 s ; 1 s] Early<br>t=4.019, df=4, p=0.0159<br><br>[0.1 s ; 1 s] Late<br>t=6.913, df=4, p=0.0023<br><br>[0.1 s ; 1 s] Early vs. Late<br>t=2.374, df=4, p=0.0765 |
| Suppl. Fig. 4.N | One sample t-test<br><br>One sample t-test<br><br>Paired t-test (two tailed) | D2R-Scr = 6 | AUC | [-0.9 s ; 0 s] Early<br>t=3.426, df=5, p=0.0187<br><br>[-0.9 s ; 0 s] Late<br>t=3.173, df=5, p=0.0247<br><br>[-0.9 s ; 0 s] Early vs. Late<br>t=0.2621, df=5, p=0.8037 |
| Suppl. Fig. 4.N | One sample t-test<br><br>One sample t-test<br><br>Paired t-test (two tailed) | D2R-Scr = 6 | AUC | [0.1 s ; 1 s] Early<br>t=4.395, df=5, p=0.0071<br><br>[0.1 s ; 1 s] Late<br>t=3.522, df=5, p=0.0169<br><br>[0.1 s ; 1 s] Early vs. Late<br>t=1.091, df=5, p=0.3249 |
| Suppl. Fig. 4.P | One sample t-test<br><br>One sample t-test<br><br>Paired t-test (two tailed) | D2R-IL3 = 8 | AUC | [-0.9 s ; 0 s] Early<br>t=4.036, df=7, p=0.0050<br><br>[-0.9 s ; 0 s] Late<br>t=3.109, df=7, p=0.0171<br><br>[-0.9 s ; 0 s] Early vs. Late<br>t=2.375, df=7, p=0.0493 |
| Suppl. Fig. 4.P | One sample t-test | D2R-IL3 = 8 | AUC | [0.1 s ; 1 s] Early<br>t=3.182, df=7, p=0.0155 |

| | | | | |
|---|---|---|---|---|
| | One sample t-test<br><br>Paired t-test (two tailed) | | | [0.1 s ; 1 s] Late<br>t=1.886, df=7, p=0.1013<br><br>[0.1 s ; 1 s] Early vs. Late<br>t=1.831, df=7, p=0.1097 |
| Suppl. Fig. 4.R | One sample t-test<br><br>One sample t-test<br><br>Paired t-test (two tailed) | GluN1-Δ = 5 | AUC | [-0.9 s ; 0 s] Early<br>t=6.058, df=4, p=0.0037<br><br>[-0.9 s ; 0 s] Late<br>t=2.220, df=4, p=0.0906<br><br>[-0.9 s ; 0 s] Early vs. Late<br>t=0.001277, df=4, p=0.9990 |
| Suppl. Fig. 4.R | One sample t-test<br><br>One sample t-test<br><br>Paired t-test (two tailed) | GluN1-Δ = 5 | AUC | [0.1 s ; 1 s] Early<br>t=4.131, df=4, p=0.0145<br><br>[0.1 s ; 1 s] Late<br>t=2.271, df=4, p=0.0856<br><br>[0.1 s ; 1 s] Early vs. Late<br>t=4.427, df=4, p=0.0115 |
| Suppl. Fig. 4.T | One sample t-test<br><br>One sample t-test<br><br>Paired t-test (two tailed) | GluN1-C1 = 5 | AUC | [-0.9 s ; 0 s] Early<br>t=5.984, df=4, p=0.0039<br><br>[-0.9 s ; 0 s] Late<br>t=2.012, df=4, p=0.1145<br><br>[-0.9 s ; 0 s] Early vs. Late<br>t=2.728, df=4, p=0.0526 |
| Suppl. Fig. 4.T | One sample t-test<br><br>One sample t-test<br><br>Paired t-test (two tailed) | GluN1-C1 = 5 | AUC | [0.1 s ; 1 s] Early<br>t=4.676, df=4, p=0.0095<br><br>[0.1 s ; 1 s] Late<br>t=2.359, df=4, p=0.0778<br><br>[0.1 s ; 1 s] Early vs. Late<br>t=6.285, df=4, p=0.0033 |
| Suppl. Fig. 4.V | One sample t-test<br><br>One sample t-test<br><br>Paired t-test (two tailed) | D2R-Scr = 6 | AUC | [-0.9 s ; 0 s] Early<br>t=5.316, df=5, p=0.0032<br><br>[-0.9 s ; 0 s] Late<br>t=2.520, df=5, p=0.0532<br><br>[-0.9 s ; 0 s] Early vs. Late<br>t=3.525, df=5, p=0.0168 |
| Suppl. Fig. 4.V | One sample t-test<br><br>One sample t-test<br><br>Paired t-test (two tailed) | D2R-Scr = 6 | AUC | [0.1 s ; 1 s] Early<br>t=7.615, df=5, p=0.0006<br><br>[0.1 s ; 1 s] Late<br>t=1.987, df=5, p=0.1036<br><br>[0.1 s ; 1 s] Early vs. Late<br>t=6.352, df=5, p=0.0014 |
| Suppl. Fig. 4.X | One sample t-test<br><br>One sample t-test | D2R-IL3 = 8 | AUC | [-0.9 s ; 0 s] Early<br>t=3.043, df=7, p=0.0188<br><br>[-0.9 s ; 0 s] Late<br>t=1.957, df=7, p=0.912 |

| | | | | |
|---|---|---|---|---|
| | Paired t-test (two tailed) | | | [-0.9 s ; 0 s] Early vs. Late<br>t=2.067, df=7, p=0.0775 |
| Suppl. Fig. 4.X | One sample t-test<br><br>One sample t-test<br><br>Paired t-test (two tailed)t-test (two tailed) | D2R-IL3 = 8 | AUC | [0.1 s ; 1 s] Early<br>t=2.791, df=7, p=0.0269<br><br>[0.1 s ; 1 s] Late<br>t=1.234, df=7, p=0.2572<br><br>[0.1 s ; 1 s] Early vs. Late<br>t=2.626, df=7, p=0.0341 |
| Supplementary Figure 5 – CPT Peptide mPFC | | | | |
| Suppl. Fig. 5.A | One-way ANOVA | Control = 30<br>GluN1-C1 = 21<br>D23-IL3 = 21 | Food Intake | F (2, 69) = 1.218, p=0.3022 |
| Suppl. Fig. 5.B | Two-way RM ANOVA | Control = 30<br>GluN1-C1 = 21<br>D23-IL3 = 21 | Food Intake | Devaluation Effect F(1, 69)=438.0, p<0.0001<br>Group Effect F(2, 69)=0.09665, p=0.9080<br>Devaluation x Group F(2, 69)=0.01484, p=0.9853 |
| Suppl. Fig. 5.C | Kruskal-Wallis test | Control = 28<br>GluN1-C1 = 21<br>D23-IL3 = 20 | Food Intake | H = 2.438, df=2, p=0.2955 |
| Suppl. Fig. 5.D | Two-way RM ANOVA | Control = 28<br>GluN1-C1 = 21<br>D23-IL3 = 20 | Food Intake | Devaluation Effect F(1, 66)=82.11, p<0.0001<br>Group Effect F(2, 66)=0.06859, p=0.9338<br>Devaluation x Group F(2, 66)=0.2150, p=0.8071 |
